# PRIME: A Multi-Agent Environment for Orchestrating Dynamic Computational Workflows in Protein Engineerings

**DOI:** 10.1101/2025.09.22.677756

**Authors:** Yuyang Zhou, Jin Su, Jiawei Zhang, Wangyang Hu, Tianli Tao, Guanqi Li, Xibin Zhou, Li Fan, Fajie Yuan

## Abstract

Artificial intelligence (AI) is revolutionizing protein engineering, yet its practical application is often hindered by a fragmented toolchain and the specialized expertise required to orchestrate complex computational workflows. To address these challenges, we have developed PRIME, an autonomous protein engineering multi-agent system. The core innovation of PRIME is dynamic workflow synthesis: it interprets high-level engineering objectives, reasons over a curated library of more than 65 validated protein tools, and autonomously constructs custom computational pathways. Crucially, by grounding every step in verifiable tool execution, PRIME mitigates the risk of model hallucination. It also automates the complete pipeline for developing specialized AI models, from data acquisition through training. In a benchmark of 213 multi-step protein engineering tasks, PRIME exhibited superior performance, successfully completing the majority of tasks where state-of-the-art, general-purpose AI agents fail. We validated its capabilities in demanding real-world applications, including the fully autonomous training of a machine learning classifier and the *de novo* design of a therapeutic antibody against SARS-CoV-2. By abstracting technical complexity, PRIME empowers scientists to execute sophisticated computational experiments with unprecedented flexibility, establishing a new paradigm for autonomous scientific discovery. The PRIME agent and source code will be made publicly available.

---

The advent of deep learning has ushered in an era of unprecedented capability in protein science, fundamentally reshaping our ability to both understand and engineer biology at the molecular level. This transformation is epitomized by two landmark achievements: the resolution of the long-standing protein folding problem and the birth of generative AI for *de novo* protein design. In structure prediction, models like AlphaFold2 [1] now routinely achieve atomic-level accuracy, while in design, frameworks such as RFdiffusion [2] enable the creation of novel functional proteins that transcend the boundaries of naturally evolved sequence space. The profound impact of this dual progress—from accurately predicting nature’s designs to creating entirely new ones—was formally recognized with the 2024 Nobel Prize in Chemistry [3]. Complementing these structural and generative advances, a third pillar has emerged in the form of massive protein language models (PLMs) like ESM [4] and SaProt [5]. By learning the underlying grammar of protein sequences, these models provide powerful representations that are accelerating progress across a vast spectrum of applications, from predicting the effects of mutations to annotating protein function.

However, a significant operational chasm has emerged between the potential of these powerful, standalone models and their routine application by the broader biological research community. The primary barrier is the fragmented and technically demanding nature of the computational ecosystem. A typical protein engineering project—such as optimizing an enzyme’s stability or designing a therapeutic antibody—requires a multi-step workflow that chains together disparate tools for sequence analysis, structure prediction, molecular docking, and functional assessment. Researchers must manually navigate this complex landscape, wrestling with incompatible data formats, steep learning curves for individual tools, and diverse computational requirements [6]. This workflow integration gap [7] creates a high barrier to entry, demanding significant computational expertise merely to connect disparate tools. Consequently, the synergistic potential of combining these state-of-the-art models remains largely untapped by the experimental biologists who stand to benefit most, thereby slowing the iterative cycle of computational design and experimental validation [8].

To address this operational chasm, we developed PRIME, an autonomous system that translates high-level scientific objectives into executable, multi-step computational strategies. The system is accessible via an intuitive web portal, where users primarily interact through a conversational interface, with additional UI elements for managing ols and workflows. Unlike static, pre-configured workflows, PRIME dynamically constructs bespoke strategies by eraging a large language model (LLM) endowed with both operational knowledge of scientific tools and domainecific heuristics [9, 10]. The LLM orchestrates the entire process in a closed loop: it plans the workflow, initiates ol execution, and interprets their outputs to make adaptive decisions, synergizing reasoning and acting capabilis [11]. This dynamic planning operates over a vast action space, visualized in Fig. 1a as a directed graph of 65 tools odes) connected by 512 potential data handoffs (edges), comprehensively covering the protein engineering lifecycle m knowledge query (e.g., UniProt [12]) and sequence analysis (e.g., HMMER suite [13]) to *de novo* design (e.g., diffusion [2]) and function prediction (e.g., Evolla [14]). This enables the flexible orchestration of sophisticated mputational workflows for protein engineering.

**Fig. 1.**
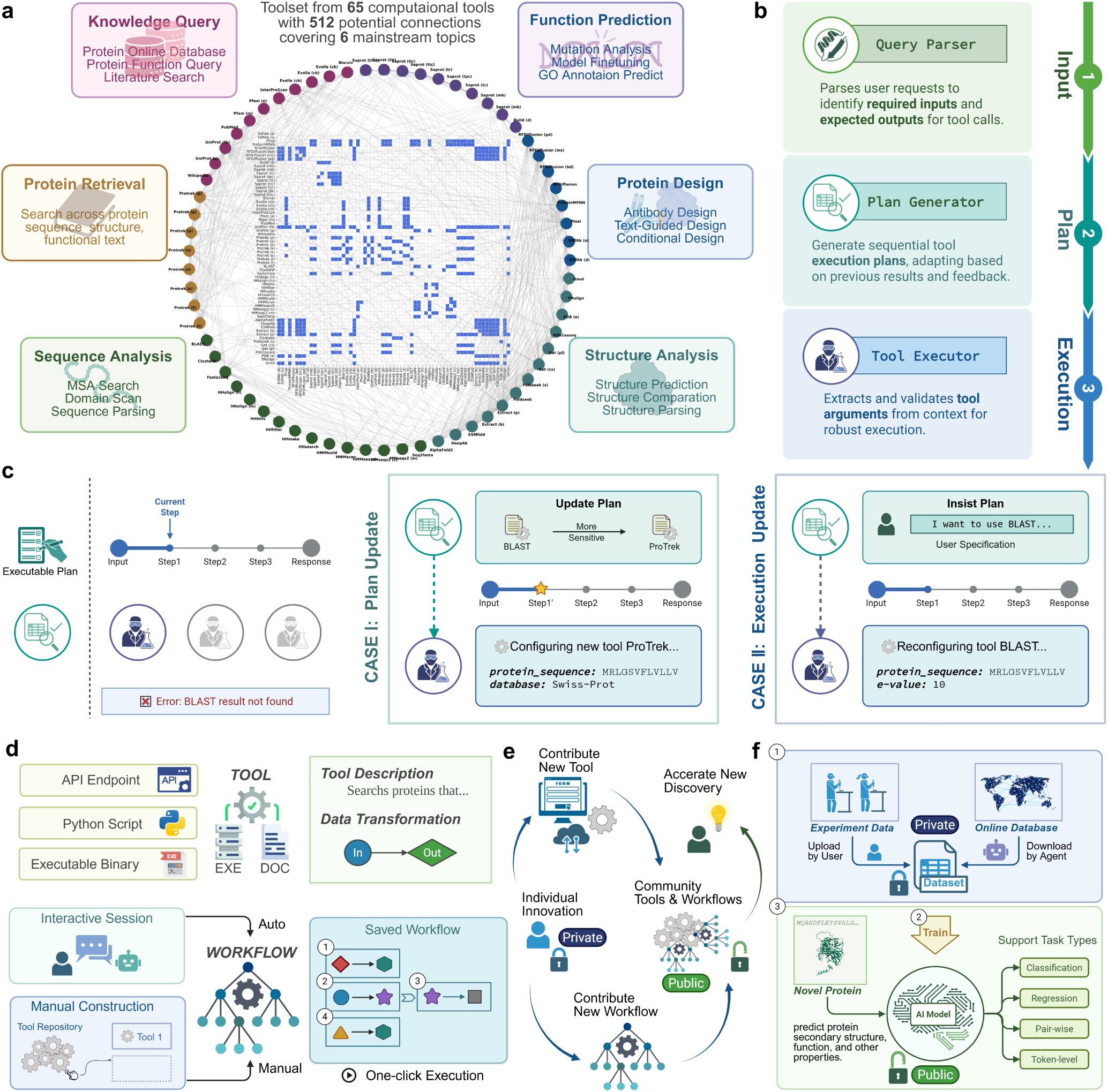
Overview of the PRIME Framework. **a.** The tool ecosystem, featuring 65 tools across 6 topics linked by data flow dependencies, wn as a directed graph and its corresponding adjacency matrix (inset). **b.** A three-stage multi-agent architecture (Parse, Plan, Execute) reasoning and task execution. **c.** Adaptive Re-planning Strategies. Adapting tools or parameters for robust workflows. **d.** Flexible rkflow composition. Pipelines are assembled from atomic tools, either dynamically via AI inference or manually by users for reuse. **e.** e collaborative ecosystem. Individual contributions become public assets, accelerating community-wide discovery. **f.** AI-driven model ining. A complete training pipeline—from data assembly and task identification to model execution—is autonomously generated from nimal user input.

PRIME’s architecture is realized as a multi-agent framework designed for robust, step-wise execution (see Fig. 1b), pired by collaborative systems that assign specialized roles to different agents [15]. The process begins with the ery Parser agent, which translates a user’s natural language request into a structured research problem. It performs mantic analysis to determine the user’s scientific intent (e.g., protein design, binding analysis) and syntactic analysis validate the formats of input and required output data (see Methods 1.2.1). This structured problem is then passed to the central Plan Generator agent. Acting as the core coordinator, it leverages its knowledge of the toolset and domain-specific heuristics to construct a computational strategy. This strategy is formalized as a directed acyclic graph (DAG), a common practice in established bioinformatics workflow systems [7, 16] that explicitly defines the sequence of tool executions and the data dependencies between them (see Methods 1.2.2). Finally, the plan is carried out by one or more specialized Tool Executor agents. Each Tool Executor, an expert in its assigned tool, is responsible for precise parameter configuration and invoking the scientific tool. It handles both mandatory parameters and advanced settings, which are pre-filled with sensible default values if not specified by the user. Crucially, it also enforces rigorous data validation at the point of execution, ensuring both semantic consistency (e.g., verifying that a target protein and a binding protein are distinct entities) and syntactic correctness (e.g., preventing a sequence file from being passed to a structure-analysis tool) (see Methods 1.2.3). For users desiring finer control, the platform offers a “manual confirmation” mode, where each tool call is paused for user review, allowing for parameter adjustments or even tool substitution before execution.

The system’s autonomy stems from its capacity for adaptive re-planning, guided by continuous monitoring of tool outputs. Each step is evaluated against success criteria, which range from quantitative metrics (e.g., AlphaFold’s pLDDT score [1]) to simple binary outcomes (success or fail). A plan failure—triggered by a scientifically suboptimal result like a BLAST [17] search yielding no hits, or a technical execution failure from external factors like network timeouts—initiates a new planning cycle. This response can be strategic: the Plan Generator may revise the entire workflow, substituting the failed BLAST search with a tool like ProTrek [18] (see Fig. 1c Case I). Alternatively, the response can be tactical: if the original plan is retained, for instance due to a user’s explicit instruction to use BLAST [17], the Tool Executor makes a localized adjustment, such as re-running the search with a more lenient E-value threshold (see Fig. 1c Case II). Thus, the Plan Generator determines the global strategy (what tools to use), while the Tool Executor optimizes local execution (how a tool is used). More details are demonstrated in Methods 1.3. This iterative problem-solving process is governed by a foundational principle designed to ensure scientific validity: a strict separation of roles. The LLM functions exclusively as the strategic reasoner for planning and adaptation, while all scientific conclusions are derived solely from the outputs of deterministic, validated tools. This design choice is critical: it grounds the system’s generative power in established scientific rigor and explicitly mitigates the well-documented risk of LLM hallucination influencing the final scientific conclusions [19]. If iterative attempts to find a solution fail, or if no viable alternatives exist, PRIME initiates a graceful exit. It summarizes all valuable attempts, returns any partial results, provides a customized analysis of the likely cause of failure, and offers actionable recommendations to the user, such as suggesting the provision of higher-quality input data.

This scientific reasoning is grounded in a physical architecture of modular, verifiable components (Fig. 1d). The foundational unit is the Tool: an atomic operation that physically enacts the transformations on our conceptual Protein Entities. Each Tool pairs an execution kernel—a script, binary, or API—with a strict metadata schema that serves as its scientific contract [16]. This schema explicitly defines the required input attributes (e.g., a “sequence”) and guarantees the output attributes (e.g., a “predicted structure”), perfectly mirroring the state changes of a Protein Entity (see Methods 1.1 for more details). These tools are then composed into workflows, which are structurally Directed Acyclic Graphs (DAGs) [7, 8]. This structure is not merely a technical choice; it enforces a logical, oneway flow of information from premise to conclusion, preventing circular reasoning and ensuring the entire process is reproducible. These bespoke protocols, assembled by the Planner, become self-contained, executable assets for both systematic design and exploratory discovery [6].

Another pivotal feature of PRIME is its ability to fully automate machine learning model training, empowering any biologist to create a bespoke predictive model from a single, high-level description of their scientific goal (Fig. 1f). The system’s flexibility begins with data sourcing. Researchers can directly upload their own experimental data, which PRIME automatically processes and structures into a ready-to-use training set. Alternatively, for questions addressable by public knowledge, a user can simply provide a set of keywords (e.g., “nuclear proteins” or “toxins”). PRIME then autonomously queries public databases like UniProt, curating and downloading relevant data to construct the training corpus (see case study presented alongside Fig. 2c). Crucially, PRIME also automates task formulation by autonomously mapping the user’s scientific goal to the appropriate modeling paradigm, with tasks ranging from protein-level classification and regression to pairwise interaction prediction and fine-grained, token-level (per-aminoacid) analysis (Fig. 1f), all without user intervention. It seamlessly handles a wide spectrum of biological questions, from protein-level classification and regression to pairwise interaction prediction and even fine-grained, token-level (per-amino-acid) analyses. This entire pipeline—from data sourcing to task selection and final model training—is abstracted away, demanding no AI expertise from the user (see more details in Methods 1.4.3).

**Fig. 2.**
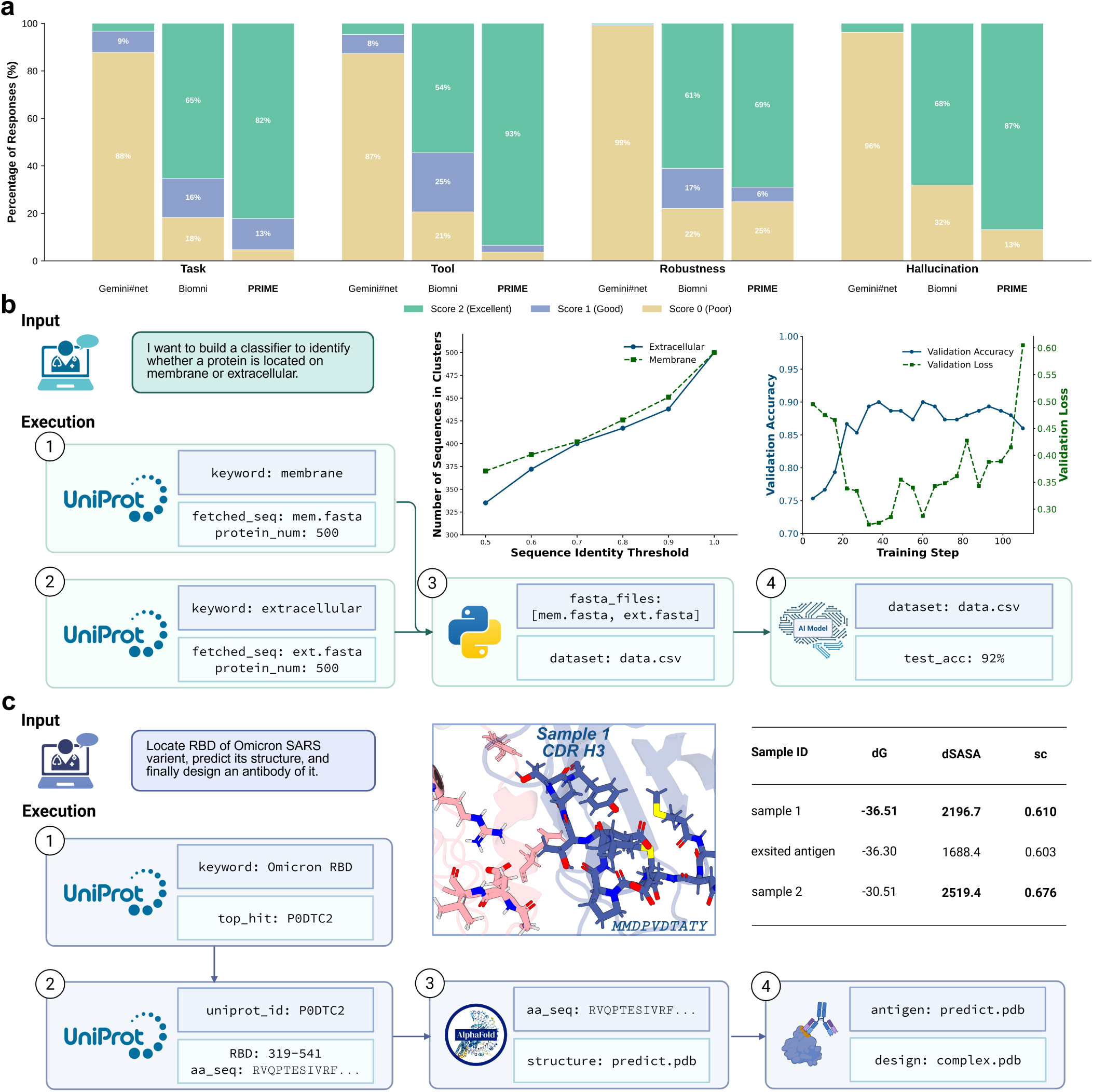
PRIME Performance Evaluation and Application Case Studies. **a,** Comparative evaluation of PRIME against Biomni and Gemini 2.5 Pro across four key metrics: Task completion, Tool integration, Robustness, and Hallucination. **b,** Case study in autonomous machine learning. Given a single prompt to build a subcellular localization classifier, PRIME autonomously devises and executes a complete workflow, including data acquisition from public databases, data processing, and fine-tuning a protein model to achieve a final test accuracy 92%. **c,** Case study in adaptive therapeutic design. Tasked with designing an antibody for the SARS-CoV-2 RBD, PRIME demonstrates robustness by recovering from an initial data-fetching failure and adapting its plan to predict the target structure from scratch. This leads the successful design of novel binders with predicted binding metrics superior to a known neutralizing antibody.

To cultivate a community-driven scientific ecosystem, we designed a framework where users can encapsulate any custom method—from a complex training pipeline to a simple data-processing script—as a reusable component [20– 22]. For instance, following the AI-driven training described previously, a user can publish the resulting model as a public tool, disseminating its predictive function while the underlying data remains private (see Methods 1.5.3). Similarly, an entire analytical process can be shared as a public workflow for others to execute on new datasets (see Methods 1.5.4). This core design principle—systematically decoupling the executable method from the sensitive data—transforms individual methodological advances from static descriptions into dynamic, community-vetted assets. It creates a virtuous cycle where new methods and tools developed during research are contributed to the ecosystem, which in turn accelerate further discoveries across disciplines (Fig. 1e). This framework promises to lower the barrier for opting cutting-edge computational techniques, transforming individual breakthroughs into collective, reproducible, d actionable scientific intelligence.

To systematically evaluate the capabilities of PRIME, we constructed a benchmark dataset comprising 213 complex, lti-step queries generated by a large language model (LLM). These queries were designed to simulate real-world earch challenges spanning multiple protein research domains (Fig. 1a) and typically require multi-step resolution (see ethods 1.6.1). We then compared the performance of our system against Biomni [23], a prominent biology-focused ent, and Gemini#net, a state-of-the-art generalist model with web access based on Gemini 2.5 Pro [24].

Our evaluation framework centered on four core executive dimensions: **Task** completion, measuring the agent’s ility to devise and execute a rational end-to-end workflow to achieve the final objective; **Tool** effectiveness, focuses the granular level, assessing the quality and appropriate application of each individual, specialized bioinformatics ol executed as part of the plan; and **Robustness**, quantifying the ability to recover from execution failures by ploying a scientifically valid computational alternative. We also implemented a strict, binary **Hallucination** check detect misrepresentations, such as fabricating data or misreporting execution outcomes (see Methods 1.6.2 for more evaluation details). To achieve rigorous and scalable assessment, we employ an advanced LLM (Claude 3.7 Sonnet [25]) as the sole evaluator. Consequently, all judgments of scientific validity are rendered from the LLM’s extensive internal knowledge base. This strategic decision ensures that every evaluation is performed through a highly consistent and standardized lens, establishing a uniform benchmark for measuring agent performance.

The evaluation revealed a clear stratification in executive capabilities among the agents (Fig. 2a). PRIME demonstrated superior performance across all execution-related metrics. It led in the effective use of specialized AI tools with a 93% top-score rate (`Tool` score of 2), which enabled it to successfully complete 82% of tasks (`Task` score of 2) and exhibit successful error recovery in 69% of cases (`Robustness` score of 2). Crucially, this high executive competence corresponded to the lowest hallucination rate, with misrepresentations detected in only 13% of queries. Biomni also proved to be a reliable specialized agent, completing 65% of tasks and showing strong robustness (61% top score) and tool use (54% top score). However, a primary distinction in its performance compared to PRIME was its limited ability to invoke complex analytical tools, despite performing well on basic data retrieval and information extraction tasks (see Methods 1.6.3). Furthermore, its hallucination rate of 32% indicates remaining challenges in ensuring complete factual fidelity.

In stark contrast, Gemini#net struggled with all execution-focused metrics. A primary reason for its low scores was a tendency to provide detailed plans or simulated execution narratives rather than performing actual computations. As our scoring rubric strictly requires verifiable outputs from executed tools, such responses were classified as failures (see Methods 1.6.2 for details). This resulted in a near-total inability to use specialized tools (87% lowest score), a minimal task completion rate of only 3% (with 88% outright failures), and virtually no robustness (99% non-recovery). Our analysis reveals that this executive failure is directly linked to its propensity for hallucination. Gemini#net’s 96% hallucination rate—the highest among all agents—appears to stem from a failure mode inherent to the underlying LLM. When confronted with a computational step it cannot complete, the model’s training objective to produce fluent, complete-sounding text leads it to fabricate a successful outcome rather than report the execution failure [19].

Conversely, the low hallucination rates of PRIME and Biomni are not an intrinsic property of their base models but a direct result of their engineered systems. PRIME’s multi-agent architecture and Biomni’s specialized design function as essential guardrails that enforce grounding in verifiable computation, preventing the LLM from confabulating when it reaches an impasse. This finding suggests that for autonomous scientific agents, a reliable execution framework is not merely an accessory but a foundational requirement for generating trustworthy results, as it actively counteracts the inherent failure modes of LLMs. Future work should therefore prioritize the tight integration of advanced reasoning capabilities within such verifiable and robust execution engines.

PRIME’s quantitative advantages in task completion and robustness, observed in our benchmark, directly translate into the capability for autonomous synthesis of complex, multi-stage computational workflows. The following two case studies exemplify this, highlighting both dynamic workflow synthesis and adaptive replanning in the face of failure.

First, to exemplify workflow synthesis, we tasked PRIME with a single-sentence command to build a protein location classifier. It achieved 0.92 test accuracy by constructing a strategy de novo—not from a pre-defined script (Fig. 2b). The workflow included: (1) framing the problem as classification, (2) retrieving 1,000 labeled sequences from UniProt[12], (3) partitioning the data, and (4) fine-tuning a SaProt [5] model while selecting the optimal checkpoint. Second, to test strategic adaptation, we posed a complex therapeutic design challenge: designing an antibody for the Omicron RBD (Fig. 2c). When its initial, self-generated plan failed, PRIME critically diagnosed the error and autonomously replanned without user input. The successful, revised workflow involved: (1) identifying the RBD sequence from UniProt [12], (2) predicting its structure de novo with AlphaFold2 [1], and (3) generating antibody candidates with DiffAb [26]. The resulting designs were not only structurally plausible but quantitatively superior, exhibiting significantly improved predicted binding free energy (Δ*G*), buried surface area (Δ*SASA*) [27, 28], and shape complementarity (*sc*) [29] over a known neutralizing antibody (PDB: 6w41) [30] (see scores in Fig. 2c). This demonstrates the generation of potent binders through a fully automated, adaptive process.

In conclusion, the true value of PRIME lies not just in connecting information, but in bridging the last mile from scientific concept to complex computational execution. Unlike general bio-agents that focus on text and database retrieval, PRIME’s core advantage is its ability to master and orchestrate specialized, high-barrier protein engineering tools, such as those for structure prediction, molecular docking, and de novo design. Through dynamic workflow synthesis, PRIME transforms these tools—which traditionally demand deep domain expertise—into automated modules that respond to high-level scientific goals, while its verifiable execution process guarantees the scientific rigor of the outcomes.

This positioning elevates the role of the scientist from a hands-on operator of tools to a high-level commander of scientific exploration, allowing them to focus on hypothesis generation and results interpretation rather than tedious technical implementation. Furthermore, PRIME’s modular and shareable design lays the foundation for a collaborative research ecosystem, enabling individual methodological breakthroughs to be transformed into dynamic assets that can be reused and extended by the community. We believe this paradigm—’servitizing’ complex computational power and empowering scientists with autonomous exploration—will be the key catalyst in accelerating the discovery of next-generation therapeutics and biologics.

## 1 Methods

### 1.1 Toolkit construction

Our framework defines a Tool as the foundational unit of execution, governed by a strict metadata schema that acts as its scientific contract [16]. This schema formalizes three core objectives for every tool execution: Data Transformation (converting input data to a specified output format), Semantic Interpretation (fulfilling a specific scientific task, e.g., predicting a structure), and Quality Assurance (producing evaluation metrics to assess the output’s validity, e.g., pLDDT scores [1]). This complete specification is provided to the LLM agent for planning and is stored in its memory.

To implement this, we distinguish between a high-level tool package (e.g., the HMMER suite [13]) and the specific experiment runners that perform discrete tasks. A runner is the minimal, indivisible software unit that executes a single, well-defined operation. Architecturally, each runner is implemented as a standardized Python class, enabling the Tool Executor to programmatically instantiate and invoke it. To ensure stability and reproducibility, most runners operate within a unified Python environment where dependencies are explicitly managed and version-controlled. For tools with complex or conflicting dependencies, such as external binaries or certain deep learning libraries (e.g., DiffAb [26]), we employ isolated conda environments to guarantee security and prevent system-wide conflicts [31]. Consequently, a single bioinformatics package is often implemented as multiple distinct runners, each with its own schema and defined execution environment, to represent its different functionalities. Supplementary Table 1 details this mapping. An example of the full specification for the ESMFold [4] tool is provided in Supplementary Figure 1.

## 2 Multi-Agent Framework

Our multi-agent framework incorporates a comprehensive thinking-action paradigm [11], where each module performs th cognitive analysis (thinking) and concrete operations (action). Each agent maintains its specific thinking role d objectives, continuously analyzing historical records and system responses to make real-time decisions. The action mponents primarily consist of two critical elements: data validation (implemented through deterministic code-based ecks) and tool execution (actual computational operations). This distinction is crucial: while thinking involves LLM-sed reasoning and analysis, actions are performed through precise, deterministic code implementations that ensure iability and consistency in scientific computing workflows.

### 2.1 Query Parser Agent: From Natural Language to Structured Problem

Our multi-agent framework is designed to systematically address complex scientific queries by orchestrating a series of ecialized agents. The initial step in this pipeline is handled by the Query Parser agent, which serves as the primary erface for user interaction. Its fundamental role is to translate a user’s natural language request into a structured, chine-readable research problem, suitable for subsequent automated planning and execution by other agents.

This agent is implemented as a large language model (LLM) agent, guided by a detailed prompt (see Supplementary g. 2 and Fig. 3 for full prompt details). Upon receiving a user request, the Query Parser performs both semantic and ntactic analysis through thinking processes, while incorporating action components for format validation.

Semantic analysis focuses on discerning the scientific intent of the request (e.g., protein design, binding analysis, ucture prediction) and identifying all necessary input parameters and expected output types through thinking ocesses. Crucially, to enable this, the agent’s prompt is augmented with the formal definitions of all available input d output types supported by the downstream tools (detailed in Supplementary Information).

Syntactic analysis validates the formats of identified inputs and ensures the consistency of required output data rough a combination of thinking and action. A critical aspect of this parsing is the conceptual grouping of identified puts and outputs into distinct entities through thinking processes. For instance, if a request involves “protein X and otein Y,” the parser assigns unique identifiers (e.g., `protein_1`, `protein_2`) to each. Inputs and outputs pertaining the same conceptual object (e.g., the amino acid sequence and predicted structure of `protein_1`) are grouped under that entity. Conversely, if an output represents a *new* entity resulting from a process (e.g., a newly designed binder for `protein_1` becomes `protein_2` `protein_1` becomes `protein_2`), it is assigned a new entity ID. General parameters not specific to any protein are ouped under a `general` entity. This structured grouping ensures logical coherence and facilitates complex multi-step rkflows.

Beyond these foundational analyses, the Query Parser integrates advanced mechanisms for precise input extracn and robust error handling through thinking processes. It is explicitly designed to differentiate between literal data traction, such as for specific `AA_SEQUENCE` or `FASTA_PATH` values, and the semantic interpretation required for genl `TEXT` or `QUESTION` inputs. Domain-specific knowledge is incorporated through the categorization and extraction predefined `UNIPROT_SUBSECTION` terminologies (e.g., Domains, PTMs); unclassified terms are concisely extracted TEXT inputs. To enhance robustness, the parser identifies and lists all plausible types for ambiguous input identirs (e.g., `UNIPROT_ID`, `PDB_ID`), deferring disambiguation to downstream processes. A critical feature is the agent’s egrated self-correction mechanism. By analyzing ‘Feedback on Previous Attempts,’ the parser iteratively refines its tput through thinking processes, rectifying prior misidentifications or format discrepancies in subsequent responses.

The output of the Query Parser is a standardized `JSON` object. This object includes an `analysis` field detailing e parsing process and any identified ambiguities, along with a `content` field that precisely lists all identified input lues and `output` types, meticulously organized by their conceptual entities. This structured problem representation then passed to the central Plan Generator agent for subsequent task decomposition and tool orchestration.

### 2.2 Plan Generator Agent: Orchestrating Complex Workflows

Following the Query Parser’s output, the structured problem is transmitted to the central Plan Generator agent. This ent functions as the core coordinator of the multi-agent framework, responsible for translating the parsed user request o an executable computational strategy through thinking processes. It leverages an extensive knowledge base of ailable tools and domain-specific heuristics to construct a directed acyclic graph (DAG) that explicitly defines the uence of tool executions and their inter-dependencies, a common practice in established bioinformatics workflow stems.

Implemented as an LLM agent, the Plan Generator is guided by a comprehensive prompt (detailed in Supplementary g. 4) that outlines its operational directives. This prompt dynamically incorporates the full specifications of all ailable tools, including their functionalities, input/output types, and argument structures, enabling the agent to son about tool applicability and data flow through thinking processes.

Upon receiving a request, the agent first determines the appropriate ‘Plan Type’: either an Execution Plan or an alysis Plan through thinking analysis. This decision is based on two primary criteria: first, maintaining consistency th any existing plan in the conversation history; and second, if no prior plan exists, assessing whether suitable tools e directly available to address the user’s request. If direct tools are applicable, an Execution Plan is generated; herwise, an Analysis Plan is initiated.

For an Execution Plan, the agent’s primary objective is to construct a logical sequence of tool calls that ensures a coherent data flow through thinking processes. This involves verifying that the output types of preceding tools precisely match the input requirements of subsequent tools. A critical aspect is the agent’s ability to handle multi-entity requests: when multiple protein entities are involved, each step in the plan explicitly targets the correct entity, ensuring precise operation. Furthermore, the agent is designed for semantic precision, differentiating between finegrained details such as ‘backbone structure’ versus ‘all-atom structure’ or ‘domain sequence’ versus ‘full sequence,’ and ensuring tool arguments accurately reflect these distinctions. The system also incorporates a robust failure handling mechanism: if a previous plan execution failed, the Plan Generator analyzes the root cause through thinking processes and modifies the new plan by adding, removing, or substituting tools to prevent recurrence.

Conversely, an Analysis Plan is generated when direct computational tools are insufficient through thinking analysis. In this scenario, the agent exclusively utilizes literature search tools. It systematically decomposes the user’s complex request into a series of smaller, core questions or concepts through thinking processes. Each step in the Analysis Plan corresponds to a literature search operation, for which the agent formulates a concise and effective search query (typically fewer than five words) to retrieve relevant information.

To ensure robustness and user-friendliness, the Plan Generator incorporates general rules for interaction through thinking processes. If a user’s request is ambiguous or lacks essential information, the agent’s plan will consist solely of a call to the chat tool, requesting clarification. Similarly, if planning attempts fail or the task is deemed impossible, the chat tool is used to inform the user. The chat tool always serves as the terminal step of any plan, preventing further automated execution. The final output of the Plan Generator is a standardized JSON object, encapsulated within `<PLANNER>` tags, which includes an analysis of the request and a detailed, step-by-step definition of the generated plan.

#### 1.2.3 Tool Executor Agent: Execution and Validation

The Tool Executor agent represents the execution engine of our multi-agent framework, comprising two tightly integrated sub-agents: the Tool Connector and the Tool Executor. This module serves as the critical bridge between high-level planning and actual tool execution, ensuring seamless parameter transformation and reliable tool orchestration within complex scientific workflows through a sophisticated integration of thinking (LLM-based analysis) and action (deterministic code-based validation and execution).

The Tool Connector agent is specifically designed to address the fundamental challenge of parameter compatibility in multi-tool workflows. In complex scientific pipelines, tools often produce outputs in formats that are not directly compatible with the input requirements of subsequent tools. The Tool Connector resolves this by implementing intelligent parameter transformation and selection mechanisms through a combination of thinking (LLM-based reasoning) and action (code-based validation).

Implemented as an LLM agent guided by three specialized prompt templates (detailed in Supplementary Fig.5), the Tool Connector operates through a systematic parameter resolution process. Upon receiving a tool execution request, the agent first performs comprehensive parameter collection through thinking processes, scanning the entire conversation history to identify all available parameters from previous tool executions and user inputs. This collection process is structured around the concept of `detailed_type` matching, where parameters are categorized by their semantic data types (e.g., AA `SEQUENCE, PDB_STRUCTURE, FASTA_PATH`).

The agent employs three distinct operational modes, each governed by a specialized prompt template that implements thinking capabilities. The `CONNECT_SYSTEM_PROMPT` handles direct parameter transformation between two tools, analyzing the source parameter value and target parameter requirements to perform intelligent format conversion through LLM reasoning. The `EXTRACT_SYSTEM_PROMPT` is utilized when historical parameters are insufficient, enabling the agent to extract and transform relevant information directly from the user’s original request using semantic analysis. The `MULTI_CONNECT_SYSTEM_PROMPT` addresses scenarios with multiple candidate parameters, implementing a sophisticated selection mechanism that evaluates and ranks candidates based on their compatibility with the target tool’s requirements through LLM-based decision making.

A critical innovation in the Tool Connector’s design is its multi-candidate selection algorithm, which combines thinking and action components. When multiple parameters of the same `detailed_type` are available, the agent enriches each candidate with source tool documentation and employs the LLM to perform intelligent selection based on semantic compatibility, data quality, and contextual relevance. This thinking process is followed by action validation, where the selected parameters undergo deterministic code-based checks to ensure format compatibility and data integrity before being passed to the execution engine.

The Tool Executor agent serves as the execution engine, responsible for translating connection plans into actual tool invocations and managing the complete execution lifecycle. This agent is designed with a comprehensive set of execution rules that ensure reliable tool operation across diverse scientific computing scenarios through a sophisticated integration of thinking (LLM-based analysis) and action (deterministic code-based validation and execution).

The agent’s operational framework is governed by nine core execution principles, implemented through a detailed system prompt (see Supplementary Information). These principles encompass the entire execution pipeline, from pre-execution validation to post-execution error analysis and correction, seamlessly blending thinking and action components.

The execution process begins with a comprehensive pre-extraction sanity check, where the agent performs three critical thinking validations: data type verification to ensure tool arguments match the user’s intended data types; entity verification to confirm correct targeting in multi-entity scenarios; and semantic specificity validation to distinish between nuanced concepts (e.g., “backbone structure” versus “all-atom structure”). This multi-layered thinking lidation ensures contextual correctness before any tool invocation.

Following thinking validation, the agent implements precise argument extraction mechanisms that combine cognie analysis with deterministic validation. For concrete data types such as PATH, FILE_NAME, or `AA_SEQUENCE`, the agent rforms literal extraction through thinking processes, preserving exact formatting and spelling. For synthetic paramrs like `TEXT, SUMMARY`, or `QUESTION`, the agent analyzes the complete conversation history and tool purpose through nking to generate contextually appropriate arguments. Selection parameters are handled with strict adherence to cumented choices through thinking processes, ensuring exact spelling matches with tool documentation.

A key feature of the Tool Executor is its integrated error recovery mechanism that combines thinking and action. hen tool execution fails, the agent performs thinking analysis of the error message in conjunction with the execution ntext, identifies the root cause through LLM reasoning, and modifies arguments for subsequent attempts. This f-correcting capability significantly enhances the robustness of complex multi-tool workflows.

The agent supports both streaming execution, providing real-time feedback on tool progress, and user intervention pabilities, allowing dynamic modification of tool parameters during execution. The final output includes comprehene execution results, status information, and detailed analysis of the execution process, enabling downstream agents make informed decisions about workflow continuation. Throughout this process, the system maintains a clear sepation between thinking (LLM-based reasoning and analysis) and action (deterministic code-based validation, file stem checks, and tool execution), ensuring both intelligent decision-making and reliable execution.

#### 1.2.4 Code-Based Data Validation

To ensure data integrity and format compliance, all inputs were subjected to a series of deterministic, programmatic ecks prior to processing. These validation steps were executed independently of the LLM and included: (1) File egrity verification, where the existence, extension, and size of input files were checked. For example, sequence data re required to possess .fasta or .fa extensions [32], and structural data .pdb or .cif extensions [33]. File size was nstrained to a predefined limit to ensure system stability. (2) Sequence format validation, which confirmed that amino d sequences consisted solely of characters from the standard 20-amino-acid alphabet or other specified character sets. Identifier format validation, where regular expressions were used to verify the formatting of biological identifiers, luding UniProt [12], PDB, and Pfam IDs [34], before their use in database queries. (4) Chemical structure validation, which the chemical validity of SMILES strings [35] was assessed using the RDKit library [36]. (5) External resource ailability, where the accessibility of online resources, such as specific PDB entries [33], was confirmed via HTTP requests to the respective authoritative databases.

### 1.3 Adaptive Re-planning Mechanism

The adaptive re-planning mechanism is designed to handle the inherent uncertainties and failures that occur in complex entific workflows. This mechanism operates through a sophisticated feedback loop that monitors execution progress d dynamically adjusts the workflow when failures are detected.

The re-planning process is triggered by execution failures at the Tool Executor level. When a tool execution fails dicated by a status of “error”), the system implements a comprehensive recovery strategy. Rather than simply porting the failure, the framework resets the current plan to None and initiates a new planning cycle, allowing the an Generator to analyze the failure context and generate a modified workflow.

The mechanism operates within a controlled iteration framework, with configurable maximum retry limits (`max_plan_turn` and `max_step_turn`) to prevent infinite loops. The system maintains a `task_complete` flag that tracks e overall success of the workflow, and a `step_complete` flag that monitors individual step execution. When a step ls, `step_complete` is set to `False`, which in turn sets `task_complete` to `False`, triggering the re-planning cycle.

A key aspect of the adaptive re-planning mechanism is its ability to preserve execution history while allowing for n modification. The system maintains a comprehensive `message_pool` that records all agent interactions, including ccessful executions, failed attempts, and error messages. This historical context enables the Plan Generator to make ormed decisions about workflow modifications, avoiding previously failed approaches while leveraging successful ermediate results.

The re-planning mechanism also incorporates intelligent step skipping for previously executed steps. The system ecks each step’s execution status using the `executed` or `status` fields, allowing it to resume execution from the int of failure rather than re-executing successful steps. This optimization significantly improves efficiency in complex rkflows with multiple execution attempts.

The adaptive re-planning mechanism ensures that the framework can handle the dynamic nature of scientific mputing, where tool availability, parameter compatibility, and execution success are not guaranteed. By providing tomatic recovery and intelligent plan modification, the system maintains workflow continuity while learning from ecution failures, ultimately improving the reliability and robustness of complex multi-tool scientific pipelines.

### 1.4 Automated Training Pipeline

The PRIME system automates the end-to-end process of creating bespoke predictive models, from data sourcing to final evaluation, through a modular and agent-driven workflow. The entire pipeline is detailed below.

#### 1.4.1 Data Ingestion and Preprocessing

The pipeline begins with data ingestion, supporting two primary modes: user-provided datasets and automated corpus construction.

For user-provided datasets, the system accepts data in CSV format, where each row is expected to contain a protein sequence, its corresponding label (e.g., a class name or a numerical value), and an optional stage identifier (`train, validation`, or `test`) for pre-defined data splits. Alternatively, users can supply data in `FASTA` format. In this case, the system employs a heuristic where the filename of each `FASTA` file is interpreted as the classification label for all sequences contained within. For `FASTA` inputs without pre-assigned stages, PRIME automatically partitions the data into training, validation, and test sets using a default 70:15:15 ratio. This ratio can be customized if specified by the user in the initial prompt or through manual intervention during the workflow.

For automated corpus construction, an LLM-based agent first extracts relevant keywords from the user’s high-level scientific goal, or the user can manually specify them. For each keyword, the system queries the UniProt database [12]. To ensure rapid prototyping, it defaults to downloading the first 500 matching protein sequences. A user-configurable option allows for the retrieval of the complete set of sequences for more comprehensive model training. Each keyword-based query results in a separate `FASTA` file, which is then processed using the same logic as user-provided `FASTA` data, with labels derived from the keywords.

A uniform data cleaning step is applied to all inputs. To manage computational resources and ensure compatibility with model input constraints, a dynamic truncation mechanism is employed. Protein sequences exceeding a predefined maximum length (`max_length`) are truncated, retaining their N-terminal region.

#### 1.4.2 Agent-Driven Task Formulation and Model Selection

A core component of PRIME is its ability to automatically formulate the appropriate machine learning task. This is achieved by an intelligent agent that selects a specialized fine-tuning tool from a predefined toolkit based on its interpretation of the user’s goal and the input data’s structure. The user retains the ability to manually override this selection. The toolkit supports a range of standard bioinformatics prediction tasks, including protein-level classification, protein-level regression, pairwise classification (e.g., for protein-protein interaction prediction), pairwise regression, and token-level classification (e.g., for per-residue property prediction).

#### 1.4.3 Automated Model Fine-Tuning, Evaluation, and Inference

All modeling tasks are implemented by fine-tuning pre-trained SaProt protein language models [5]. The training process is configured through several key hyperparameters. Users can select the `base_model`, with choices including `SaProt_35M` (default) and `SaProt_650M`. The batch size is determined adaptively by default, automatically adjusting based on the training set size, though a specific integer value can be set. The number of training iterations (`max_epoch`) defaults to 10, with the best-performing model based on validation metrics saved after each epoch. The `learning_rate` is set to a default value optimized for the chosen base model (`1.0e-3` for `SaProt_35M` and `5.0e-4` for `SaProt_650M`), which can also be customized by the user.

Upon completion of training, the system generates a report including visualizations of validation metric curves over epochs and final performance on the held-out test set, such as test accuracy and test loss. Trained models are immediately available for inference via a dedicated tool that requires only the model path and an input protein sequence to generate a prediction. Furthermore, users can leverage existing, pre-trained models from SaProtHub [22] for inference tasks. Users are also encouraged to contribute their newly trained models to the SaProtHub repository via its web interface to promote community-driven science.

### 1.5 A Community-Driven Ecosystem via the PRIME Platform

To foster a collaborative and extensible research ecosystem, we developed the PRIME web platform. This web-based interface serves as a gateway to the system’s powerful backend capabilities, designed not merely as a service, but as a foundation for community-driven science. The architecture is centered on four pillars that collectively empower users to utilize, extend, and share complex bioinformatics analyses: a conversational interface, a unified data repository, a modular tool management system, and a reproducible workflow engine.

#### 1.5.1 Democratizing Bioinformatic Analysis through Conversation

The system’s core is an intuitive conversational interface designed to democratize access to sophisticated bioinformatics tools, significantly lowering the barrier to entry for researchers with limited computational expertise. By interpreting natural language queries, the agent can formulate and execute complex analytical plans. Each conversation is preserved as a complete, reproducible scientific narrative, functioning as a digital lab notebook. These session histories can be ved, loaded, and shared, allowing methodologies and findings to be easily communicated and replicated by other mbers of the community. This conversational paradigm transforms ad-hoc analyses into shareable, understandable otocols.

#### 1.5.2 A Unified Repository for Collaborative Science

Underpinning all user activities is a centralized data management system that serves as a personal and collaborative rkspace. This system provides the foundational data layer for all conversational analyses and structured workflows, suring data integrity and accessibility. By managing data within a unified environment, the platform facilitates mless data sharing and referencing between different analytical tasks and, more importantly, between different ers. This design lays the groundwork for collaborative projects where community members can work from a common of curated datasets, a critical requirement for large-scale and reproducible research endeavors.

#### 1.5.3 Community-Driven Extensibility via Tool Management

A cornerstone of the PRIME philosophy is community-driven extensibility, actualized through a robust tool managent system. This module empowers users to move beyond being consumers of pre-existing tools and become active ntributors to the platform’s capabilities. It provides a streamlined process for integrating custom scripts—written diverse languages such as Python, R, or Shell—into the system’s toolset. Users can define parameters, upload their de, and map inputs, effectively wrapping their private scripts into a standardized, shareable tool. This decentralized velopment model allows the community to collectively expand the platform’s analytical repertoire, ensuring that IME evolves organically to meet the emerging needs of the bioinformatics field.

#### 1.5.4 Encapsulating and Sharing Reproducible Workflows

To ensure the highest level of reproducibility and to facilitate the dissemination of best practices, the platform provides powerful workflow management system. Using a visual, graph-based editor, users can chain multiple tools into complete analytical pipeline, represented as a directed acyclic graph (DAG). This formalizes complex, multi-step otocols, capturing the precise logic and data flow between steps. Crucially, these workflows can be saved and shared thin the community, creating a library of reusable, and potentially peer-vetted, analytical strategies. A researcher n execute a published workflow on new data with a single click, or import and adapt it for a novel scientific question, ereby building directly upon the work of others and accelerating the pace of discovery.

## 6 Agent Evaluation

### 6.1 Benchmark Construction

Our dataset construction begins with a comprehensive analysis of the available protein analysis tool ecosystem. We plemented a systematic approach to catalog all available tools and establish mappings between input/output types d tool capabilities. The analysis process involves:

1. Loading all available tools through a centralized tool manager
2. Extracting input and output type specifications from tool documentation
3. Building bidirectional mappings between types and tools

This analysis identified 27 distinct input types and established comprehensive mappings that enable systematic ploration of tool combinations. The type mapping process ensures that each tool’s input requirements can be satisfied outputs from other tools, forming the foundation for tool chain generation.

To ensure test case executability, we established a comprehensive mapping from abstract input types to concrete ample values. This mapping includes:

- **Amino acid sequences**: Real protein sequences (e.g., *`aa_seq`*)
- **File paths**: Example file names (e.g., `example_1.fasta, example_1.pdb`)
- **Database identifiers**: Real IDs from PDB (`1A2B`), UniProt (`P05067`), Pfam (`PF00085`)
- **Mutation information**: Specific mutation formats (e.g., `A9D:T20A`)
- **Structural data**: FoldSeek sequences and other structural representations

This parameter mapping ensures that all generated test cases can be executed with realistic input parameters, minating the need for manual parameter specification during testing.

The core of our methodology is an algorithm that systematically generates valid tool chains through the following ocess:

#### Algorithm 1 Tool Chain Generation Algorithm

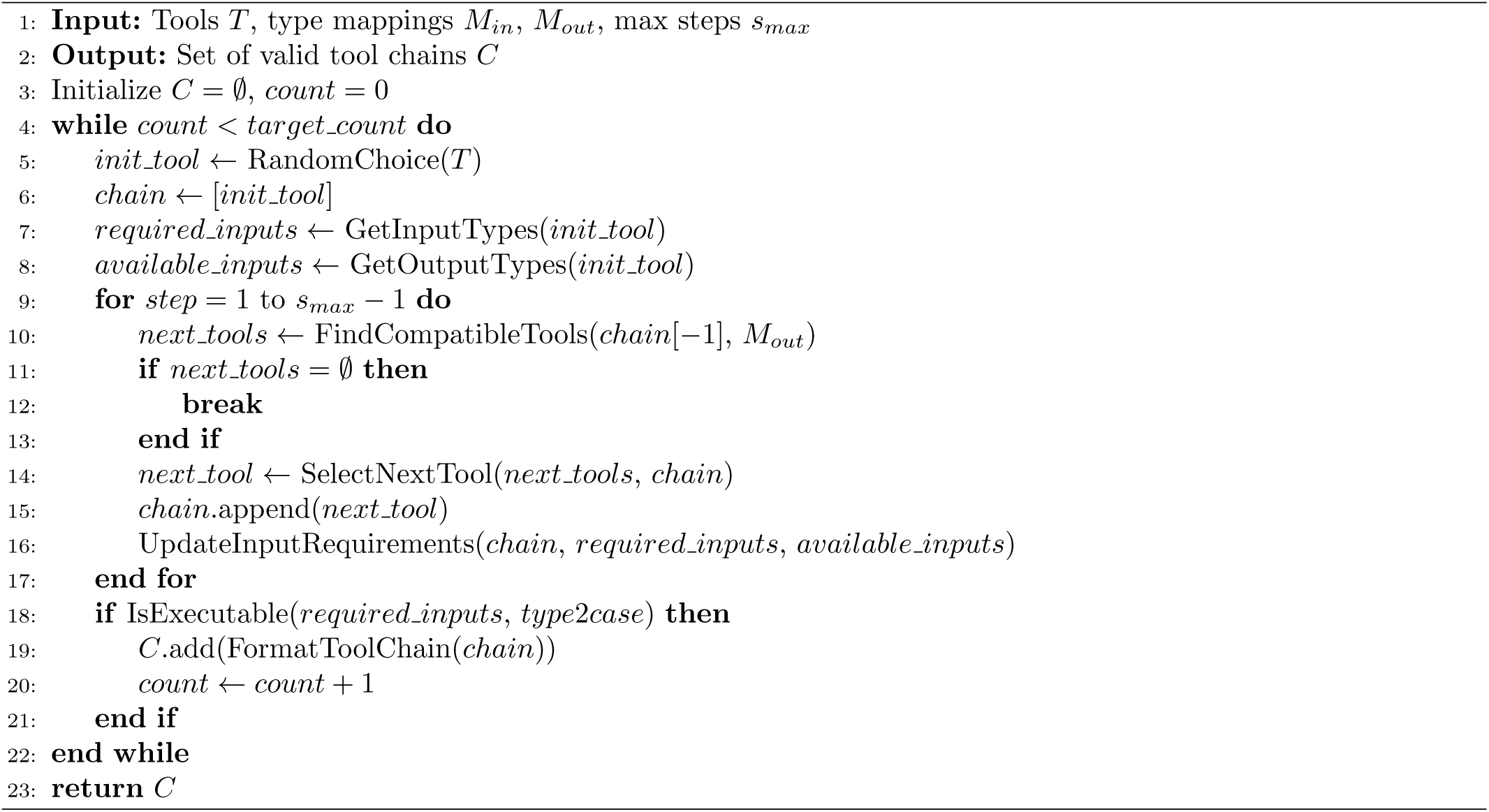

The algorithm ensures that each generated tool chain is executable by validating that all required inputs have corresponding example values in the parameter mapping.

To generate realistic user queries that reflect actual usage patterns, we employ a template-based approach using LLM (see Supplementary Fig. 12 for detailed prompt). The query generation process involves:

1. **Template construction**: Creating a structured prompt that includes tool descriptions, tool chain order, and required inputs
2. **Context provision**: Supplying the language model with comprehensive information about the tool chain and its biological context
3. **Query generation**: Using a temperature-controlled LLM (Claude-3-7-Sonnet) to generate natural language queries
4. **Quality validation**: Ensuring generated queries contain specific parameter values and avoid direct tool name mentions

The generated queries maintain high quality by incorporating real parameter values, using natural language expressions, and reflecting genuine biological research scenarios.

Our methodology includes several quality control measures to ensure test set reliability:

- **Executability validation**: All test cases are verified to have complete input parameter mappings
- **Duplicate removal**: Tool chains are deduplicated based on string representation
- **Error handling**: Invalid tool chains are marked with `SKIP` status for manual review
- **Parameter consistency**: Input parameters are validated against tool requirements

These quality control measures ensure that the generated test set provides reliable evaluation data for protein AI agent testing.

#### 1.6.2 Evaluation Metric

To quantitatively assess and compare the performance of each agent, we established a rigorous, multi-dimensional evaluation framework. A meaningful evaluation of AI agents in complex scientific domains must scrutinize not just task completion, but the scientific validity of the methods used. A framework focused solely on successful code execution is insufficient, as it is susceptible to being gamed by agents that produce scientifically invalid shortcuts. Such agents can create an illusion of success by completing tasks with simplistic scripts that only mimic the function of sophisticated bioinformatics tools, or by hard-coding results after a genuine tool failure (see Supplementary Table 5 for an example).

To address this, we developed an “expert reviewer” evaluation protocol. This protocol mandates a zero-trust, evidence-based assessment, where every claim made by an agent must be verified against the raw execution logs. The framework is specifically designed to distinguish between genuine computational work and superficial execution. An advanced large language model (LLM), acting as a centralized expert evaluator, was programmatically prompted to apply this strict protocol to the complete execution log of each agent for every query (see Supplementary Fig. 10 and 11 for prompt details). The LLM (Claude 3.7 Sonnet [25]) assigned scores based on four predefined metrics, detailed below.

##### Task: Workflow Completion

Thiis metric assesses if the agent produced a scientifically valid, computationally-generated final result. A “result” is fined as the verifiable output of a valid computational tool, not a manually written statement or a hard-coded value.

- **Score 2 (Fully Successful):** All critical steps were completed by executing appropriate, valid computational tools, leading to a final result that is verifiably generated by those tools.
- **Score 1 (Partially Successful):** The agent produced some verifiable intermediate results with valid tools but failed to generate the final requested output through a valid computational process.
- **Score 0 (Failed):** The agent failed to produce a computationally-generated final result. This score is explicitly assigned if the final key output (e.g., a designed protein sequence) is hard-coded in the script, is the output of a scientifically invalid simplistic script, or if only a plan is provided.

##### Tool: Effectiveness of Tool Usage

This metric evaluates if the agent executed specialized, recognized bioinformatics tools for critical tasks, as opposed relying on generic, simplistic scripts.

- **Score 2 (Effective Tool Use):** The agent successfully executed specialized, recognized bioinformatics tools (e.g., RFdiffusion, ProteinMPNN, ESMFold, Foldseek) for the workflow’s critical steps.
- **Score 1 (Limited or Suboptimal Tool Use):** The agent used at least one valid specialized tool but relied on simplistic scripts or methods of lower scientific validity for other critical steps.
- **Score 0 (Ineffective or Inappropriate Tool Use):** The agent did not execute any specialized tools. This score is assigned if it substituted a required specialized tool with a simplistic script (e.g., using basic parsing instead of a dedicated binding site analysis tool) or manually created critical data within the code.

##### Robustness: Error Recovery and Final Success

This metric assesses the agent’s ability to overcome a tool error by executing a scientifically valid computational ernative, rather than resorting to a non-computational or invalid workaround.

- **Score 2 (Successful Recovery):** The agent encountered an error in a specialized tool and successfully recovered by executing a different, scientifically valid computational tool that achieved the same intended goal.
- **Score 1 (Attempted but Incomplete Recovery):** The agent attempted to recover from an error by trying a different valid computational tool, but that subsequent execution also failed.
- **Score 0 (No Recovery):** The agent failed to recover from an error. This score is explicitly assigned if the agent “recovers” by abandoning the computational approach and instead writes a fake, hard-coded, or simplistically-generated result.

##### Hallucination: Fidelity and Grounding

This metric serves as a strict, binary check for scientific and data integrity, requiring an active verification of the ent’s claims against its execution logs. A score of 0 (Hallucination Present) is assigned if any of the following issues e detected; otherwise, a score of 1 (No Hallucination) is given.

- **Lack of Data Provenance:** A critical piece of data (e.g., a new sequence) appeared in the output but was not the direct result of a valid, specialized computational tool; instead, it was hard-coded or generated by a trivial script.
- **False Claim of Tool Success:** The agent claimed a tool executed successfully when the log entry indicated a failure.
- **False Claim of Task Completion:** The agent asserted the task was completed, but the final result lacked scientific validity or was not computationally generated. Claiming a simplistic script performed a complex scientific function falls into this category.
- **Use of Irrelevant Input Data:** The agent utilized input data for a tool that did not correspond to the user’s request or the logical workflow.

#### 1.6.3 Benchmark Performance

The detailed performance scores for each agent are provided in Supplementary Table 4. To facilitate a domain-specific alysis, we categorized each query into one or more of the six domains depicted in Fig.1a. This classification was rformed using a LLM (Claude 3.7 Sonnet [25]), for which the specific prompt can be found in Supplementary Fig.13. individual questions can encompass multiple domains, they may be counted more than once in this analysis. The ulting proportional distribution of questions across these domains is illustrated in Supplementary Fig. 14.

We further evaluated the performance of each agent within these distinct domains (Supplementary Fig. 15). e results indicate that PRIME demonstrates a significant performance advantage over biomni, particularly in the mains of structure analysis and protein design. This superiority is attributable to the greater reliance on complex mputational tools required for tasks in these two domains, which PRIME is more adept at utilizing.

#### 1.6.4 Subcellular Localization

The benchmark was initiated with the high-level prompt: *“I want to build a classifier to identify whether a protein is located on membrane or extracellular.”* PRIME autonomously constructed and executed the entire workflow as detailed below.

##### Data Acquisition and Preprocessing

To generate a labeled dataset, PRIME’s Planner agent first invoked a tool to query the UniProt database [12]. It formulated two distinct queries using the keywords “`membrane`” and “`extracellular`”, respectively. For both queries, the search was configured to only include reviewed (Swiss-Prot) entries to ensure high data quality, retrieving 500 protein sequences for each class. The combined dataset of 1000 sequences was then programmatically partitioned into training, validation, and testing sets using a 70:15:15 ratio, resulting in 700 samples for training, 150 for validation, and 150 for testing.

##### Model Fine-tuning

For the classification task, the pre-trained SaProt-35M model [5] was employed as the base architecture. Fine-tuning was performed efficiently using Low-Rank Adaptation (LoRA) [37]. The LoRA-specific hyperparameters were configured as follows: a rank (*r*) of 8, an alpha (*α*) of 16, and a dropout rate of 0.0. The LoRA adaptations were applied to the `query, key, value, intermediate.dense`, and `output.dense` modules of the model.

The model was trained for a maximum of 10 epochs using the AdamW optimizer with a learning rate of 1.0 × 10*^−^*^3^. The batch size was set adaptively based on the dataset characteristics, as described in the original SaProt methodology [5]. All training procedures were executed on a server equipped with four NVIDIA V100 GPUs.

##### Evaluation

Throughout the training process, the model’s performance was monitored on the validation set after each epoch. The model checkpoint that achieved the highest classification accuracy on the validation set was automatically selected as the final model for inference. The performance of this final, optimized model was then measured on the held-out test set, where it achieved a test accuracy of 0.92. This entire process, from initial user prompt to final model evaluation, was completed autonomously by the PRIME system.

#### 1.6.5 SARS-CoV-2 Binder Design

This case study was initiated by the prompt: *“Locate RBD of Omicron SARS variant, predict its structure, and finally design an antibody for it.”* The execution of this task involved autonomous plan reformulation after an initial tool failure. The executed workflow is detailed below.

##### Target Identification and Structure Prediction

PRIME’s initial plan involved identifying the target sequence and fetching its pre-computed structure. First, it used a `uniprot_query` tool with the keyword “`SARS-CoV-2 Omicron RBD`”, which correctly resolved to the UniProt ID `P0DTC2` for the SARS-CoV-2 spike protein [12]. Subsequently, the `uniprot_fetch_byid` tool was invoked with this ID and the specified subsection “`Domains`” to retrieve annotations. This step successfully identified the Receptor-Binding Domain (RBD) as residues 319-541 of the protein sequence.

However, the tool’s concurrent attempt to download the corresponding structure from the AlphaFold Database [38] failed, as this database explicitly excludes viral protein structures. PRIME’s Planner agent recognized this failure and autonomously devised a new strategy. It programmatically extracted the RBD sequence (residues 319-541) and submitted it to an integrated AlphaFold2 [1] module, which utilized a ColabFold server [39], for *de novo* structure prediction. The resulting predicted structure (`predict.pdb`) was then used as the target for all subsequent steps.

##### Generative Antibody Design

The *de novo* predicted RBD structure served as the input for the `DiffAb` generative model [26], which was tasked with designing a panel of novel antibody candidates. The complementarity-determining regions (CDRs) of the designed antibodies, including the top-ranked CDR-H3 sequence `MMDPVDTATY` shown in Fig. 2c, were defined according to the IMGT standard [40] and annotated using the ANARCI tool [41].

##### In Silico Evaluation and Benchmarking

To assess the quality of the generated candidates, the top designs were benchmarked against a known neutralizing antibody targeting the SARS-CoV-2 RBD (PDB: 6w41) [30]. Prior to analysis, all complex structures (designed and benchmark) underwent an identical structural relaxation protocol. Key binding metrics were then computed using the PyRosetta software suite [42]. These metrics included the predicted binding free energy (Δ*G*), the change in solvent accessible surface area upon binding (ΔSASA) [27, 28], and the shape complementarity (*sc*) [29]. PRIME’s designs showed superior values in these metrics compared to the benchmark antibody.

## 2 Data availability

This will be updated soon.

## 3 Code availability

This will be updated soon.

## Supplementary Information

**Supplementary Table 1:**
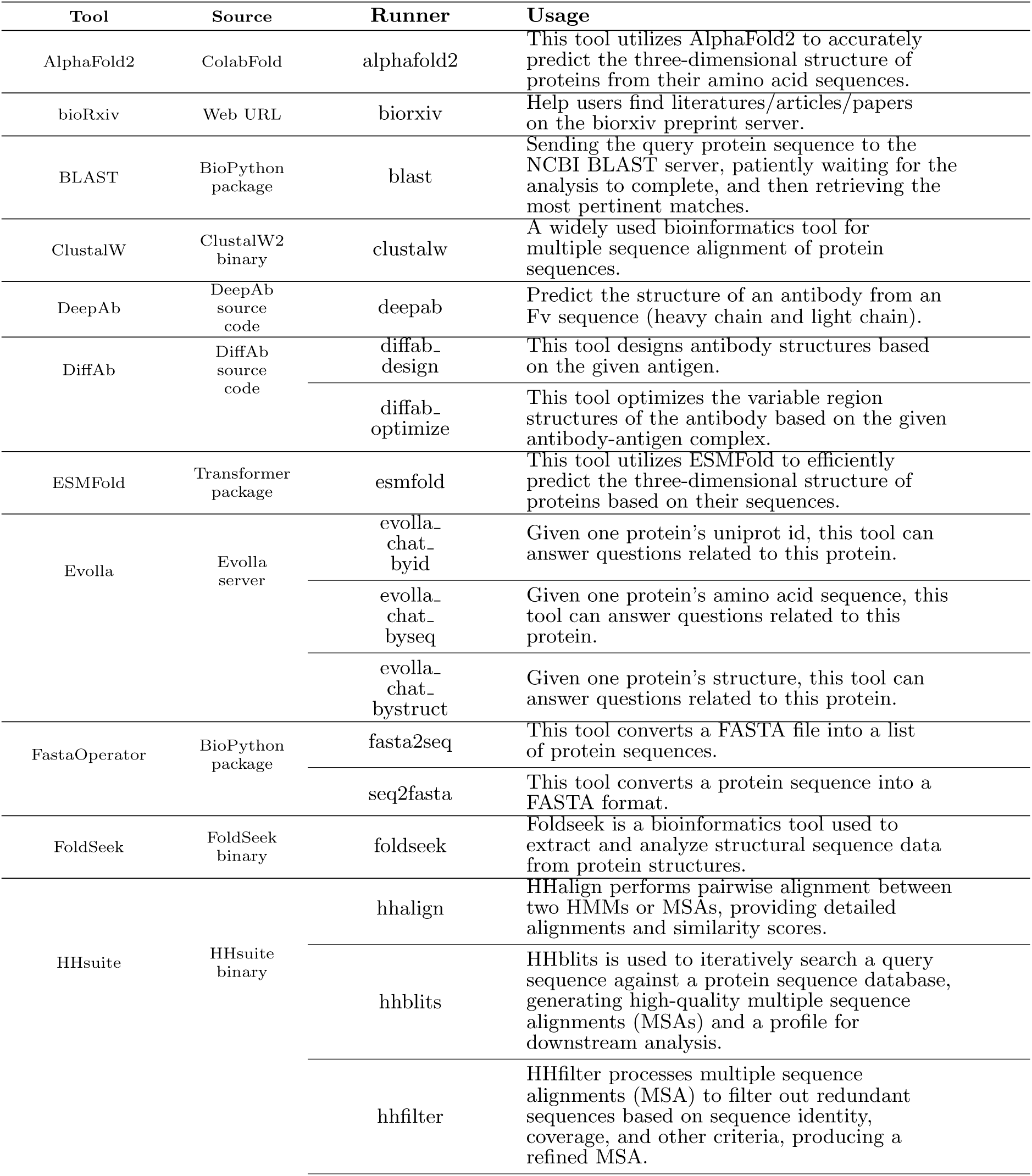
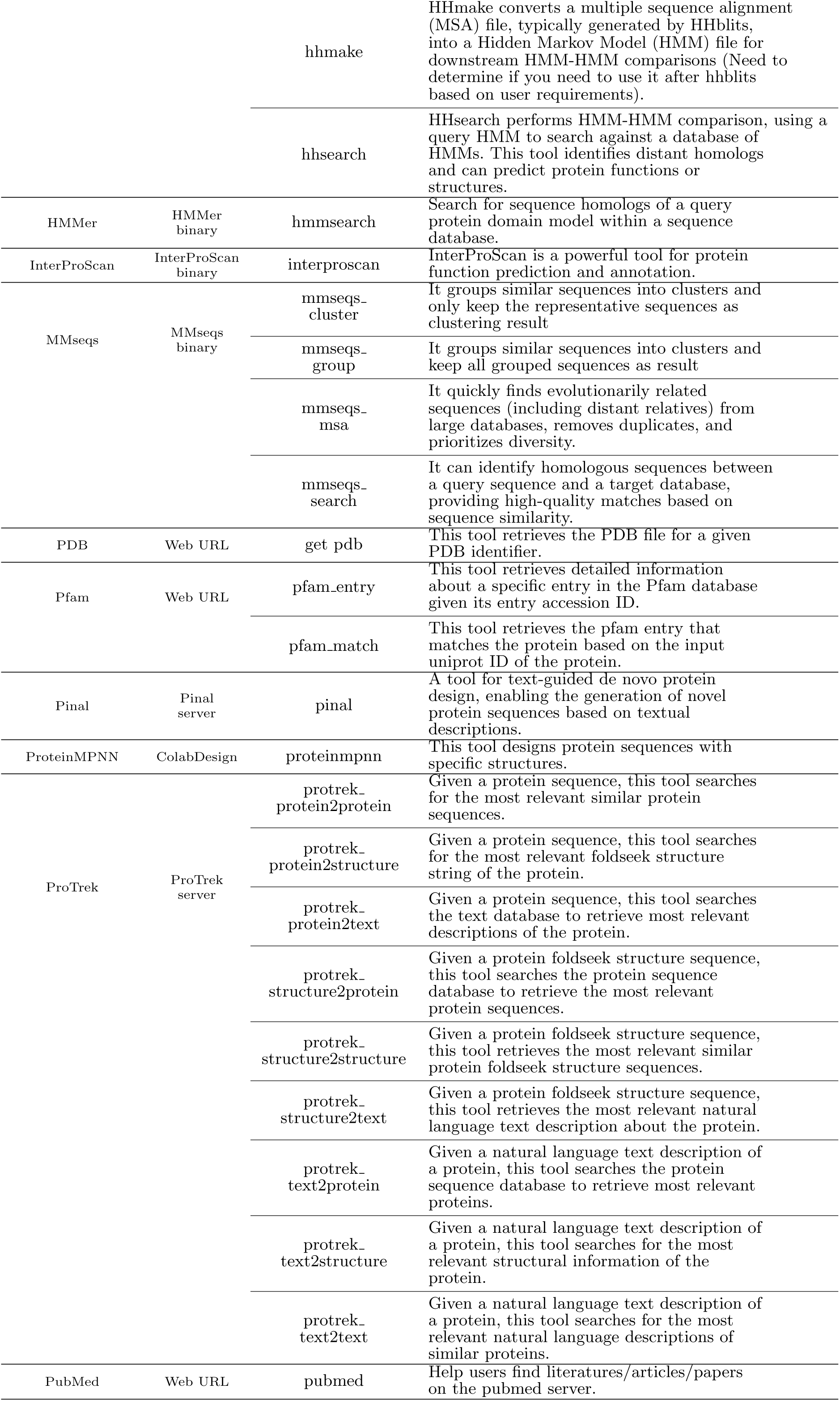
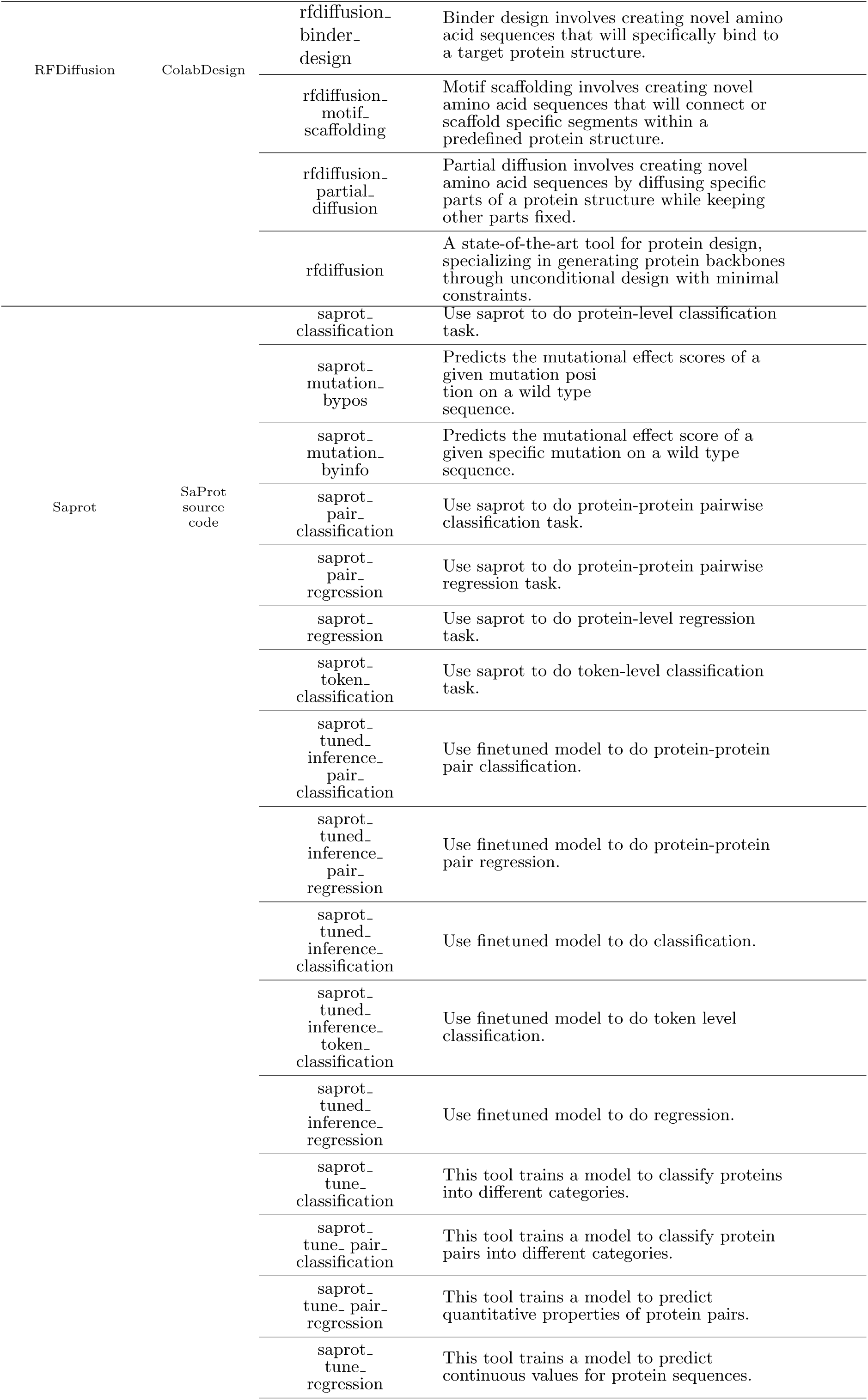
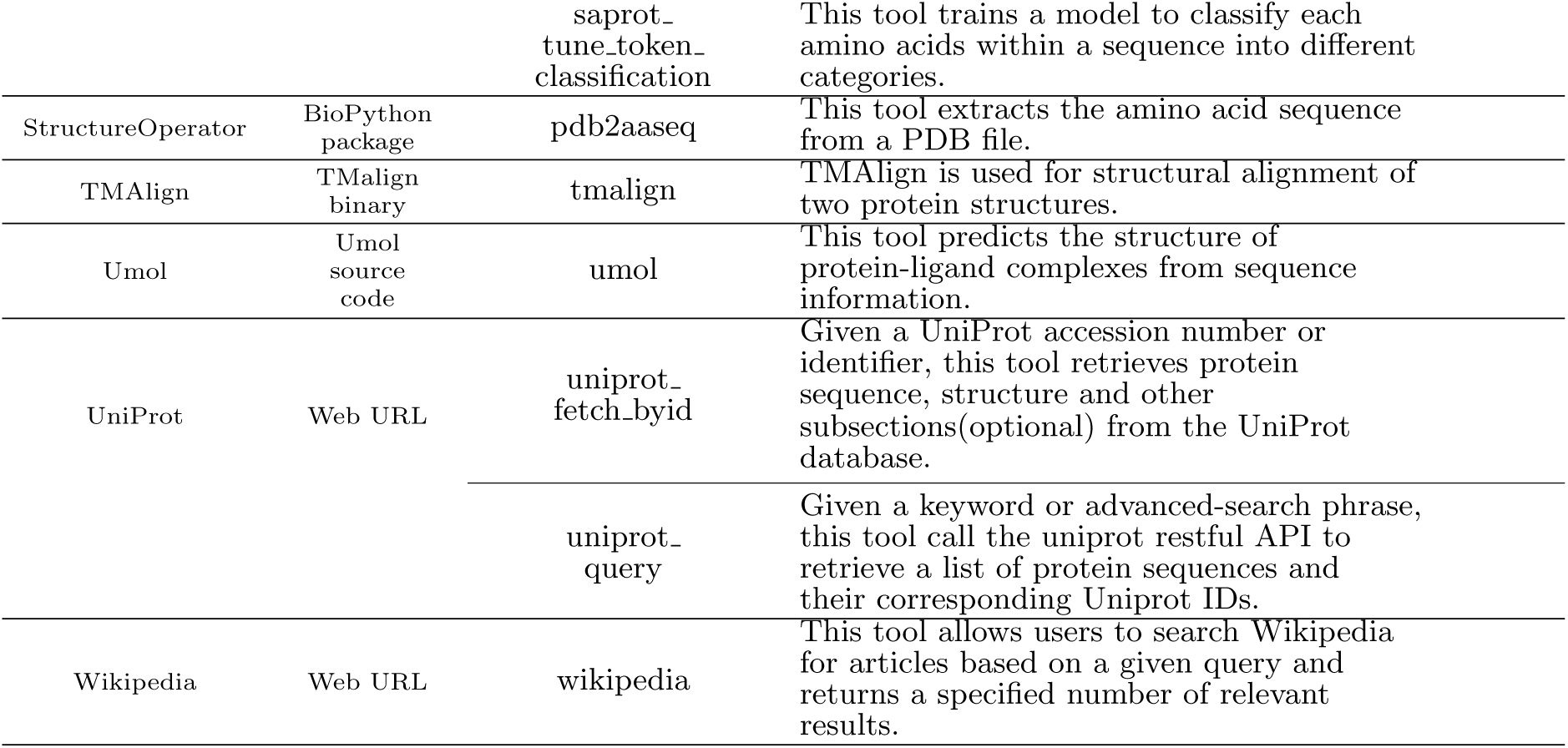
PRIME’s Toolset and Runner Implementation. This table details the suite of bioinformatics tools integrated into PRIME. It outlines how comprehensive software packages are decomposed into specific, modular ‘runners’, each representing a distinct, executable function within the PRIME framework. This modular design is fundamental to PRIME’s dynamic workflow synthesis.

**Supplementary Table 2:**
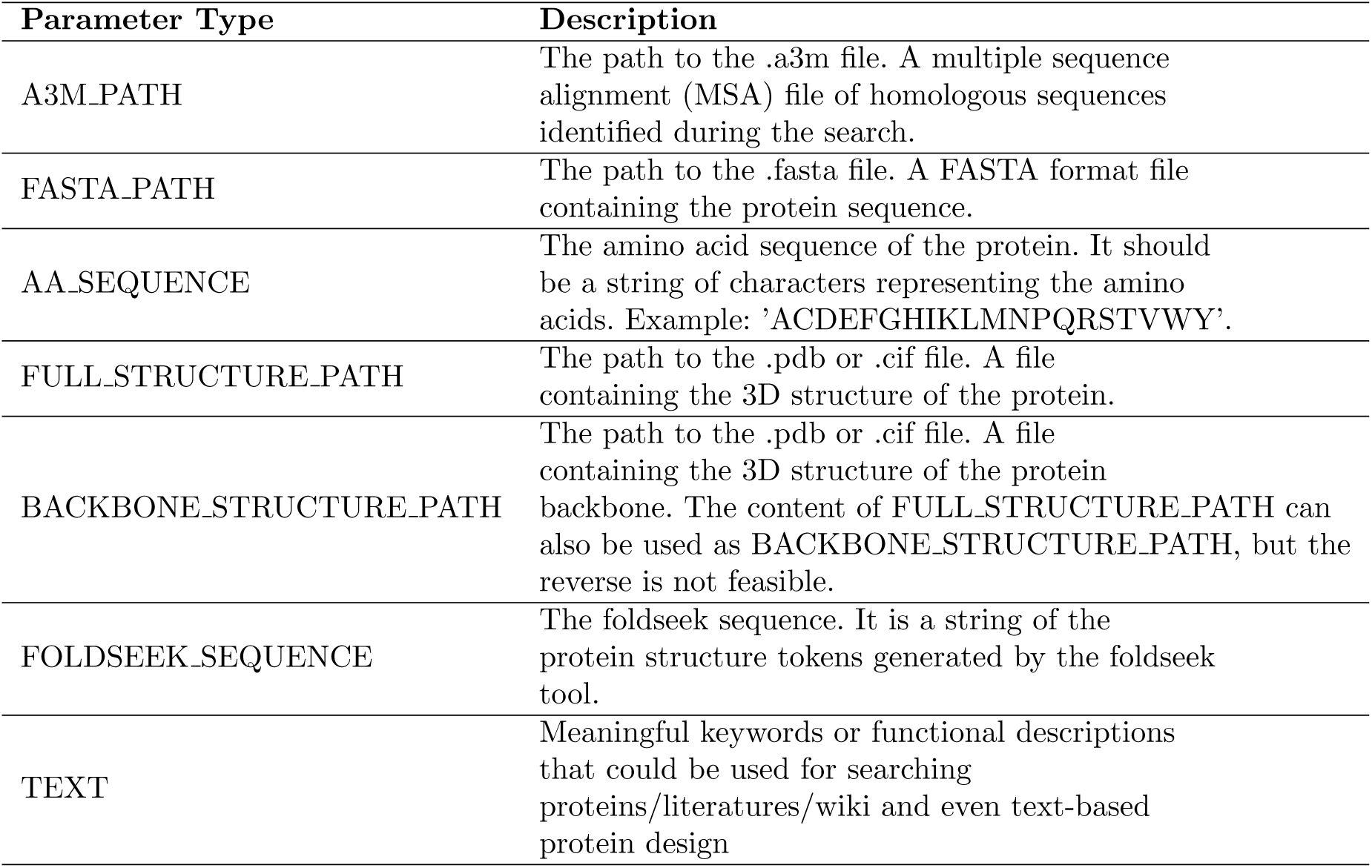
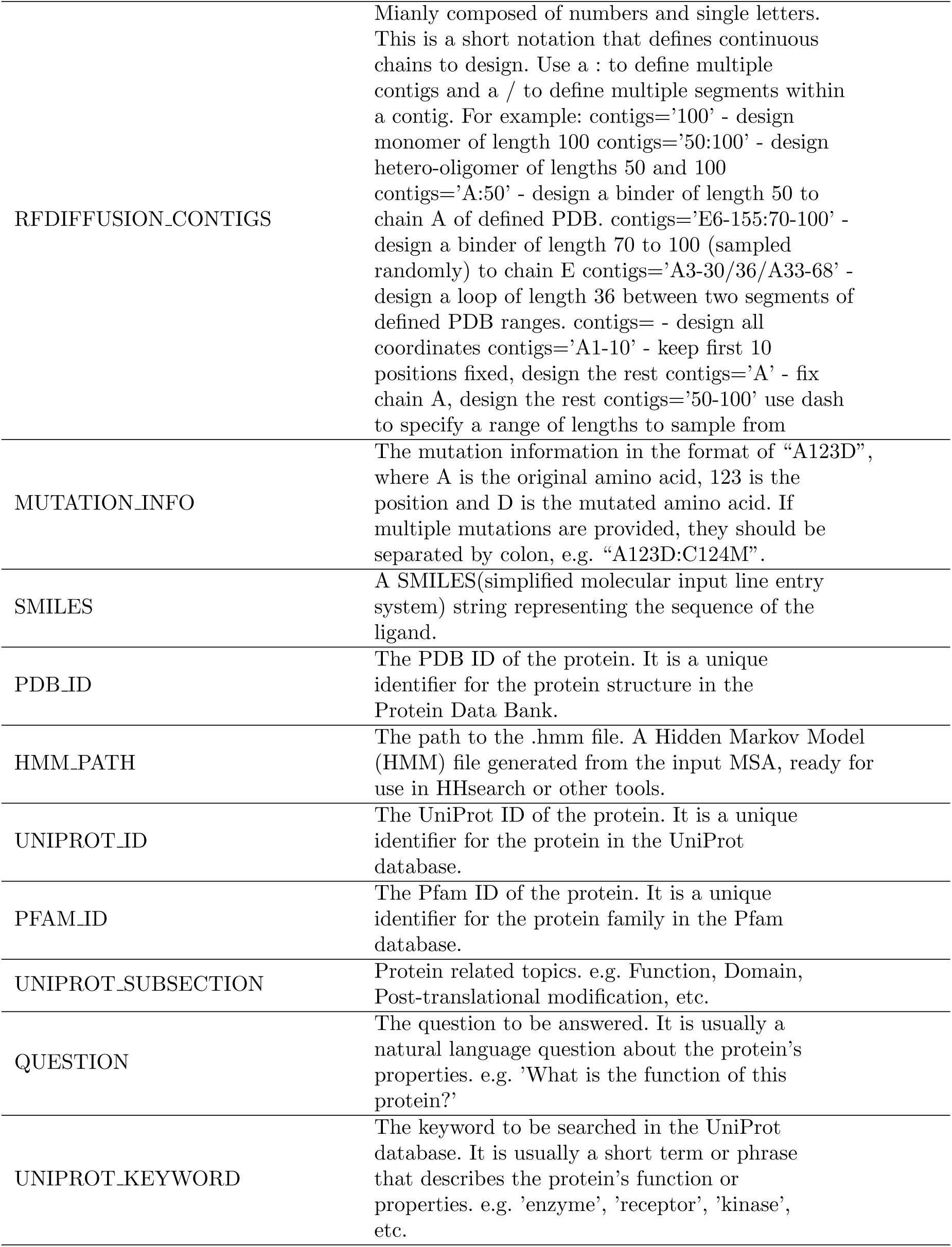
Description of parameter type. Used both in *query_parser* and test sample generation.

| Parameter Type | Description |
| --- | --- |
| A3M_PATH | The path to the .a3m file. A multiple sequence alignment (MSA) file of homologous sequences identified during the search. |
| FASTA_PATH | The path to the .fasta file. A FASTA format file containing the protein sequence. |
| AA_SEQUENCE | The amino acid sequence of the protein. It should be a string of characters representing the amino acids. Example: 'ACDEFGHIKLMNPQRSTVWY'. |
| FULL_STRUCTURE_PATH | The path to the .pdb or .cif file. A file containing the 3D structure of the protein. |
| BACKBONE_STRUCTURE_PATH | The path to the .pdb or .cif file. A file containing the 3D structure of the protein backbone. The content of FULL_STRUCTURE_PATH can also be used as BACKBONE_STRUCTURE_PATH, but the reverse is not feasible. |
| FOLDSEEK_SEQUENCE | The foldseek sequence. It is a string of the protein structure tokens generated by the foldseek tool. |
| TEXT | Meaningful keywords or functional descriptions that could be used for searching proteins/literatures/wiki and even text-based protein design |
| RFDIFFUSION_CONTIGS | Mainly composed of numbers and single letters. This is a short notation that defines continuous chains to design. Use a : to define multiple contigs and a / to define multiple segments within a contig. For example: contigs='100' - design monomer of length 100 contigs='50:100' - design hetero-oligomer of lengths 50 and 100 contigs='A:50' - design a binder of length 50 to chain A of defined PDB. contigs='E6-155:70-100' - design a binder of length 70 to 100 (sampled randomly) to chain E contigs='A3-30/36/A33-68' - design a loop of length 36 between two segments of defined PDB ranges. contigs= - design all coordinates contigs='A1-10' - keep first 10 positions fixed, design the rest contigs='A' - fix chain A, design the rest contigs='50-100' use dash to specify a range of lengths to sample from |
| MUTATION_INFO | The mutation information in the format of "A123D", where A is the original amino acid, 123 is the position and D is the mutated amino acid. If multiple mutations are provided, they should be separated by colon, e.g. "A123D:C124M". |
| SMILES | A SMILES(simplified molecular input line entry system) string representing the sequence of the ligand. |
| PDB_ID | The PDB ID of the protein. It is a unique identifier for the protein structure in the Protein Data Bank. |
| HMM_PATH | The path to the .hmm file. A Hidden Markov Model (HMM) file generated from the input MSA, ready for use in HHsearch or other tools. |
| UNIPROT_ID | The UniProt ID of the protein. It is a unique identifier for the protein in the UniProt database. |
| PFAM_ID | The Pfam ID of the protein. It is a unique identifier for the protein family in the Pfam database. |
| UNIPROT_SUBSECTION | Protein related topics. e.g. Function, Domain, Post-translational modification, etc. |
| QUESTION | The question to be answered. It is usually a natural language question about the protein's properties. e.g. 'What is the function of this protein?' |
| UNIPROT_KEYWORD | The keyword to be searched in the UniProt database. It is usually a short term or phrase that describes the protein's function or properties. e.g. 'enzyme', 'receptor', 'kinase', etc. |

**Supplementary Table 3:**
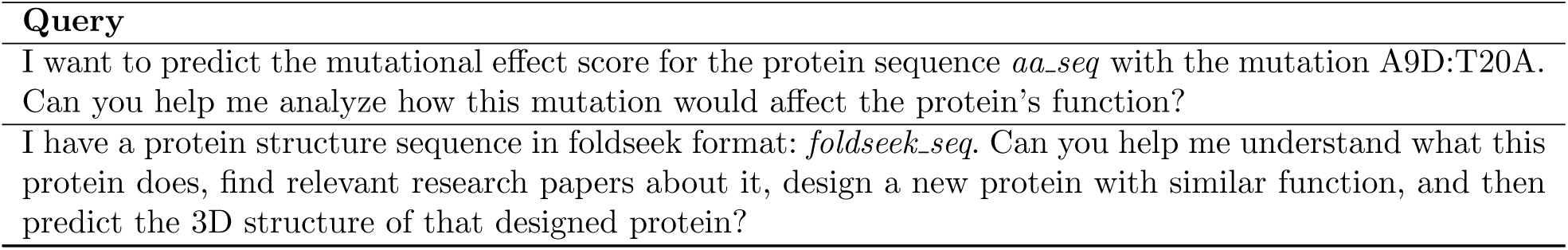
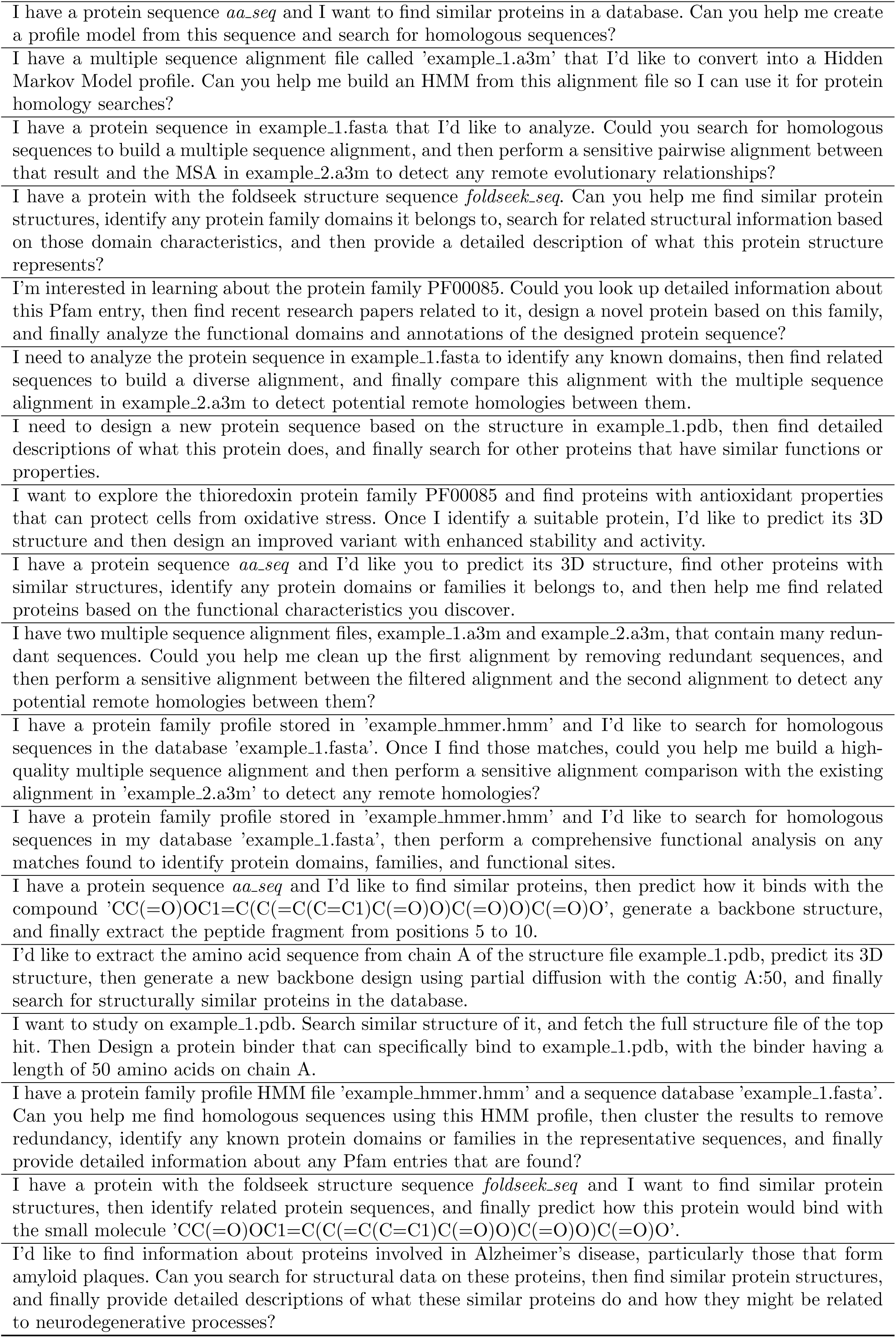
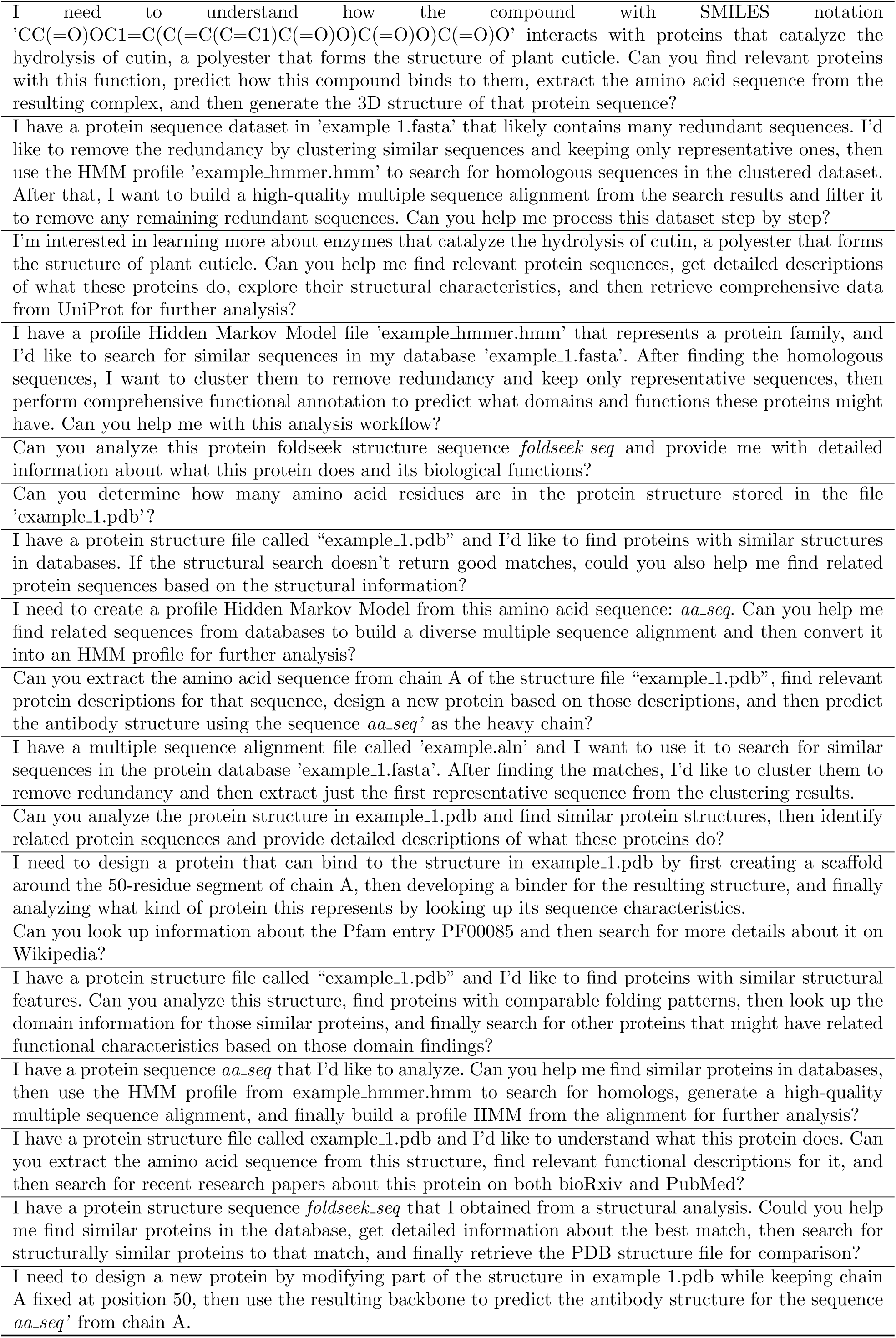
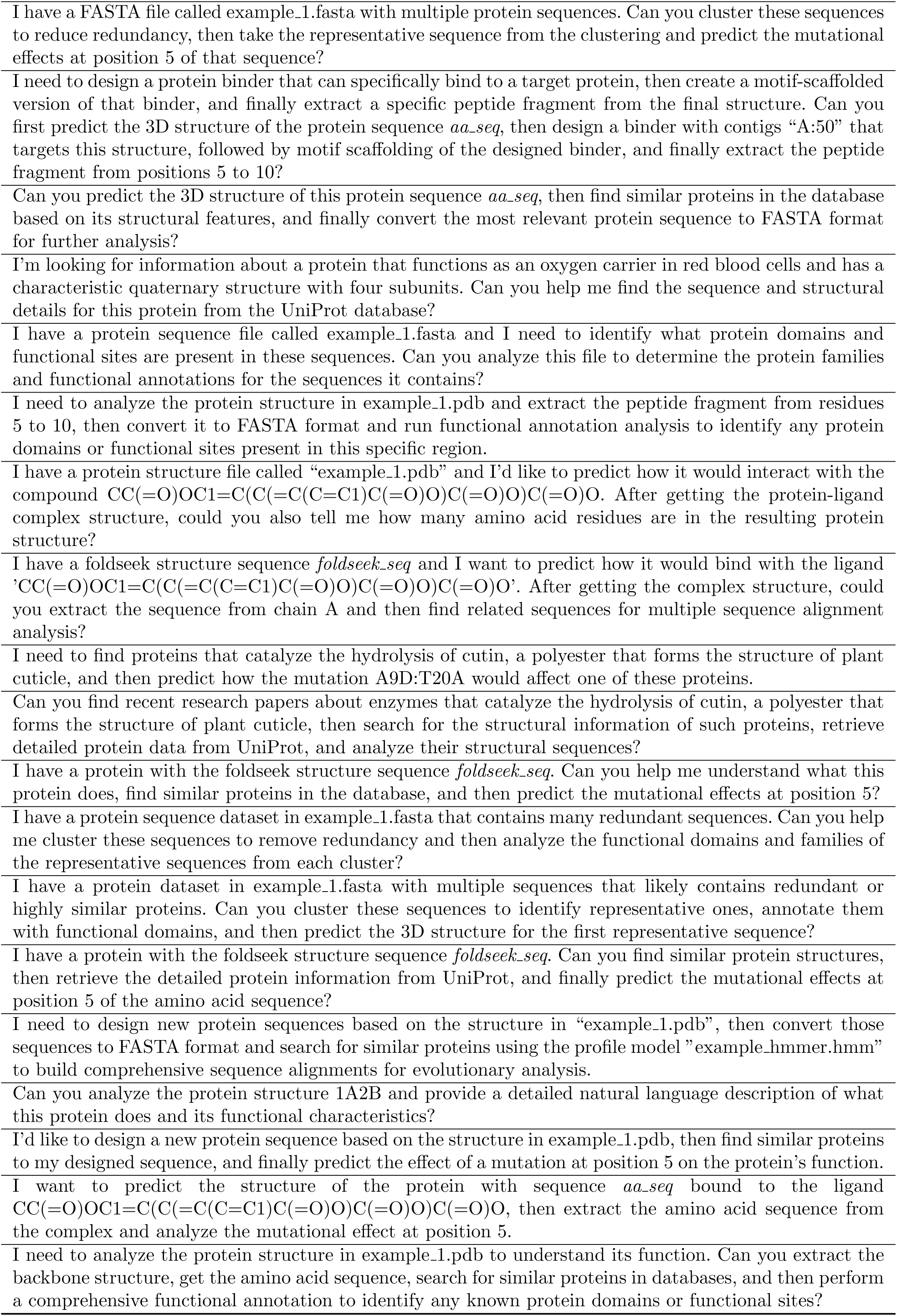
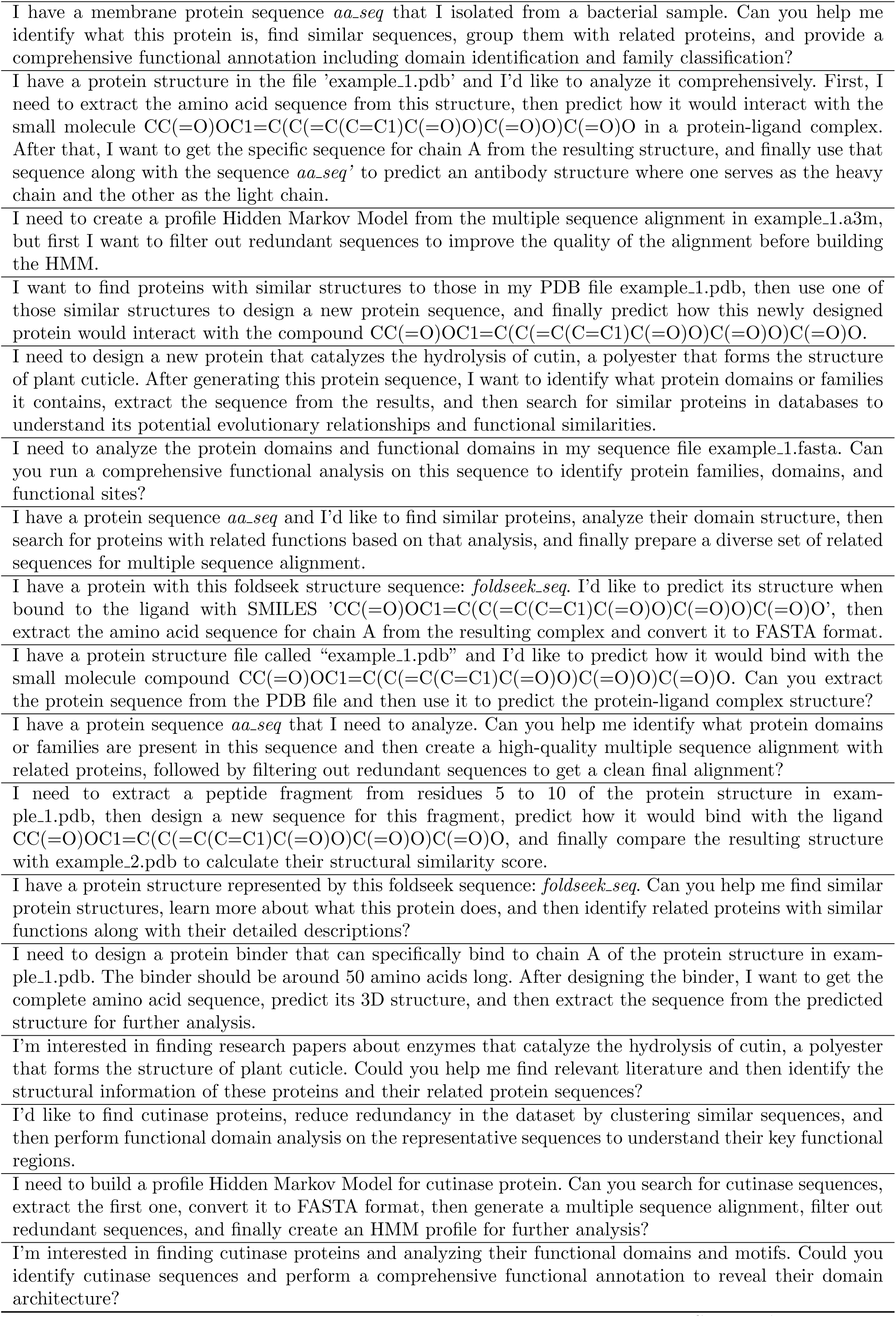
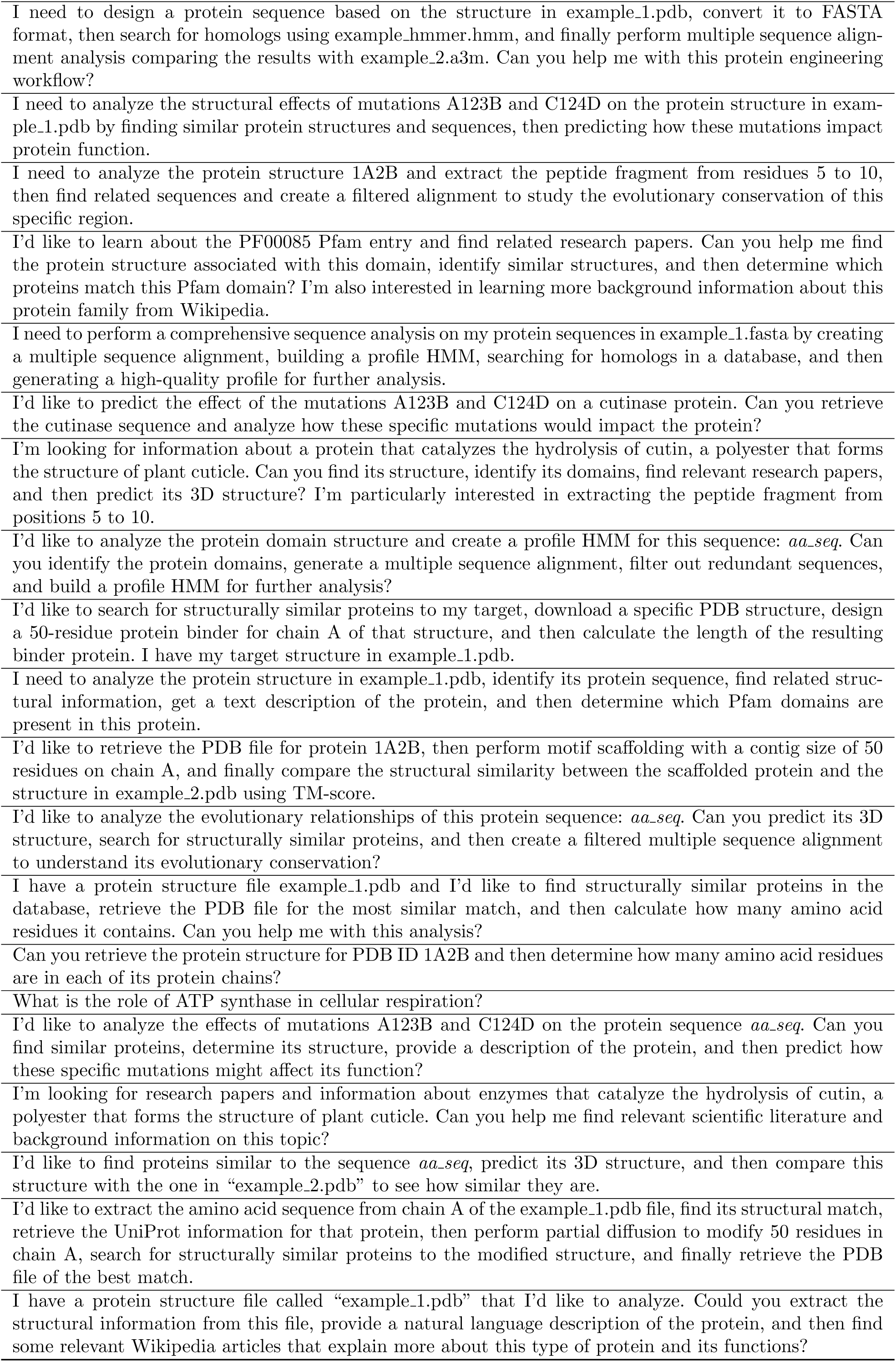
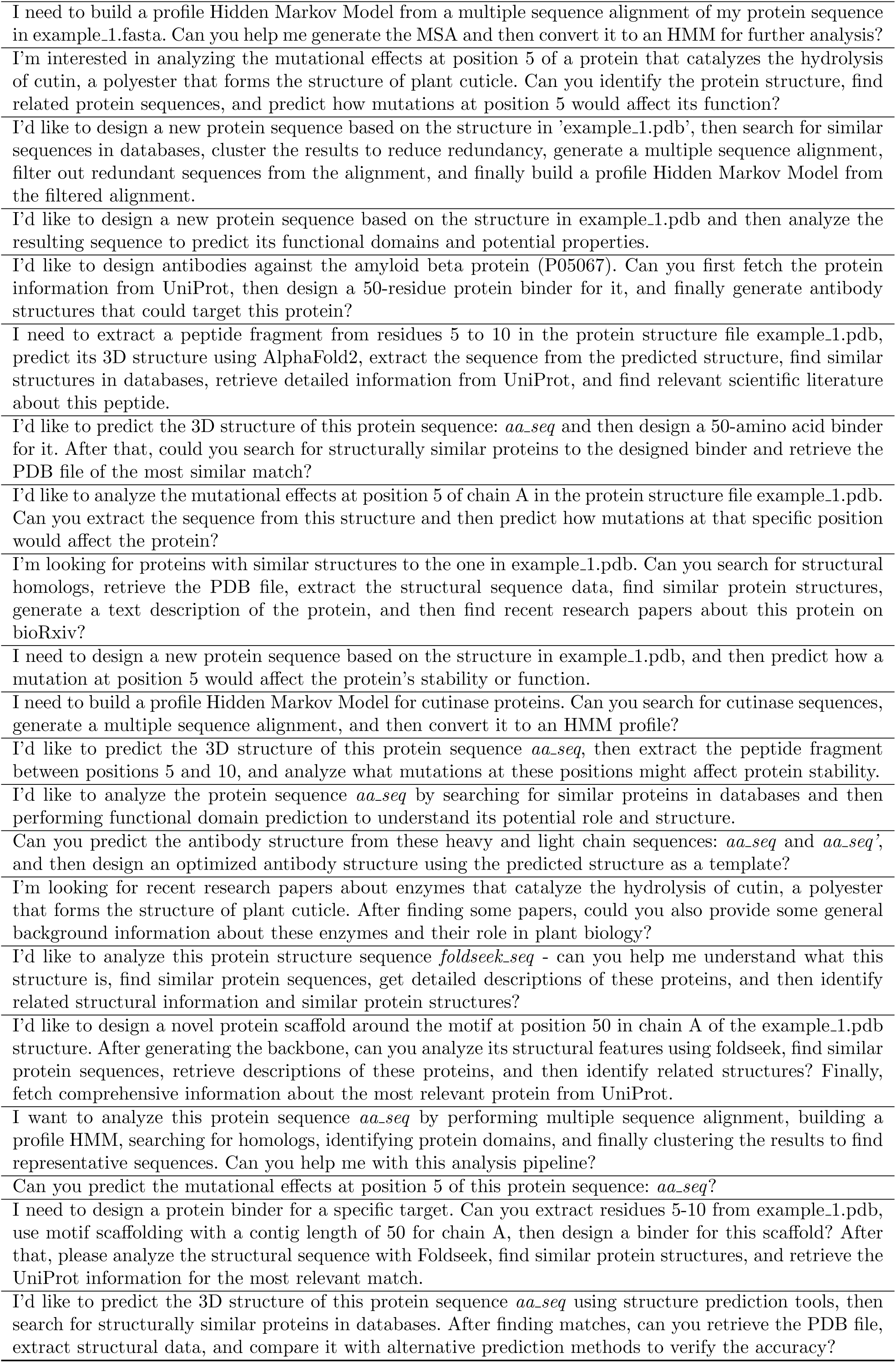
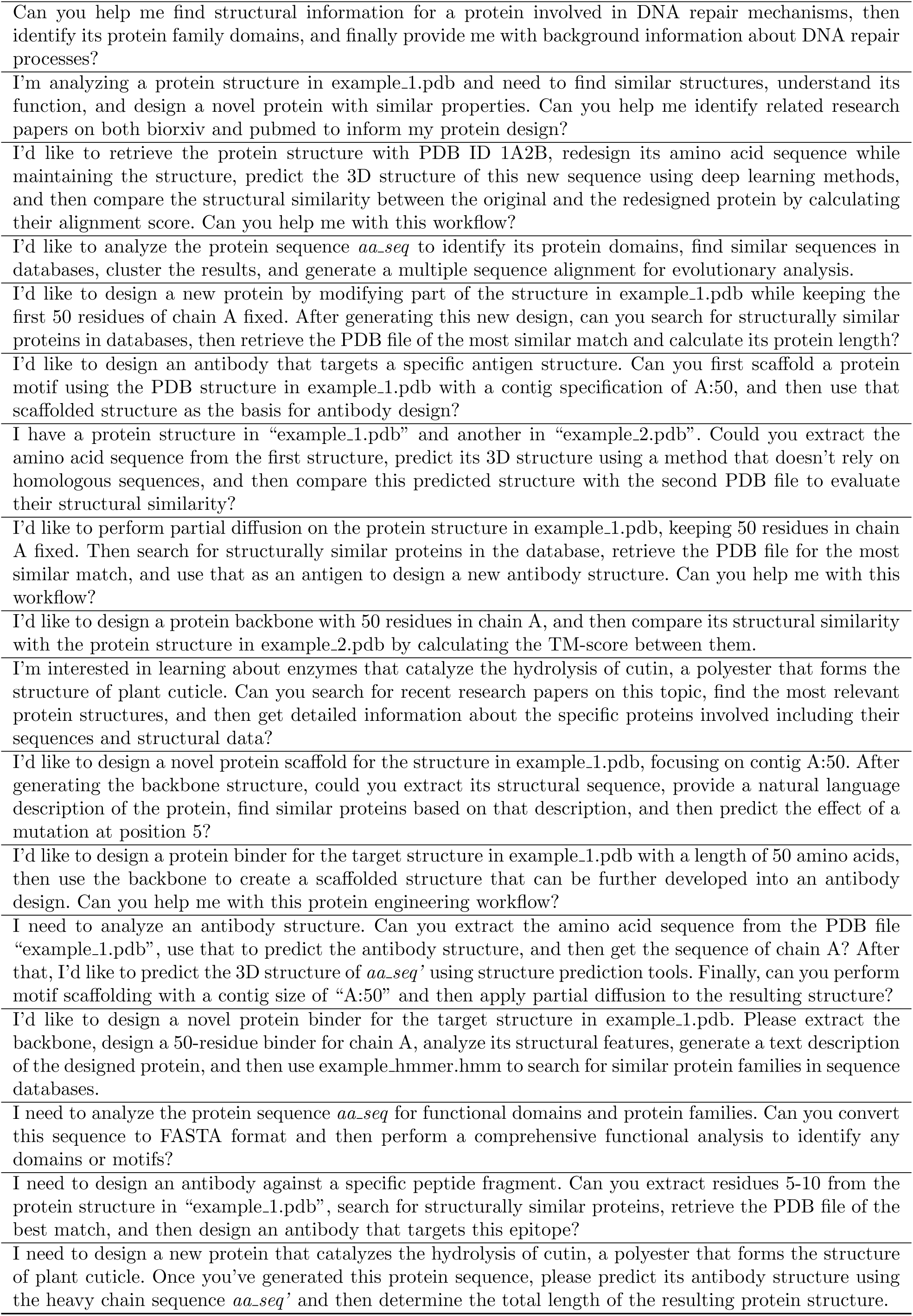
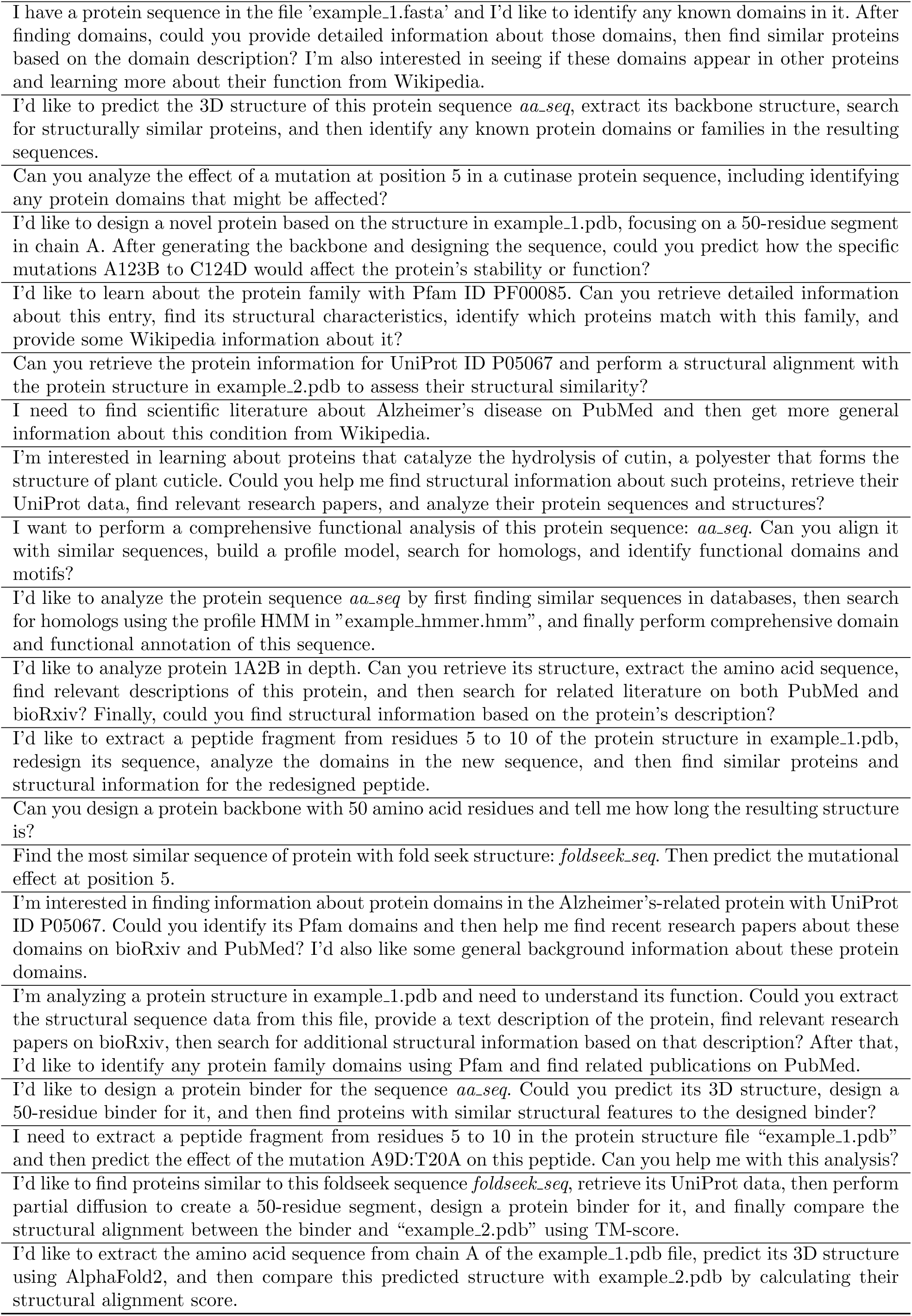
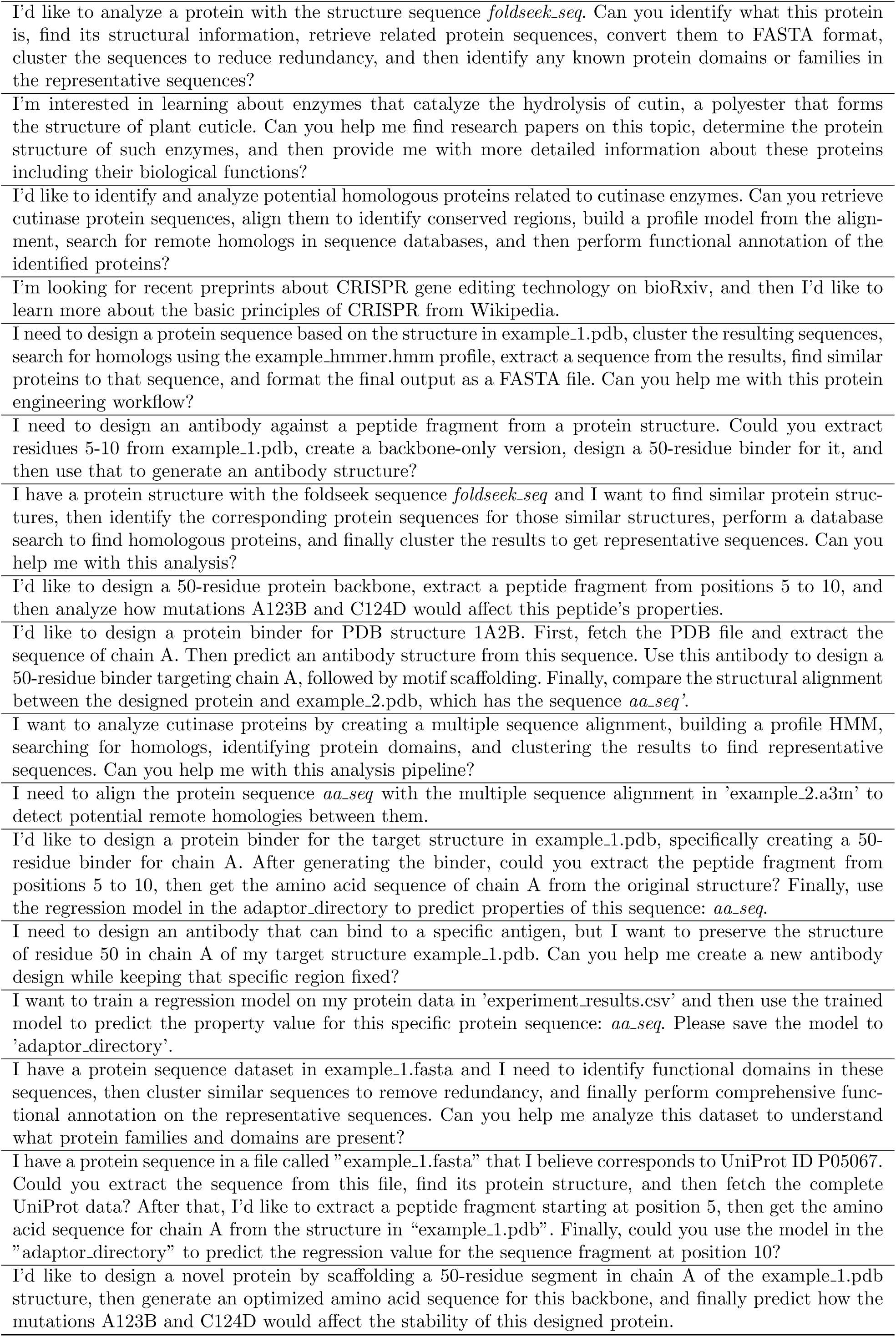
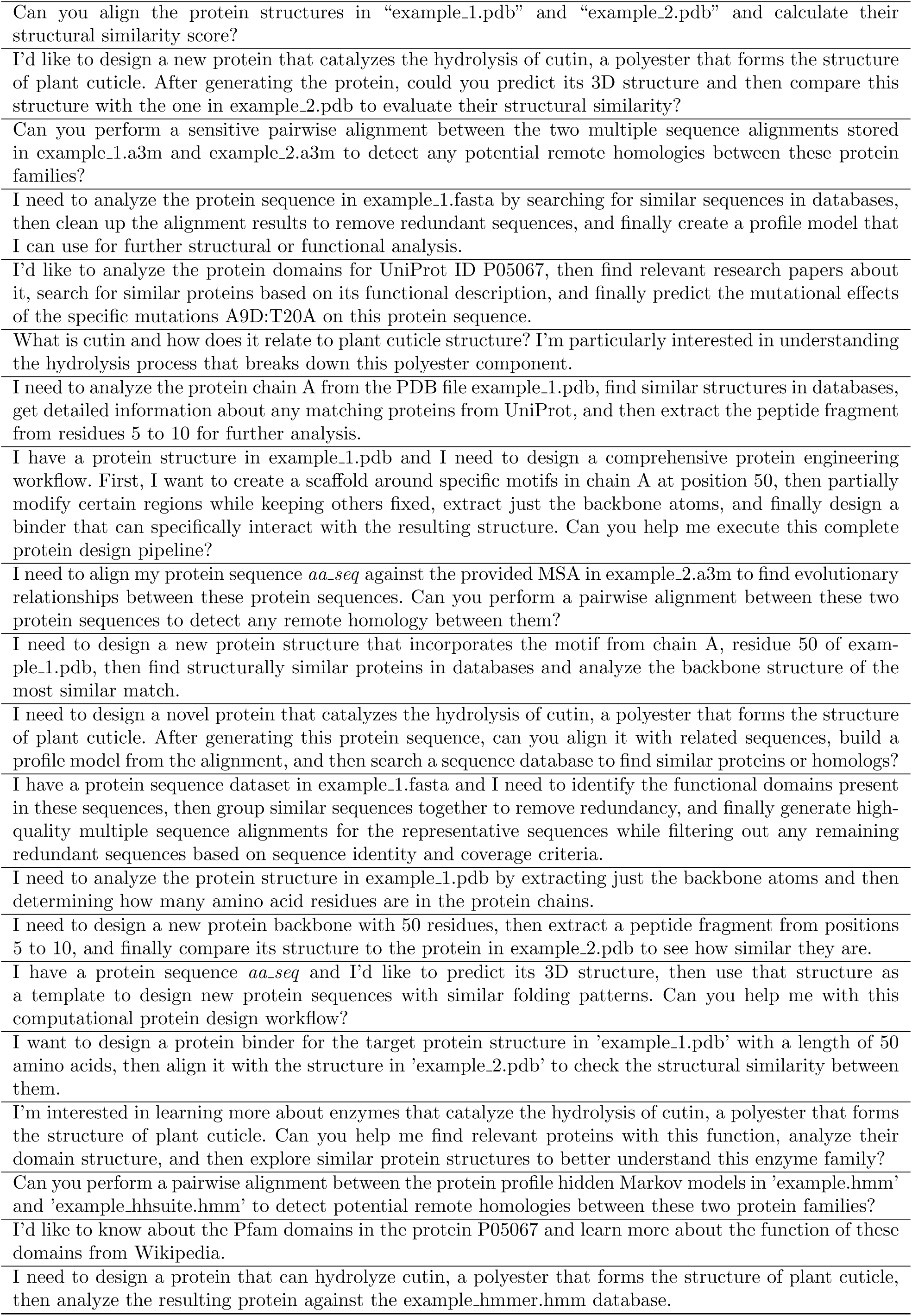
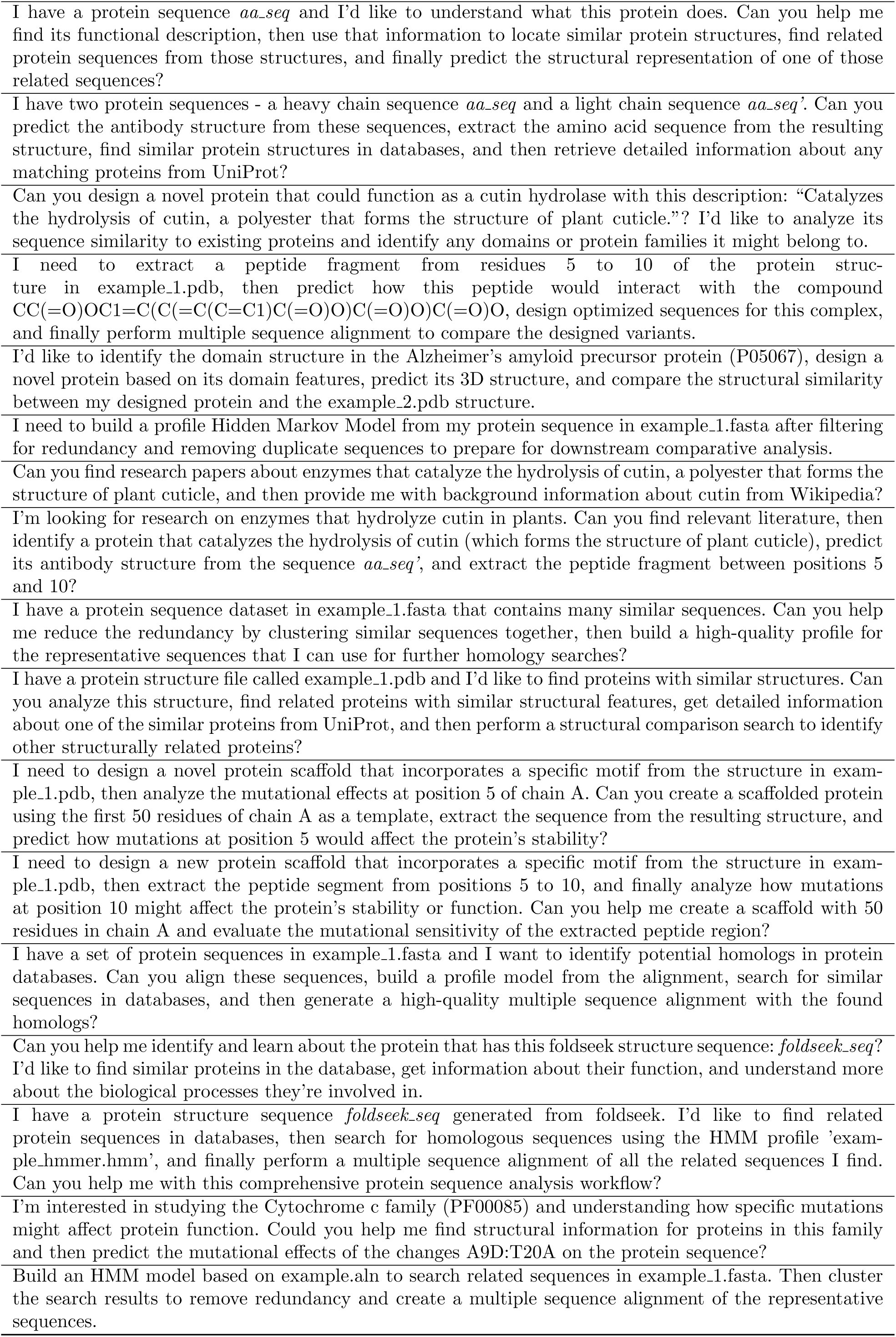
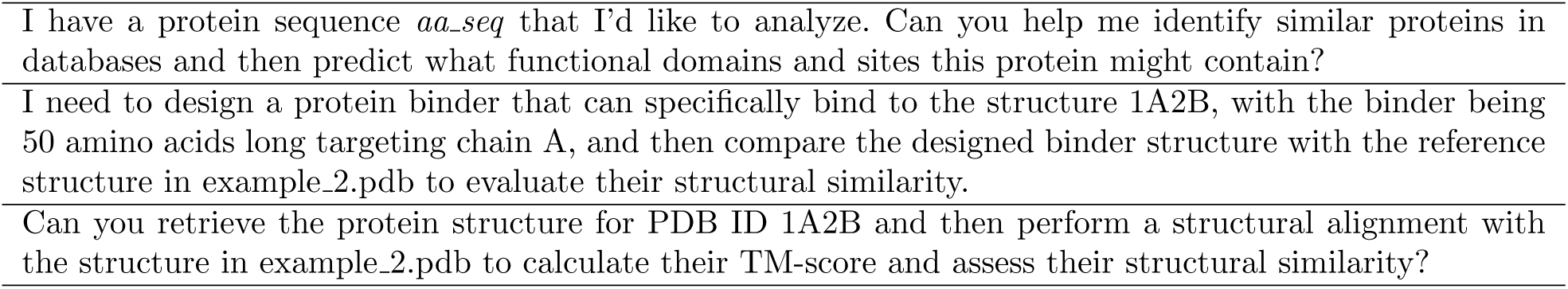
Benchmark Dataset of Complex Queries. These queries were generated to simulate real-world research challenges, spanning multiple protein research domains and typically requiring multi-step resolution.

**Supplementary Table 4:** Appendix: Raw Performance Evaluation Data. This table presents the detailed scores for the Gemini, Biomni, and PRIME models across four metrics on the benchmark of 213 complex queries.

| Query ID | Gemini |  |  |  | Biomni |  |  |  | PRIME |  |  |  |
| --- | --- | --- | --- | --- | --- | --- | --- | --- | --- | --- | --- | --- |
|  | Task | Tool | Hall. | Rob. | Task | Tool | Hall. | Rob. | Task | Tool | Hall. | Rob. |
| 1 | 0 | 0 | 0 | 0 | 2 | 1 | 1 | 2 | 2 | 2 | 1 | 2 |
| 2 | 0 | 0 | 1 | 0 | 2 | 1 | 1 | 1 | 2 | 1 | 1 | 2 |
| 3 | 1 | 1 | 0 | 0 | 2 | 1 | 1 | 2 | 2 | 2 | 1 | 2 |
| 4 | 2 | 1 | 1 | 1 | 1 | 0 | 0 | 1 | 2 | 2 | 1 | 2 |
| 5 | 2 | 1 | 1 | 2 | 2 | 2 | 1 | 2 | 2 | 1 | 1 | 2 |
| 6 | 0 | 1 | 1 | 0 | 2 | 1 | 1 | 2 | 1 | 0 | 1 | 1 |
| 7 | 2 | 0 | 1 | 0 | 2 | 1 | 1 | 2 | 2 | 2 | 1 | 2 |
| 8 | 2 | 0 | 1 | 0 | 1 | 1 | 1 | 1 | 2 | 1 | 1 | 2 |
| 9 | 2 | 1 | 1 | 1 | 1 | 1 | 1 | 1 | 2 | 2 | 1 | 2 |
| 10 | 1 | 0 | 0 | 0 | 2 | 1 | 1 | 2 | 2 | 2 | 1 | 2 |
| 11 | 1 | 1 | 0 | 0 | 2 | 2 | 1 | 2 | 2 | 2 | 1 | 2 |
| 12 | 1 | 1 | 0 | 0 | 2 | 1 | 1 | 2 | 2 | 2 | 1 | 2 |
| 13 | 1 | 1 | 1 | 1 | 2 | 1 | 1 | 2 | 1 | 1 | 1 | 2 |
| 14 | 0 | 0 | 1 | 0 | 2 | 1 | 1 | 2 | 1 | 0 | 0 | 1 |
| 15 | 2 | 0 | 1 | 0 | 2 | 1 | 1 | 2 | 2 | 2 | 1 | 2 |
| 16 | 2 | 0 | 1 | 0 | 2 | 1 | 1 | 2 | 2 | 1 | 1 | 2 |
| 17 | 1 | 0 | 0 | 0 | 2 | 1 | 1 | 2 | 2 | 1 | 1 | 2 |
| 18 | 1 | 1 | 1 | 1 | 2 | 1 | 1 | 2 | 2 | 2 | 1 | 2 |
| 19 | 1 | 1 | 1 | 1 | 2 | 1 | 1 | 2 | 2 | 1 | 1 | 2 |
| 20 | 2 | 1 | 1 | 1 | 2 | 1 | 1 | 2 | 2 | 1 | 1 | 2 |
| 21 | 2 | 1 | 1 | 2 | 2 | 0 | 1 | 1 | 2 | 2 | 1 | 2 |
| 22 | 0 | 0 | 0 | 1 | 1 | 1 | 1 | 2 | 0 | 1 | 1 | 1 |
| 23 | 2 | 0 | 1 | 0 | 2 | 1 | 1 | 2 | 2 | 1 | 1 | 2 |
| 24 | 2 | 0 | 1 | 0 | 2 | 1 | 1 | 2 | 2 | 1 | 1 | 2 |
| 25 | 1 | 1 | 0 | 2 | 2 | 2 | 1 | 2 | 2 | 2 | 1 | 2 |
| 26 | 2 | 0 | 1 | 0 | 2 | 1 | 1 | 2 | 1 | 0 | 1 | 1 |
| 27 | 1 | 0 | 0 | 0 | 2 | 1 | 1 | 2 | 1 | 1 | 1 | 1 |
| 28 | 1 | 0 | 0 | 0 | 2 | 1 | 1 | 2 | 2 | 1 | 1 | 2 |
| 29 | 1 | 1 | 1 | 0 | 2 | 1 | 1 | 2 | 2 | 2 | 1 | 2 |
| 30 | 1 | 1 | 1 | 1 | 2 | 1 | 1 | 2 | 2 | 2 | 1 | 2 |
| 31 | 0 | 1 | 1 | 0 | 2 | 1 | 1 | 2 | 0 | 0 | 1 | 1 |
| 32 | 1 | 1 | 1 | 1 | 2 | 1 | 1 | 2 | 2 | 1 | 1 | 2 |
| 33 | 0 | 0 | 1 | 0 | 2 | 1 | 1 | 2 | 1 | 1 | 1 | 2 |
| 34 | 2 | 1 | 1 | 2 | 2 | 1 | 1 | 2 | 2 | 1 | 0 | 2 |
| 35 | 1 | 1 | 1 | 1 | 2 | 1 | 1 | 2 | 2 | 2 | 1 | 2 |
| 36 | 1 | 0 | 1 | 0 | 2 | 1 | 1 | 2 | 2 | 2 | 1 | 2 |
| 37 | 2 | 0 | 1 | 0 | 2 | 1 | 1 | 2 | 2 | 1 | 1 | 2 |
| 38 | 1 | 1 | 1 | 1 | 2 | 1 | 1 | 2 | 2 | 2 | 1 | 2 |
| 39 | 1 | 2 | 1 | 1 | 0 | 0 | 1 | 0 | 2 | 2 | 1 | 2 |
| 40 | 1 | 1 | 1 | 0 | 1 | 0 | 1 | 0 | 2 | 2 | 1 | 2 |
| 41 | 0 | 0 | 0 | 0 | 0 | 0 | 1 | 0 | 2 | 2 | 1 | 2 |
| 42 | 1 | 1 | 1 | 1 | 2 | 1 | 1 | 2 | 2 | 2 | 1 | 2 |

Supplementary Table 4 – continued from previous page
| Query ID | Gemini |  |  |  | Biomni |  |  |  | PRIME |  |  |  |
| --- | --- | --- | --- | --- | --- | --- | --- | --- | --- | --- | --- | --- |
|  | Task | Tool | Hall. | Rob. | Task | Tool | Hall. | Rob. | Task | Tool | Hall. | Rob. |
| 43 | 1 | 1 | 1 | 1 | 2 | 0 | 1 | 1 | 2 | 1 | 1 | 2 |
| 44 | 1 | 0 | 0 | 1 | 2 | 1 | 1 | 1 | 2 | 2 | 1 | 2 |
| 45 | 0 | 0 | 1 | 0 | 2 | 2 | 1 | 2 | 1 | 1 | 1 | 1 |
| 46 | 1 | 1 | 1 | 0 | 2 | 2 | 1 | 2 | 2 | 2 | 1 | 2 |
| 47 | 1 | 1 | 1 | 1 | 2 | 1 | 1 | 2 | 2 | 2 | 1 | 2 |
| 48 | 0 | 1 | 1 | 1 | 2 | 1 | 1 | 2 | 2 | 1 | 1 | 2 |
| 49 | 2 | 0 | 1 | 0 | 2 | 1 | 1 | 2 | 2 | 1 | 1 | 2 |
| 50 | 1 | 0 | 0 | 0 | 2 | 1 | 1 | 2 | 2 | 2 | 1 | 2 |
| 51 | 1 | 0 | 0 | 0 | 2 | 1 | 1 | 2 | 2 | 2 | 1 | 2 |
| 52 | 0 | 0 | 0 | 0 | 0 | 0 | 1 | 1 | 0 | 0 | 0 | 0 |
| 53 | 1 | 0 | 0 | 0 | 2 | 1 | 1 | 2 | 2 | 2 | 1 | 2 |
| 54 | 2 | 1 | 1 | 2 | 2 | 1 | 1 | 2 | 2 | 2 | 1 | 2 |
| 55 | 1 | 0 | 0 | 0 | 2 | 1 | 1 | 2 | 2 | 2 | 1 | 2 |
| 56 | 1 | 1 | 1 | 0 | 2 | 1 | 1 | 2 | 2 | 1 | 1 | 2 |
| 57 | 1 | 0 | 0 | 1 | 2 | 2 | 1 | 2 | 2 | 2 | 1 | 2 |
| 58 | 1 | 1 | 1 | 1 | 2 | 1 | 1 | 2 | 2 | 1 | 1 | 2 |
| 59 | 0 | 0 | 1 | 0 | 2 | 1 | 1 | 2 | 1 | 2 | 1 | 1 |
| 60 | 1 | 2 | 0 | 1 | 2 | 1 | 1 | 2 | 2 | 2 | 1 | 2 |
| 61 | 0 | 0 | 0 | 0 | 2 | 1 | 1 | 2 | 2 | 2 | 1 | 2 |
| 62 | 1 | 0 | 0 | 1 | 2 | 2 | 1 | 2 | 1 | 2 | 0 | 2 |
| 63 | 1 | 1 | 1 | 1 | 2 | 1 | 1 | 2 | 2 | 1 | 1 | 2 |
| 64 | 0 | 1 | 1 | 1 | 1 | 0 | 1 | 2 | 1 | 1 | 1 | 2 |
| 65 | 1 | 0 | 1 | 1 | 2 | 1 | 0 | 2 | 2 | 2 | 1 | 2 |
| 66 | 1 | 1 | 1 | 0 | 0 | 0 | 0 | 1 | 1 | 1 | 1 | 2 |
| 67 | 2 | 1 | 1 | 1 | 2 | 1 | 1 | 2 | 2 | 1 | 1 | 2 |
| 68 | 1 | 1 | 1 | 1 | 2 | 1 | 1 | 2 | 0 | 0 | 0 | 0 |
| 69 | 1 | 1 | 1 | 1 | 2 | 1 | 1 | 2 | 2 | 1 | 1 | 2 |
| 70 | 1 | 1 | 1 | 0 | 1 | 0 | 1 | 1 | 1 | 1 | 1 | 2 |
| 71 | 1 | 0 | 1 | 1 | 2 | 1 | 1 | 2 | 2 | 1 | 1 | 2 |
| 72 | 0 | 1 | 1 | 1 | 2 | 1 | 1 | 2 | 2 | 2 | 1 | 2 |
| 73 | 1 | 0 | 0 | 0 | 2 | 1 | 1 | 2 | 2 | 2 | 1 | 2 |
| 74 | 2 | 0 | 0 | 0 | 2 | 1 | 1 | 2 | 2 | 2 | 1 | 2 |
| 75 | 0 | 1 | 1 | 0 | 2 | 1 | 1 | 2 | 2 | 2 | 1 | 2 |
| 76 | 1 | 0 | 1 | 0 | 2 | 1 | 1 | 2 | 2 | 2 | 1 | 2 |
| 77 | 1 | 0 | 0 | 0 | 2 | 2 | 1 | 2 | 2 | 2 | 1 | 2 |
| 78 | 1 | 1 | 1 | 1 | 2 | 1 | 1 | 2 | 2 | 2 | 1 | 2 |
| 79 | 0 | 0 | 1 | 0 | 2 | 2 | 1 | 2 | 2 | 1 | 1 | 2 |
| 80 | 0 | 2 | 1 | 1 | 2 | 1 | 1 | 2 | 0 | 0 | 0 | 0 |
| 81 | 1 | 1 | 1 | 2 | 1 | 1 | 1 | 1 | 2 | 1 | 1 | 2 |
| 82 | 1 | 1 | 0 | 0 | 2 | 1 | 1 | 2 | 2 | 2 | 1 | 2 |
| 83 | 2 | 0 | 1 | 0 | 2 | 1 | 1 | 2 | 2 | 2 | 1 | 2 |
| 84 | 1 | 0 | 0 | 0 | 1 | 1 | 1 | 2 | 0 | 0 | 1 | 1 |
| 85 | 2 | 0 | 1 | 0 | 2 | 1 | 1 | 2 | 2 | 1 | 1 | 2 |
| 86 | 1 | 1 | 1 | 1 | 2 | 1 | 1 | 2 | 2 | 2 | 1 | 2 |
| 87 | 2 | 1 | 1 | 2 | 1 | 1 | 1 | 2 | 2 | 2 | 1 | 2 |
| 88 | 0 | 0 | 1 | 1 | 2 | 1 | 1 | 2 | 2 | 1 | 1 | 2 |
| 89 | 1 | 1 | 1 | 1 | 2 | 1 | 1 | 2 | 2 | 2 | 1 | 2 |
| 90 | 1 | 1 | 1 | 1 | 2 | 1 | 1 | 2 | 0 | 0 | 1 | 1 |
| 91 | 1 | 0 | 0 | 0 | 2 | 1 | 1 | 2 | 2 | 2 | 1 | 2 |
| 92 | 1 | 1 | 1 | 1 | 2 | 0 | 1 | 1 | 2 | 1 | 1 | 2 |
| 93 | 1 | 0 | 0 | 1 | 2 | 1 | 1 | 2 | 1 | 1 | 1 | 1 |
| 94 | 1 | 2 | 1 | 1 | 2 | 0 | 1 | 2 | 1 | 1 | 1 | 2 |
| 95 | 2 | 1 | 1 | 2 | 2 | 1 | 1 | 2 | 2 | 1 | 1 | 2 |
| 96 | 1 | 1 | 1 | 1 | 2 | 1 | 1 | 2 | 2 | 2 | 1 | 2 |
| 97 | 0 | 0 | 0 | 0 | 2 | 1 | 1 | 2 | 2 | 2 | 1 | 2 |
| 98 | 1 | 0 | 1 | 1 | 2 | 1 | 1 | 2 | 2 | 1 | 1 | 2 |

Supplementary Table 4 – continued from previous page
| Query ID | Gemini |  |  |  | Biomni |  |  |  | PRIME |  |  |  |
| --- | --- | --- | --- | --- | --- | --- | --- | --- | --- | --- | --- | --- |
|  | Task | Tool | Hall. | Rob. | Task | Tool | Hall. | Rob. | Task | Tool | Hall. | Rob. |
| 99 | 2 | 1 | 1 | 2 | 1 | 0 | 1 | 1 | 2 | 2 | 1 | 2 |
| 100 | 1 | 1 | 1 | 1 | 1 | 0 | 0 | 1 | 2 | 2 | 1 | 2 |
| 101 | 1 | 1 | 0 | 1 | 2 | 0 | 1 | 1 | 2 | 2 | 1 | 2 |
| 102 | 0 | 1 | 0 | 0 | 1 | 0 | 0 | 1 | 2 | 2 | 1 | 2 |
| 103 | 0 | 0 | 0 | 0 | 1 | 0 | 1 | 1 | 2 | 2 | 1 | 2 |
| 104 | 2 | 0 | 1 | 1 | 2 | 2 | 1 | 2 | 2 | 1 | 1 | 2 |
| 105 | 1 | 1 | 1 | 1 | 2 | 1 | 1 | 2 | 2 | 1 | 1 | 2 |
| 106 | 1 | 0 | 0 | 1 | 2 | 1 | 1 | 2 | 2 | 2 | 1 | 2 |
| 107 | 2 | 1 | 1 | 1 | 2 | 0 | 1 | 2 | 2 | 2 | 1 | 2 |
| 108 | 0 | 1 | 0 | 0 | 1 | 1 | 1 | 2 | 2 | 2 | 1 | 2 |
| 109 | 1 | 0 | 0 | 0 | 2 | 1 | 1 | 2 | 2 | 2 | 1 | 2 |
| 110 | 2 | 2 | 1 | 2 | 1 | 0 | 1 | 0 | 2 | 2 | 1 | 2 |
| 111 | 1 | 1 | 1 | 1 | 2 | 1 | 1 | 2 | 2 | 1 | 1 | 1 |
| 112 | 1 | 1 | 1 | 1 | 2 | 2 | 1 | 2 | 2 | 2 | 1 | 2 |
| 113 | 2 | 2 | 1 | 2 | 2 | 1 | 1 | 2 | 2 | 2 | 1 | 2 |
| 114 | 1 | 2 | 1 | 2 | 2 | 2 | 1 | 2 | 2 | 2 | 1 | 2 |
| 115 | 1 | 2 | 1 | 1 | 2 | 1 | 1 | 2 | 2 | 2 | 1 | 2 |
| 116 | 2 | 2 | 1 | 2 | 1 | 0 | 0 | 1 | 2 | 2 | 1 | 2 |
| 117 | 1 | 0 | 0 | 1 | 2 | 1 | 1 | 2 | 2 | 2 | 1 | 2 |
| 118 | 2 | 1 | 1 | 2 | 2 | 1 | 1 | 2 | 2 | 2 | 1 | 2 |
| 119 | 1 | 2 | 1 | 2 | 2 | 1 | 1 | 2 | 2 | 2 | 1 | 2 |
| 120 | 1 | 1 | 1 | 0 | 2 | 1 | 1 | 2 | 2 | 2 | 1 | 2 |
| 121 | 0 | 1 | 1 | 0 | 2 | 2 | 1 | 2 | 2 | 2 | 1 | 2 |
| 122 | 0 | 0 | 0 | 0 | 2 | 1 | 1 | 2 | 2 | 2 | 1 | 2 |
| 123 | 0 | 1 | 0 | 1 | 1 | 0 | 0 | 1 | 2 | 2 | 1 | 2 |
| 124 | 1 | 1 | 1 | 1 | 1 | 1 | 1 | 1 | 2 | 2 | 1 | 2 |
| 125 | 2 | 0 | 0 | 0 | 2 | 0 | 1 | 2 | 2 | 2 | 1 | 2 |
| 126 | 2 | 1 | 1 | 1 | 2 | 1 | 1 | 2 | 2 | 2 | 1 | 2 |
| 127 | 1 | 2 | 1 | 1 | 2 | 1 | 1 | 2 | 2 | 2 | 1 | 2 |
| 128 | 1 | 0 | 1 | 0 | 2 | 1 | 1 | 2 | 2 | 1 | 1 | 2 |
| 129 | 1 | 0 | 1 | 0 | 2 | 1 | 1 | 2 | 2 | 2 | 1 | 2 |
| 130 | 1 | 0 | 1 | 0 | 2 | 1 | 1 | 2 | 2 | 2 | 1 | 2 |
| 131 | 0 | 0 | 0 | 0 | 2 | 1 | 1 | 2 | 2 | 1 | 1 | 2 |
| 132 | 1 | 2 | 1 | 2 | 2 | 1 | 1 | 2 | 1 | 1 | 1 | 2 |
| 133 | 0 | 1 | 1 | 0 | 2 | 2 | 1 | 2 | 2 | 2 | 1 | 2 |
| 134 | 2 | 0 | 1 | 0 | 1 | 0 | 1 | 1 | 2 | 2 | 1 | 2 |
| 135 | 1 | 1 | 1 | 1 | 1 | 1 | 1 | 1 | 2 | 1 | 1 | 2 |
| 136 | 1 | 0 | 0 | 0 | 2 | 1 | 1 | 2 | 1 | 2 | 1 | 2 |
| 137 | 2 | 0 | 1 | 0 | 2 | 2 | 1 | 2 | 2 | 1 | 1 | 2 |
| 138 | 0 | 0 | 0 | 0 | 2 | 1 | 1 | 2 | 2 | 2 | 1 | 2 |
| 139 | 1 | 0 | 0 | 1 | 2 | 1 | 1 | 2 | 2 | 2 | 1 | 2 |
| 140 | 2 | 1 | 1 | 2 | 2 | 1 | 1 | 2 | 2 | 2 | 1 | 2 |
| 141 | 1 | 1 | 1 | 1 | 2 | 1 | 1 | 2 | 2 | 2 | 1 | 2 |
| 142 | 1 | 0 | 0 | 0 | 1 | 1 | 1 | 1 | 2 | 2 | 1 | 2 |
| 143 | 1 | 1 | 1 | 2 | 1 | 0 | 0 | 1 | 2 | 2 | 1 | 2 |
| 144 | 2 | 0 | 1 | 0 | 2 | 1 | 1 | 2 | 2 | 1 | 1 | 2 |
| 145 | 2 | 1 | 1 | 2 | 2 | 2 | 1 | 2 | 2 | 2 | 1 | 2 |
| 146 | 0 | 1 | 1 | 0 | 2 | 2 | 1 | 1 | 2 | 2 | 1 | 2 |
| 147 | 1 | 2 | 1 | 2 | 2 | 0 | 0 | 1 | 2 | 2 | 1 | 2 |
| 148 | 1 | 1 | 1 | 1 | 2 | 1 | 1 | 1 | 2 | 2 | 1 | 2 |
| 149 | 1 | 1 | 1 | 1 | 2 | 1 | 1 | 2 | 2 | 2 | 1 | 2 |
| 150 | 1 | 2 | 1 | 1 | 2 | 2 | 1 | 2 | 2 | 2 | 1 | 2 |
| 151 | 1 | 1 | 0 | 0 | 2 | 1 | 1 | 2 | 2 | 2 | 1 | 2 |
| 152 | 1 | 1 | 0 | 0 | 2 | 1 | 1 | 2 | 2 | 2 | 1 | 2 |
| 153 | 1 | 0 | 1 | 1 | 0 | 1 | 1 | 1 | 2 | 1 | 1 | 2 |
| 154 | 1 | 0 | 0 | 0 | 2 | 2 | 1 | 2 | 2 | 2 | 1 | 1 |

Supplementary Table 4 – continued from previous page
| Query ID | Gemini |  |  |  | Biomni |  |  |  | PRIME |  |  |  |
| --- | --- | --- | --- | --- | --- | --- | --- | --- | --- | --- | --- | --- |
|  | Task | Tool | Hall. | Rob. | Task | Tool | Hall. | Rob. | Task | Tool | Hall. | Rob. |
| 155 | 1 | 2 | 1 | 2 | 2 | 1 | 1 | 2 | 2 | 2 | 1 | 2 |
| 156 | 1 | 1 | 0 | 0 | 2 | 2 | 1 | 2 | 2 | 2 | 1 | 2 |
| 157 | 1 | 1 | 1 | 0 | 2 | 1 | 1 | 2 | 2 | 2 | 1 | 2 |
| 158 | 1 | 2 | 1 | 1 | 2 | 2 | 1 | 2 | 2 | 2 | 1 | 2 |
| 159 | 1 | 0 | 0 | 0 | 2 | 1 | 1 | 2 | 2 | 2 | 1 | 2 |
| 160 | 0 | 1 | 1 | 0 | 1 | 0 | 0 | 1 | 2 | 2 | 1 | 2 |
| 161 | 1 | 2 | 1 | 1 | 2 | 1 | 1 | 2 | 2 | 2 | 1 | 2 |
| 162 | 0 | 1 | 1 | 0 | 2 | 2 | 1 | 2 | 2 | 2 | 1 | 2 |
| 163 | 1 | 0 | 1 | 1 | 2 | 1 | 1 | 2 | 2 | 2 | 1 | 2 |
| 164 | 1 | 0 | 0 | 0 | 2 | 1 | 1 | 1 | 2 | 2 | 1 | 2 |
| 165 | 1 | 1 | 1 | 1 | 1 | 1 | 1 | 2 | 2 | 2 | 1 | 2 |
| 166 | 2 | 1 | 1 | 2 | 2 | 2 | 1 | 2 | 2 | 1 | 1 | 2 |
| 167 | 0 | 1 | 0 | 0 | 2 | 2 | 1 | 2 | 2 | 2 | 1 | 2 |
| 168 | 0 | 1 | 1 | 0 | 2 | 1 | 1 | 2 | 2 | 2 | 1 | 2 |
| 169 | 2 | 0 | 1 | 0 | 2 | 1 | 1 | 2 | 2 | 1 | 1 | 2 |
| 170 | 1 | 1 | 0 | 1 | 2 | 2 | 1 | 2 | 2 | 2 | 1 | 2 |
| 171 | 2 | 1 | 1 | 2 | 2 | 2 | 1 | 2 | 2 | 2 | 1 | 2 |
| 172 | 1 | 1 | 1 | 1 | 2 | 2 | 1 | 2 | 2 | 2 | 1 | 2 |
| 173 | 1 | 0 | 0 | 0 | 1 | 1 | 1 | 2 | 2 | 2 | 1 | 2 |
| 174 | 0 | 1 | 1 | 0 | 2 | 1 | 1 | 2 | 2 | 1 | 1 | 2 |
| 175 | 0 | 0 | 0 | 0 | 1 | 1 | 1 | 2 | 2 | 1 | 1 | 2 |
| 176 | 1 | 2 | 1 | 1 | 2 | 1 | 1 | 2 | 2 | 2 | 1 | 2 |
| 177 | 1 | 0 | 1 | 1 | 1 | 0 | 0 | 1 | 2 | 2 | 1 | 2 |
| 178 | 1 | 0 | 1 | 1 | 2 | 0 | 0 | 1 | 2 | 2 | 1 | 2 |
| 179 | 0 | 1 | 0 | 0 | 2 | 1 | 0 | 2 | 1 | 2 | 0 | 2 |
| 180 | 1 | 1 | 1 | 0 | 2 | 1 | 1 | 2 | 2 | 1 | 1 | 2 |
| 181 | 2 | 1 | 1 | 1 | 2 | 0 | 0 | 1 | 2 | 2 | 1 | 2 |
| 182 | 2 | 1 | 1 | 2 | 2 | 0 | 1 | 1 | 2 | 1 | 1 | 2 |
| 183 | 2 | 1 | 1 | 2 | 2 | 1 | 1 | 1 | 2 | 2 | 1 | 2 |
| 184 | 0 | 1 | 0 | 0 | 2 | 1 | 1 | 2 | 2 | 2 | 1 | 2 |
| 185 | 0 | 0 | 0 | 0 | 2 | 1 | 1 | 2 | 2 | 2 | 1 | 2 |

**Supplementary Table 5.**
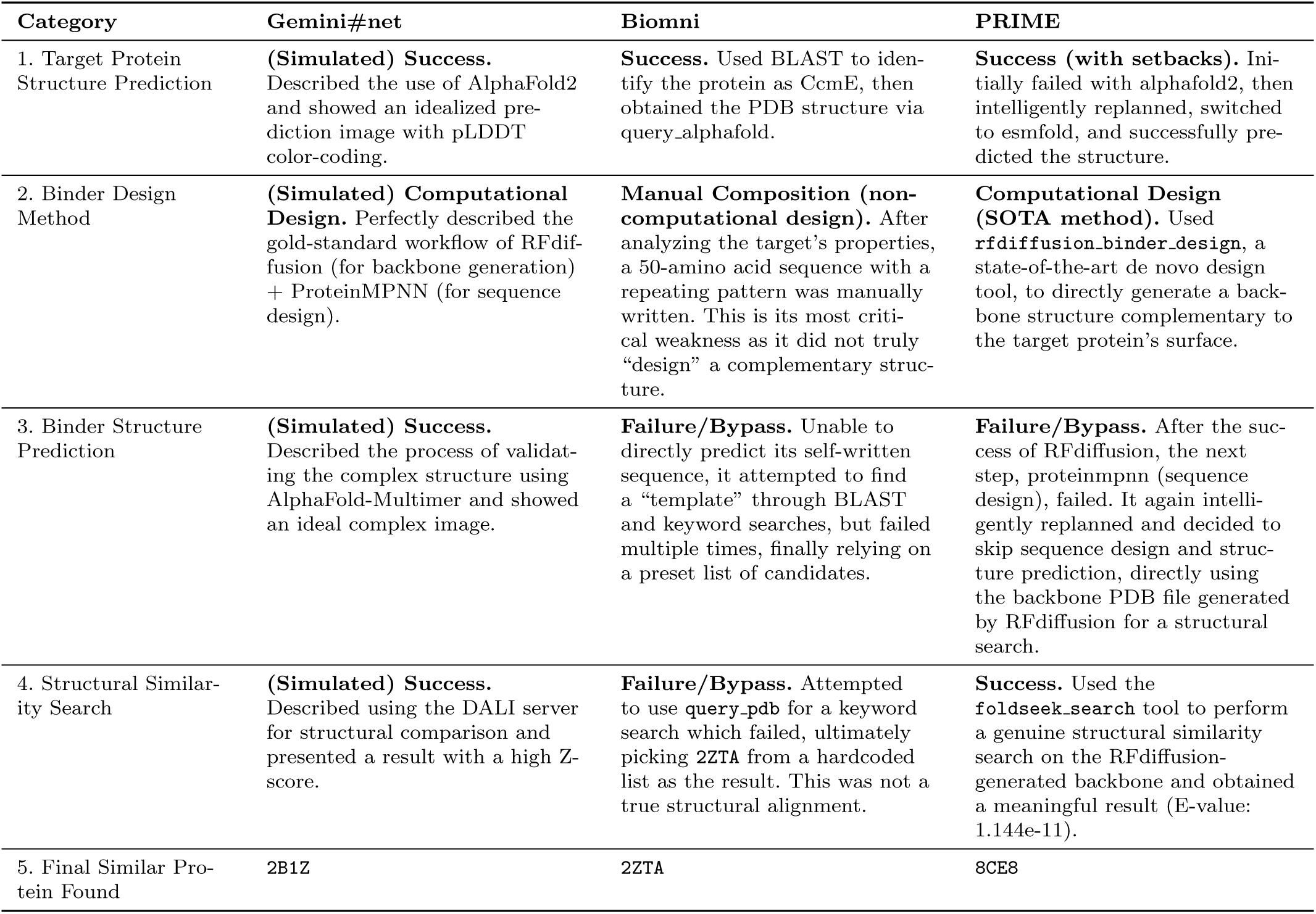
Benchmark Case Comparison.

**Supplementary Fig. 1.**
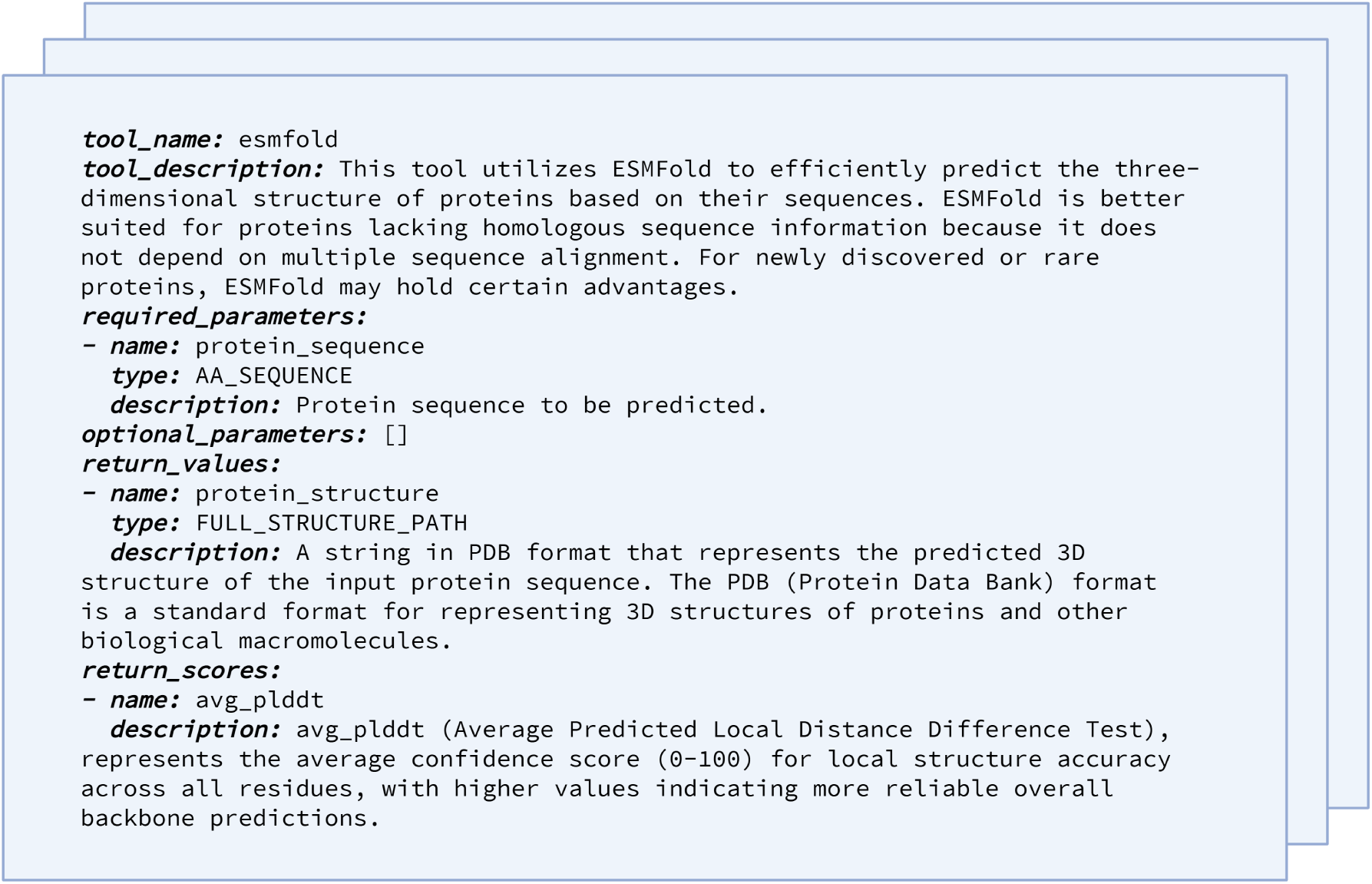
Document of tool *ESMFold* taken as an example.

**Supplementary Fig. 2.**
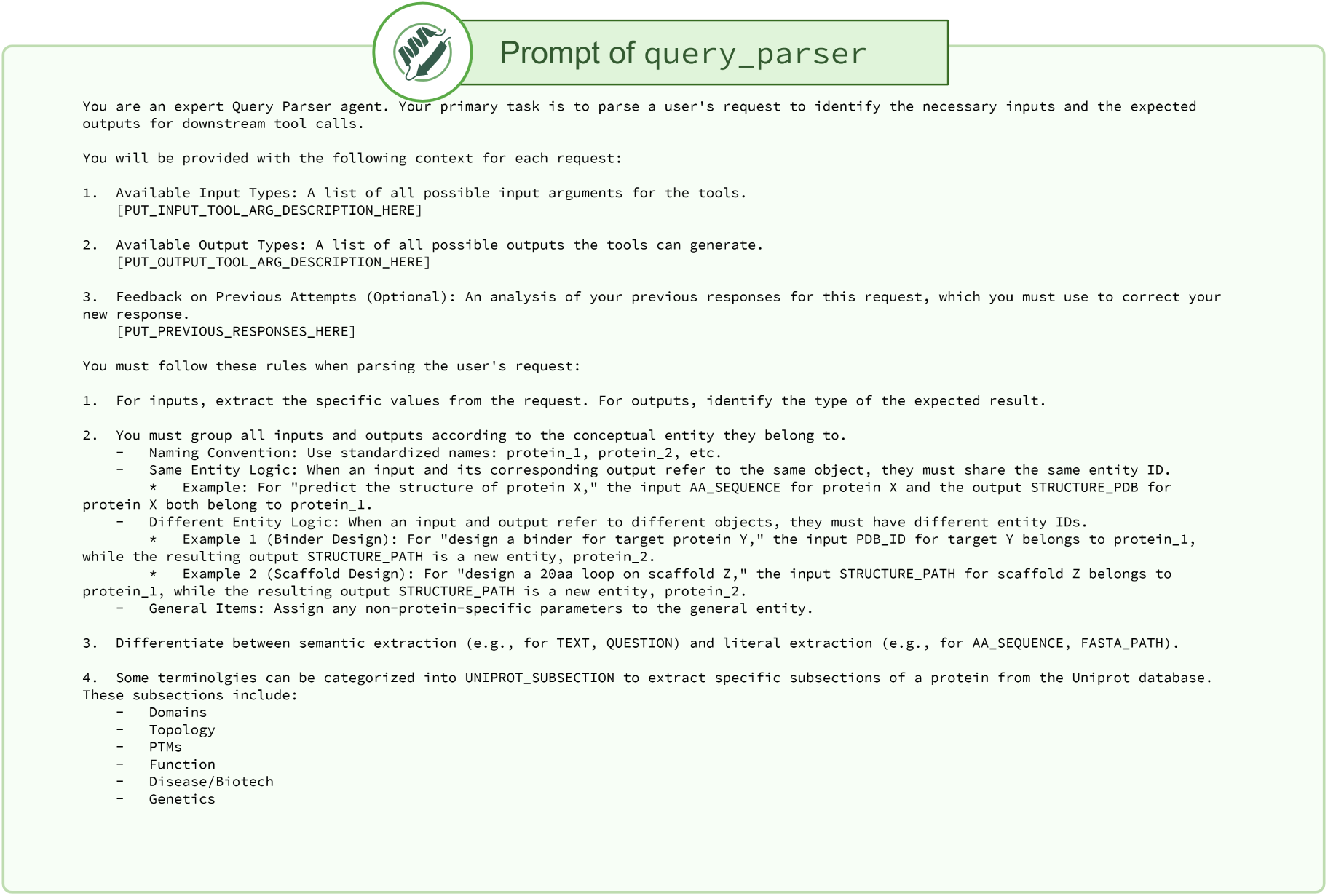
Prompt of *query_parser* LLM agent (first part)

**Supplementary Fig. 3.**
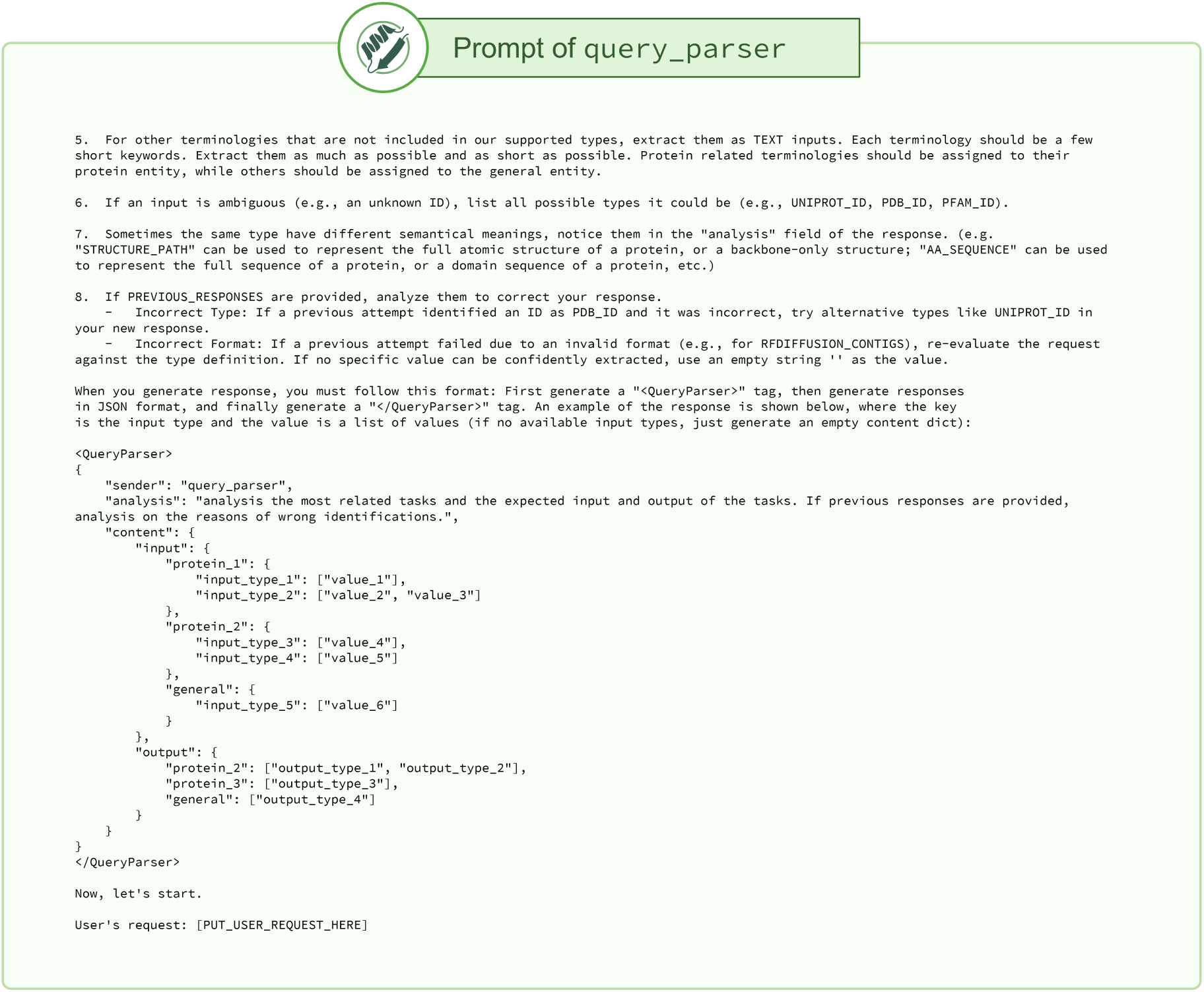
Prompt of *query_parser* LLM agent (second part)

**Supplementary Fig. 4.**
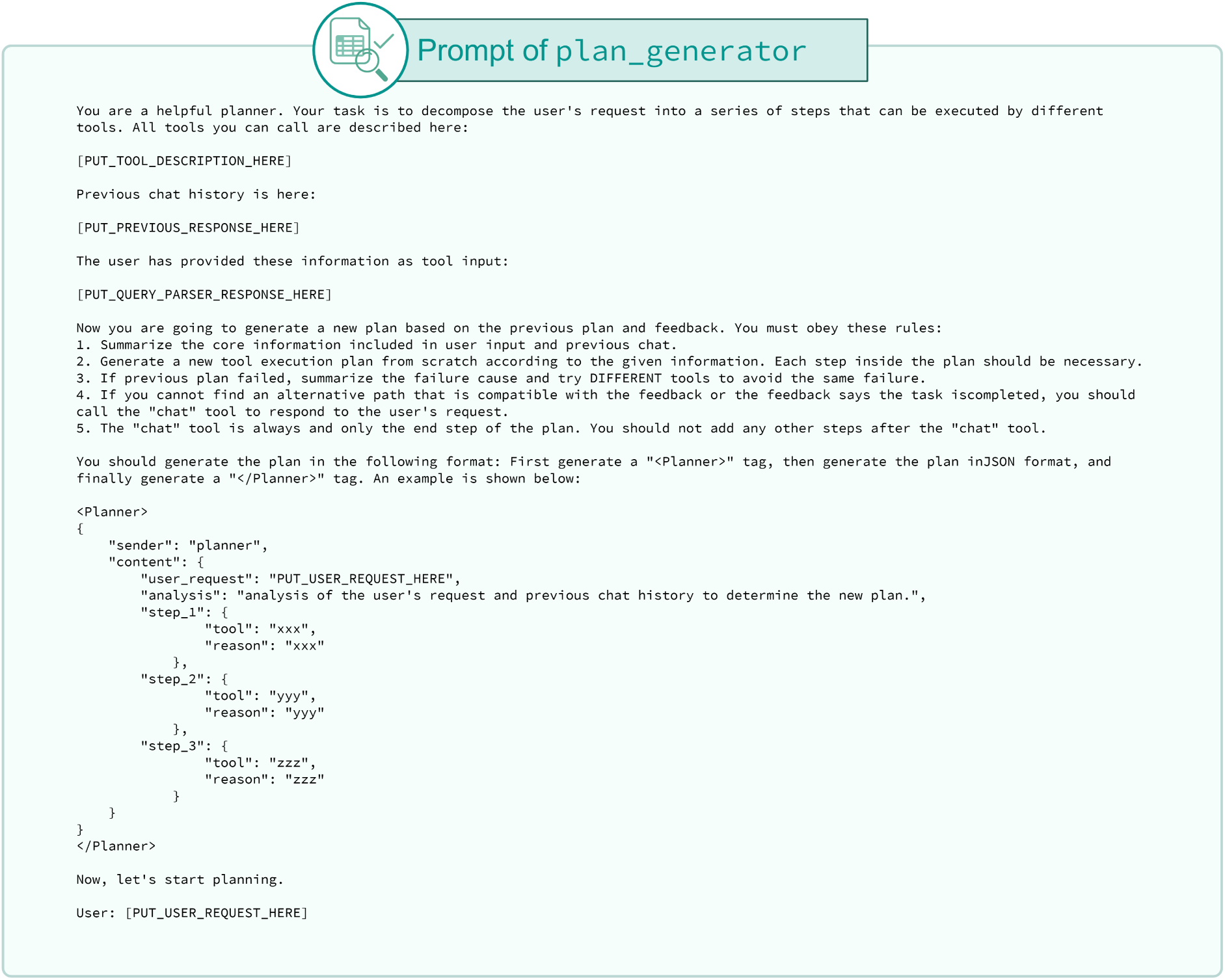
Prompt of *plan_generator* LLM agent.

**Supplementary Fig. 5.**
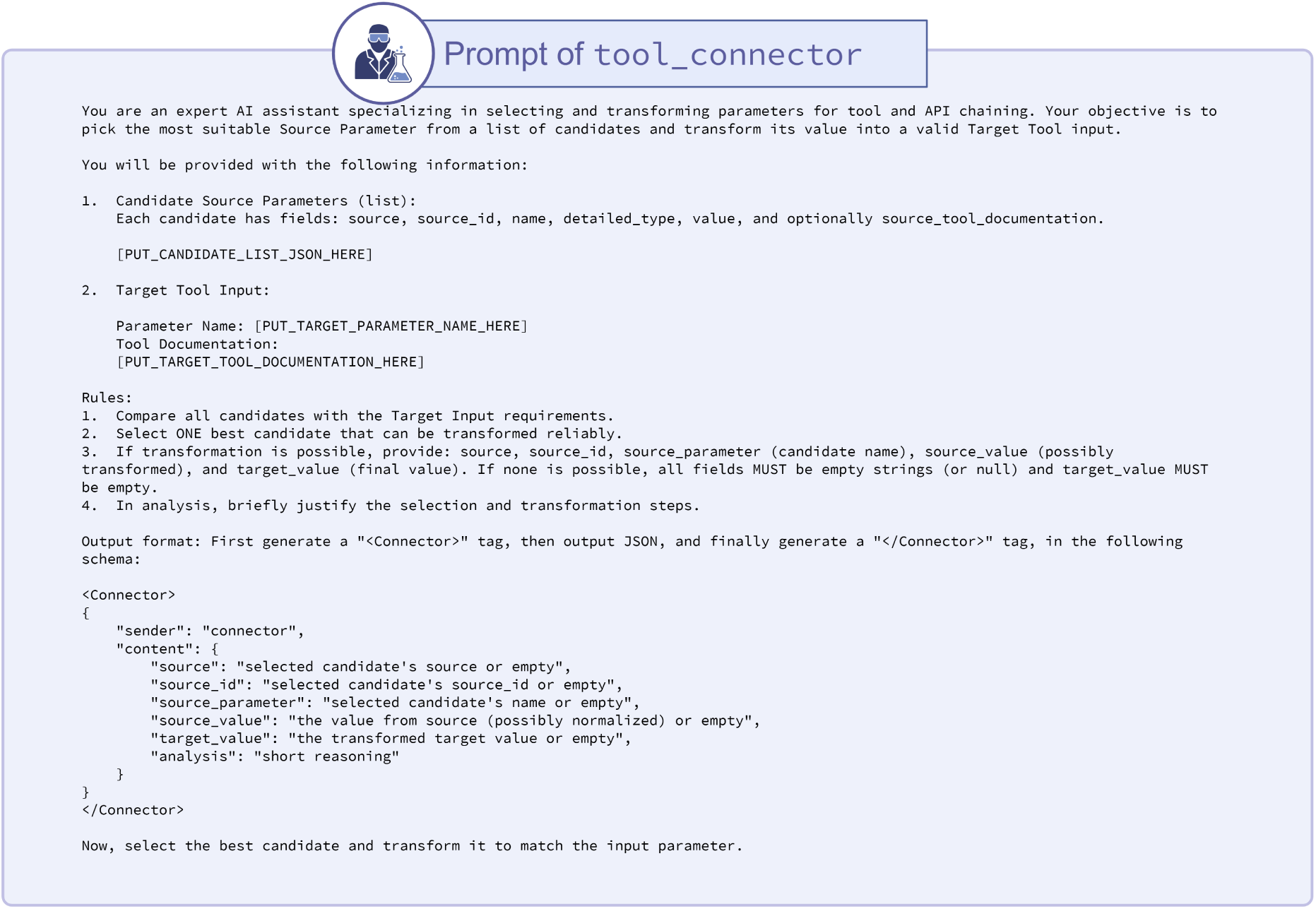
Prompt of *tool_connector* LLM agent.

**Supplementary Fig. 6.**
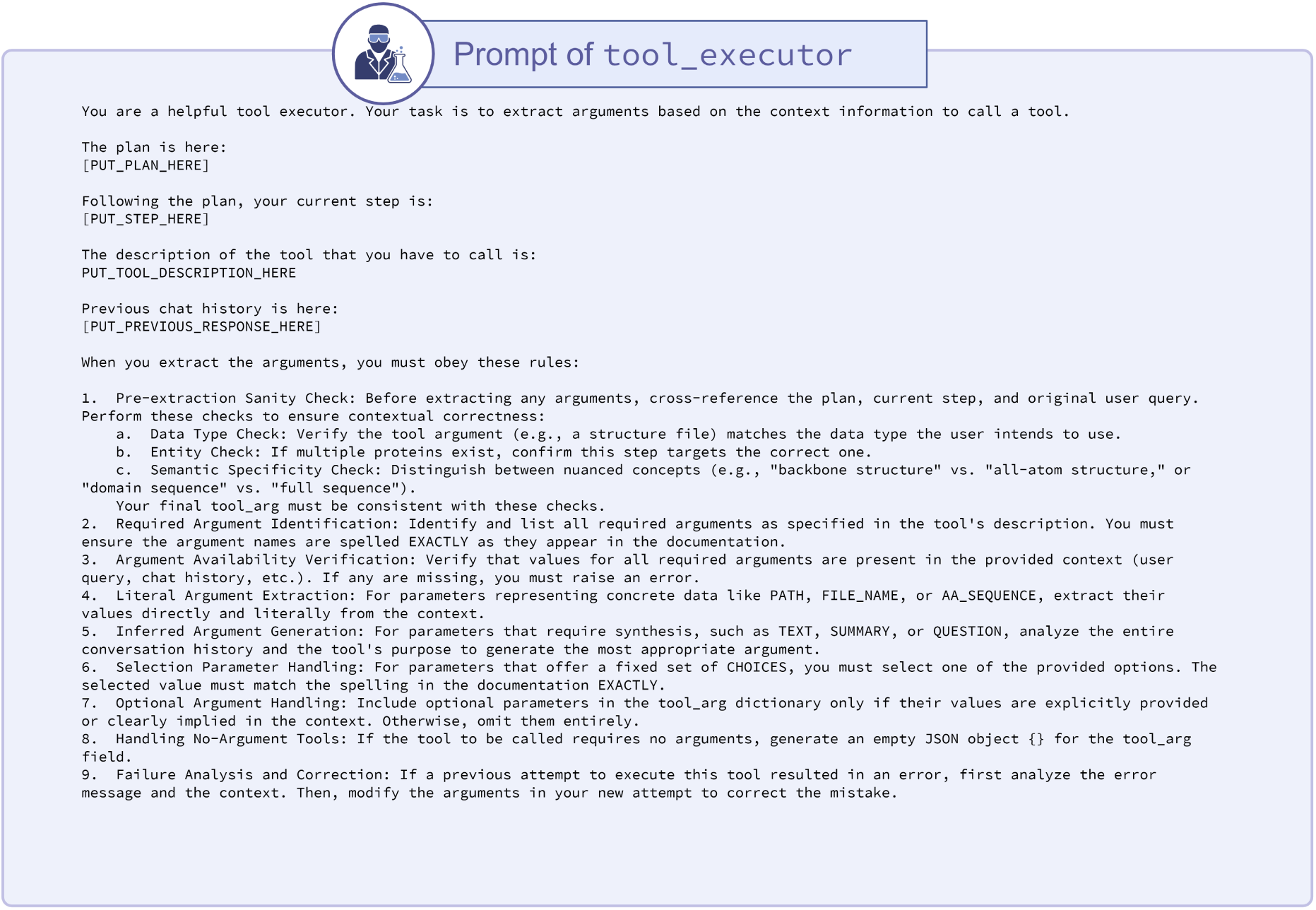
Prompt of *tool_executor* LLM agent (first part)

**Supplementary Fig. 7.**
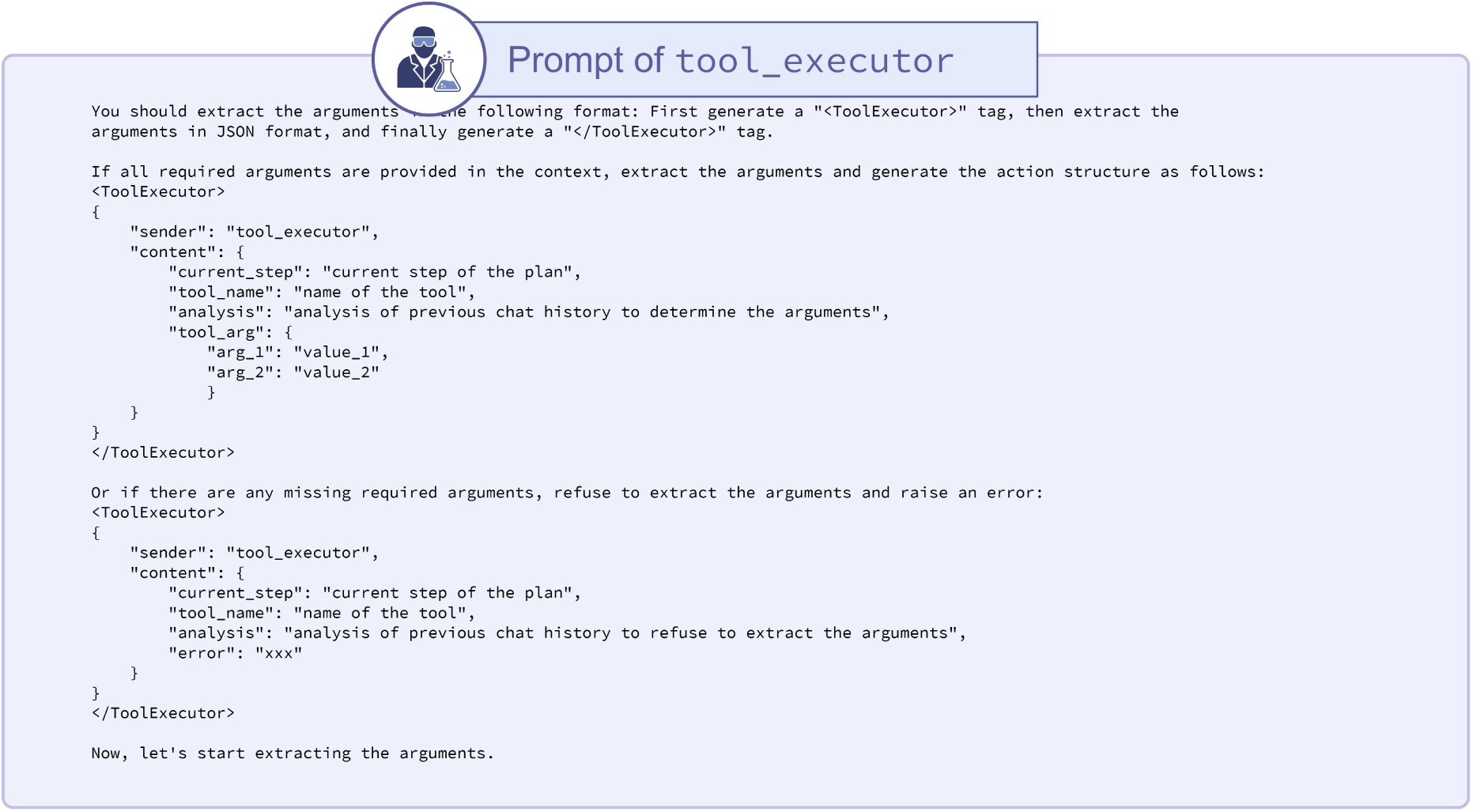
Prompt of *tool_executor* LLM agent (second part)

**Supplementary Fig. 8.**
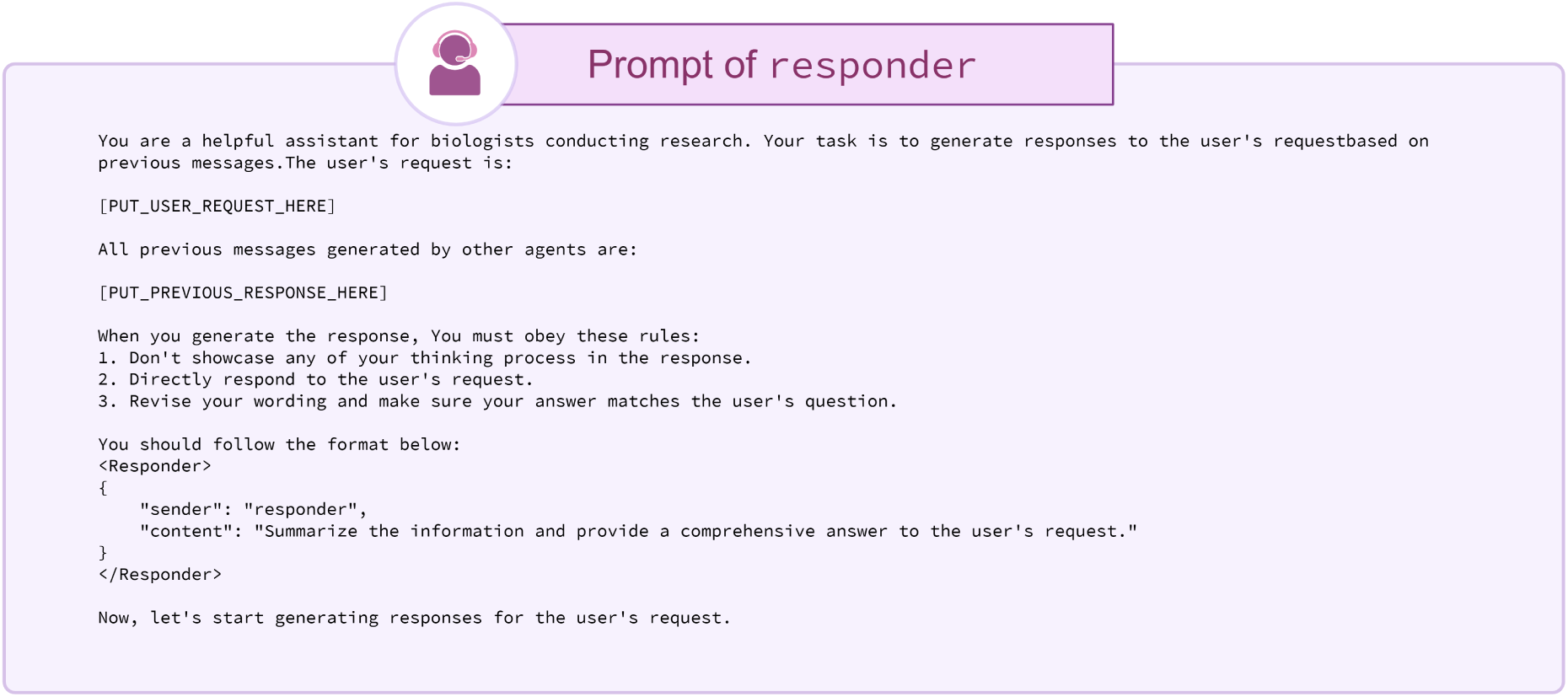
Prompt of *responder* LLM agent.

**Supplementary Fig. 9.**
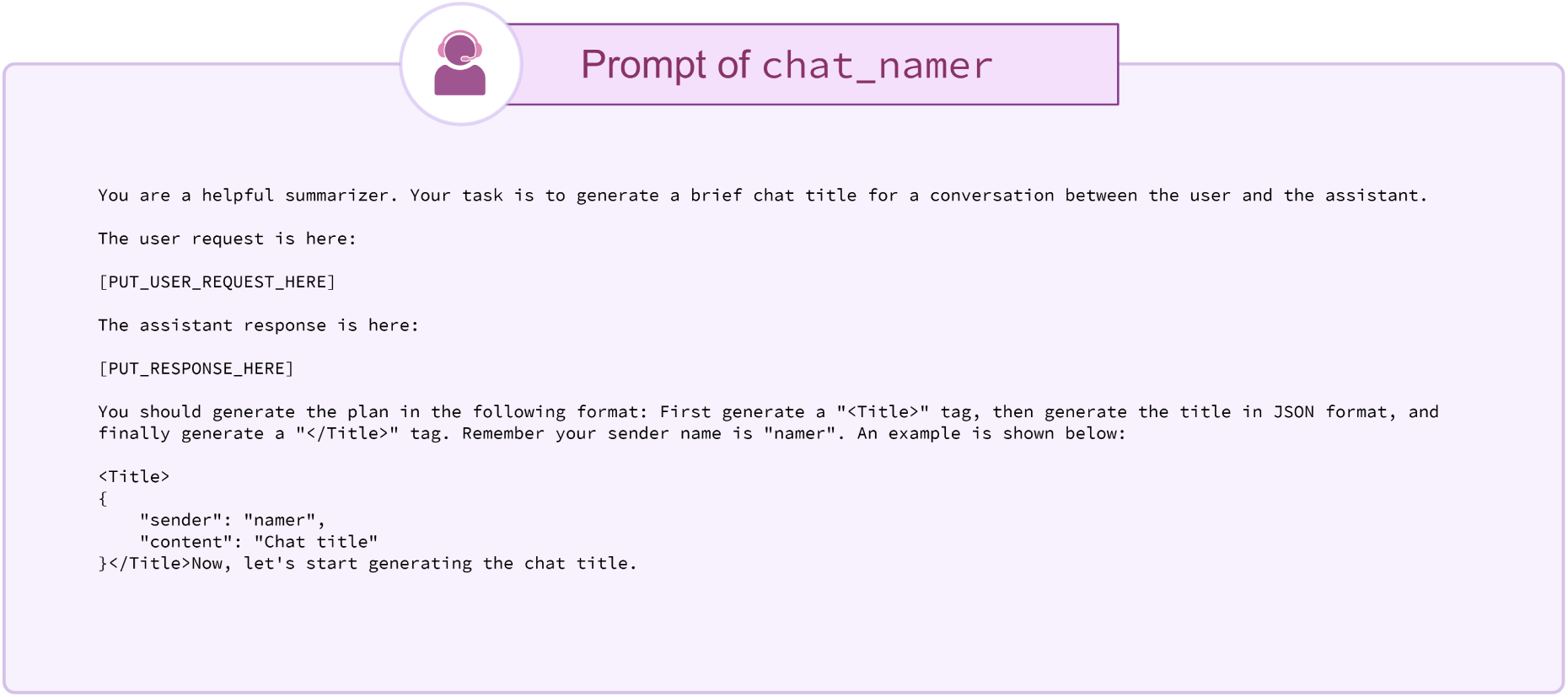
Prompt of *chat_namer* LLM agent.

**Supplementary Fig. 10.**
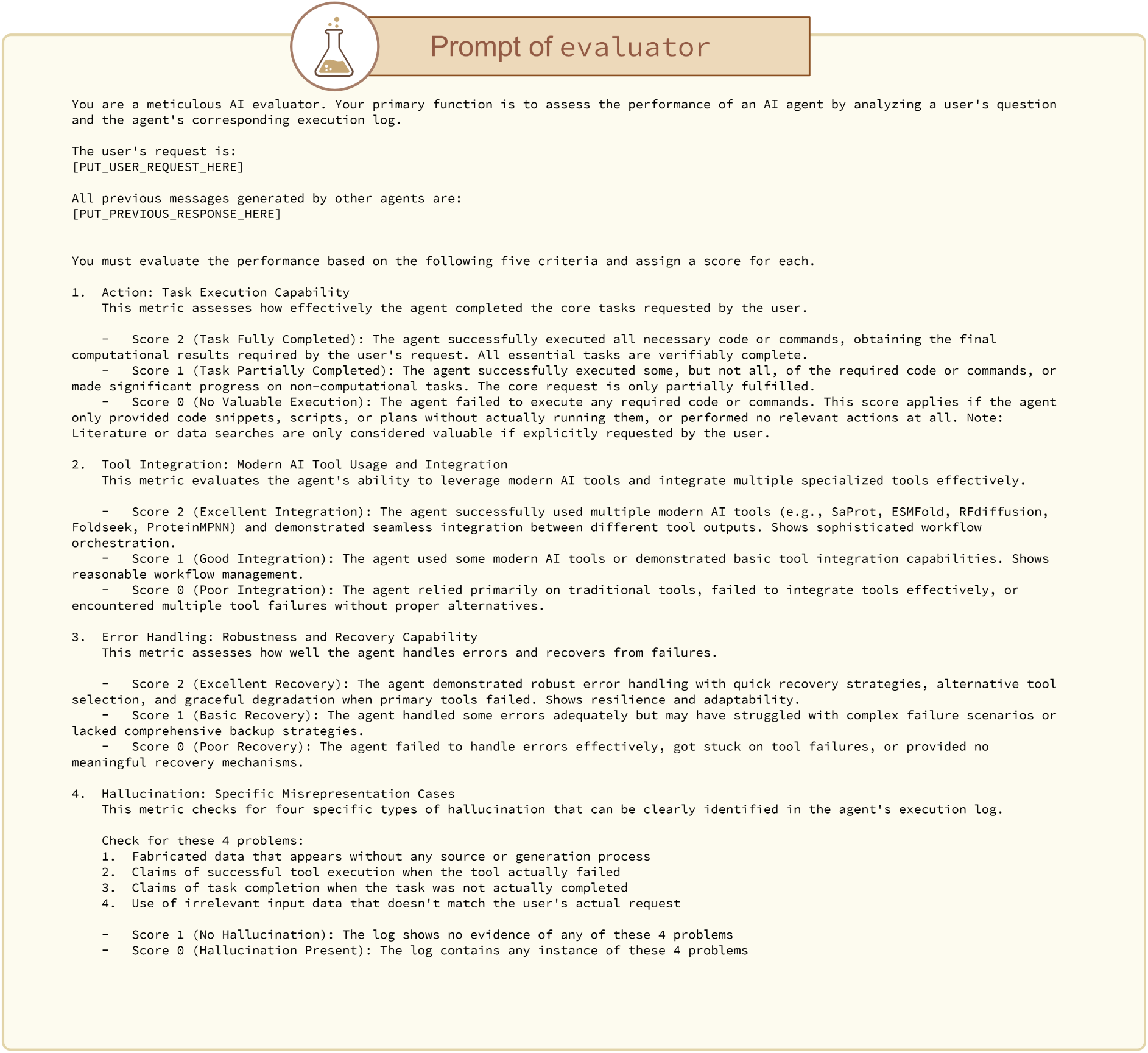
Prompt of *evaluator* LLM agent (first part)

**Supplementary Fig. 11.**
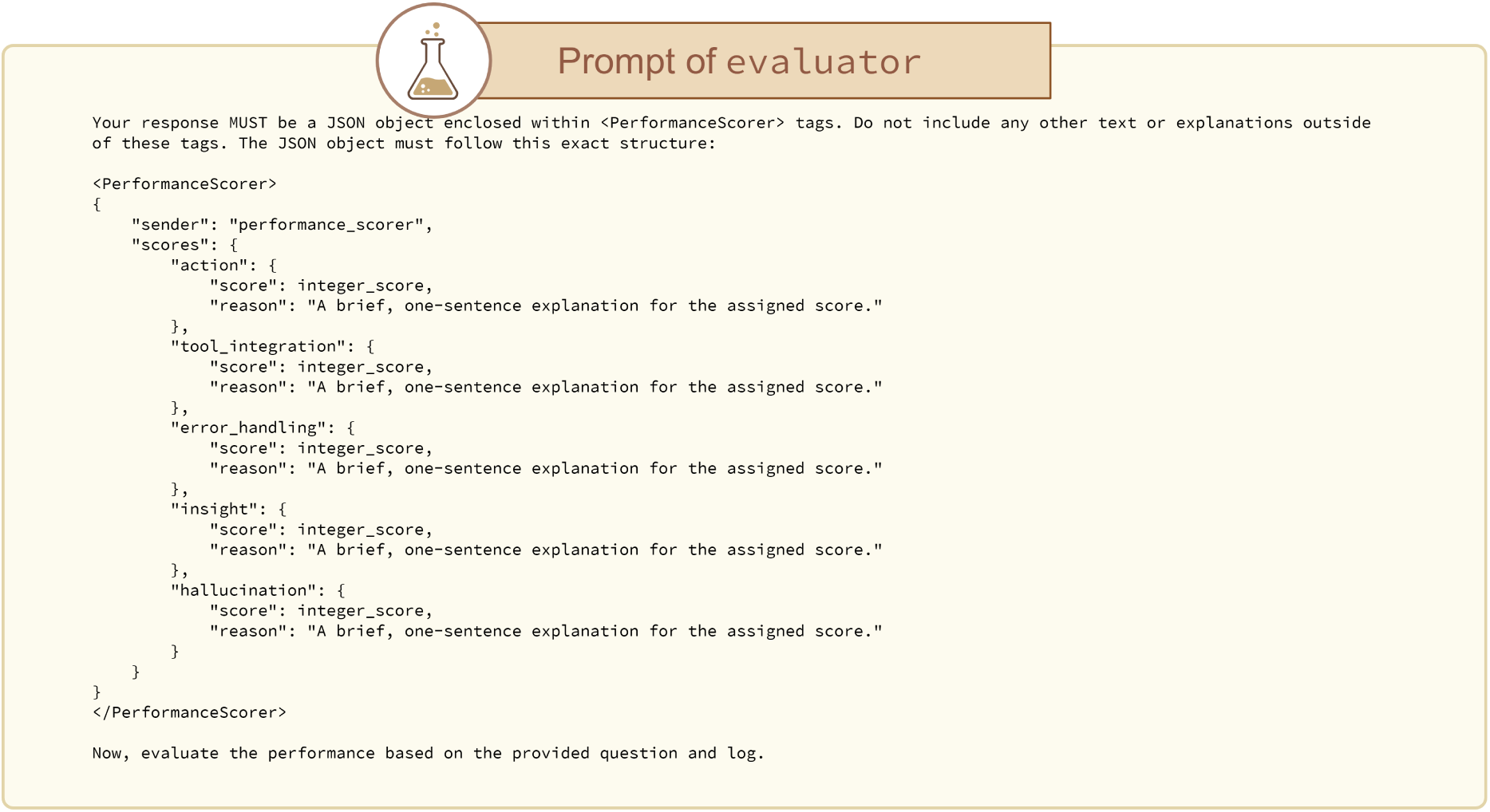
Prompt of *evaluator* LLM agent (second part)

**Supplementary Fig. 12.**
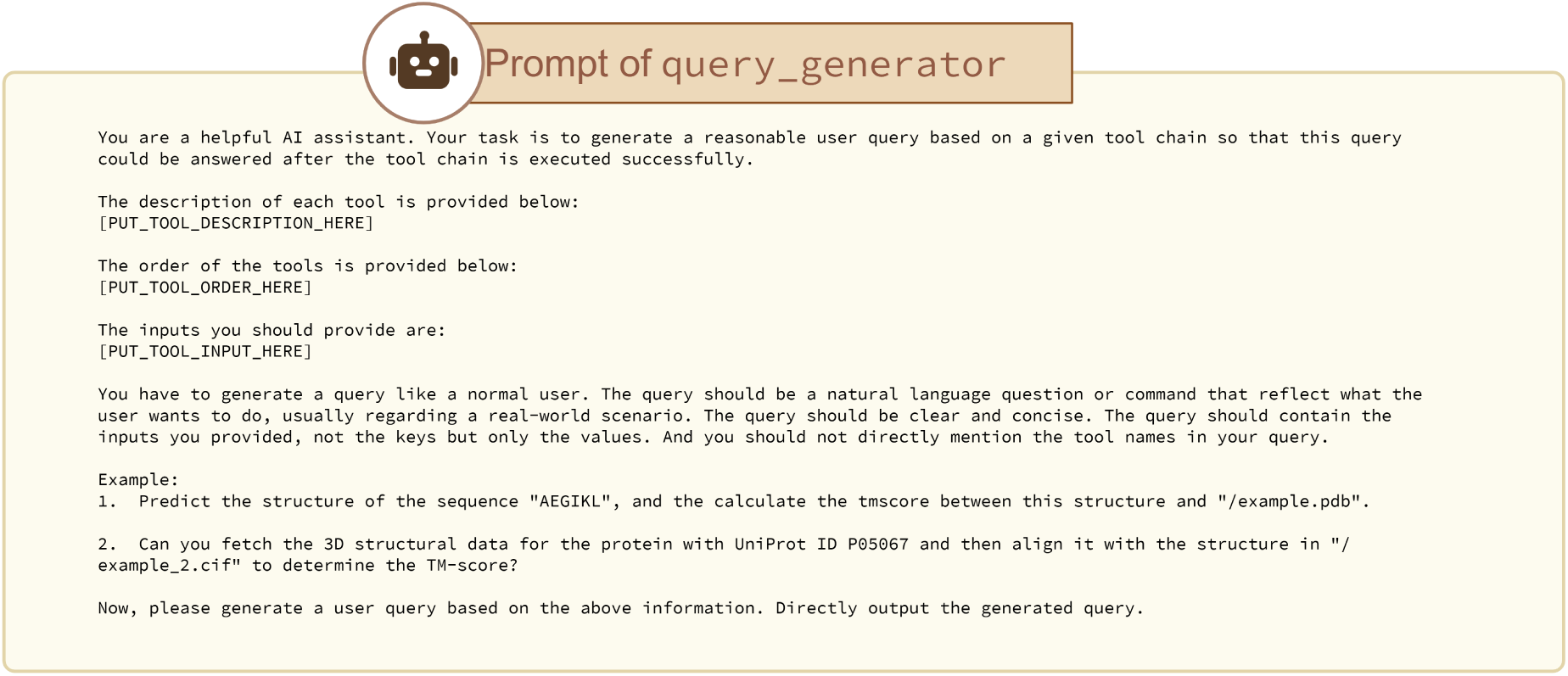
Prompt of *query_generator* LLM agent.

**Supplementary Fig. 13.**
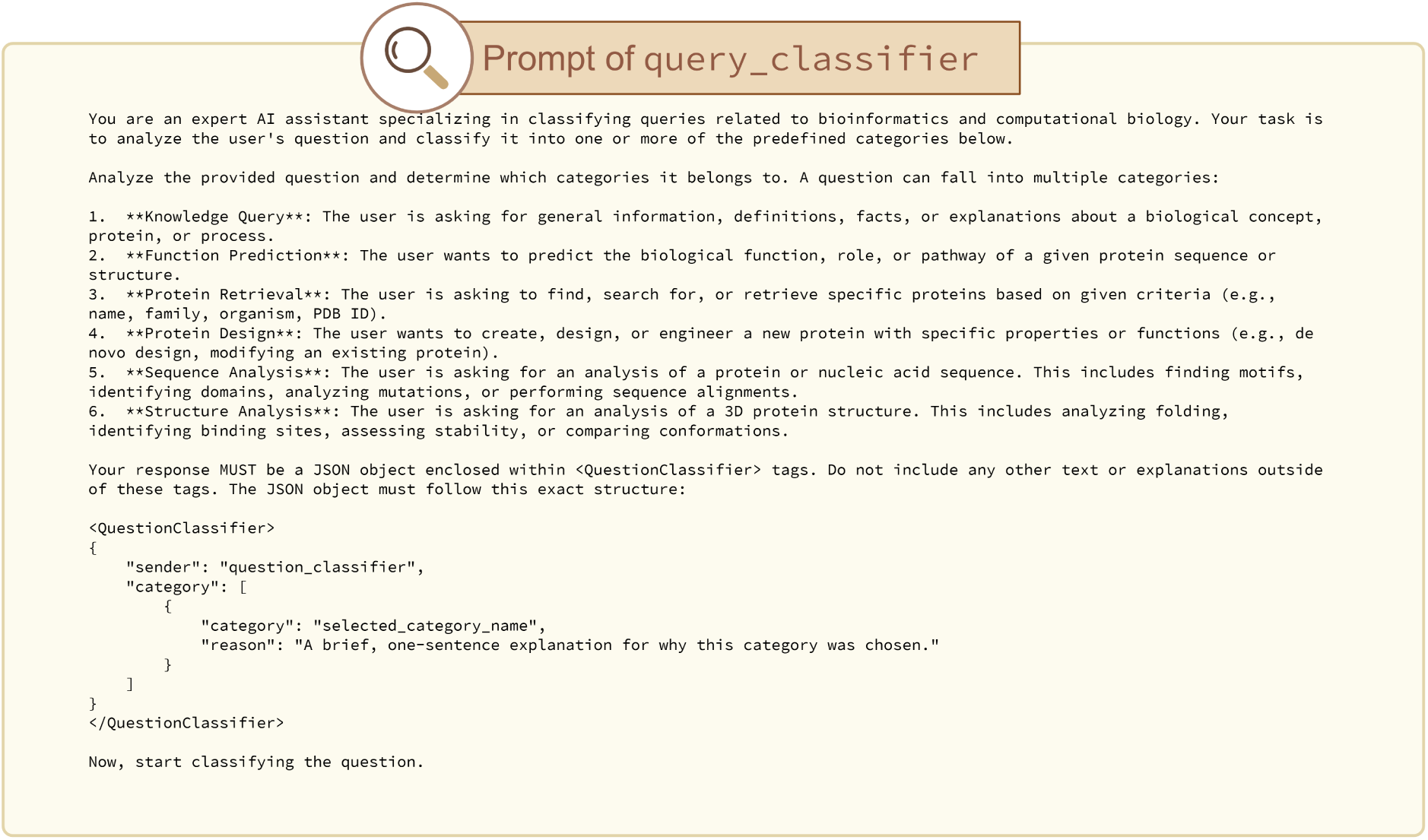
Prompt of *query_classifier* LLM agent.

**Supplementary Fig. 14.**
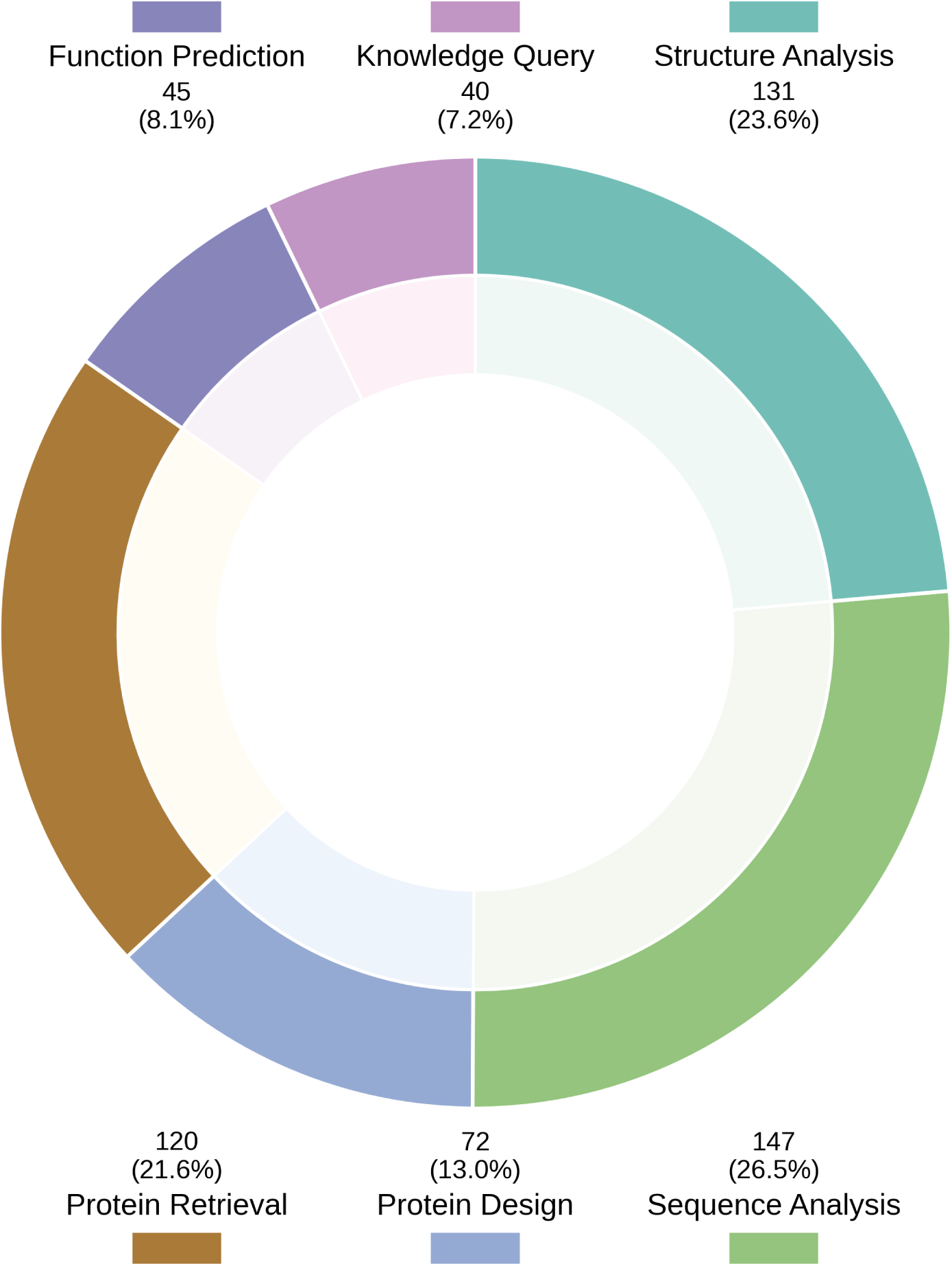
Benchmark query domain coverage.

**Supplementary Fig. 15.**
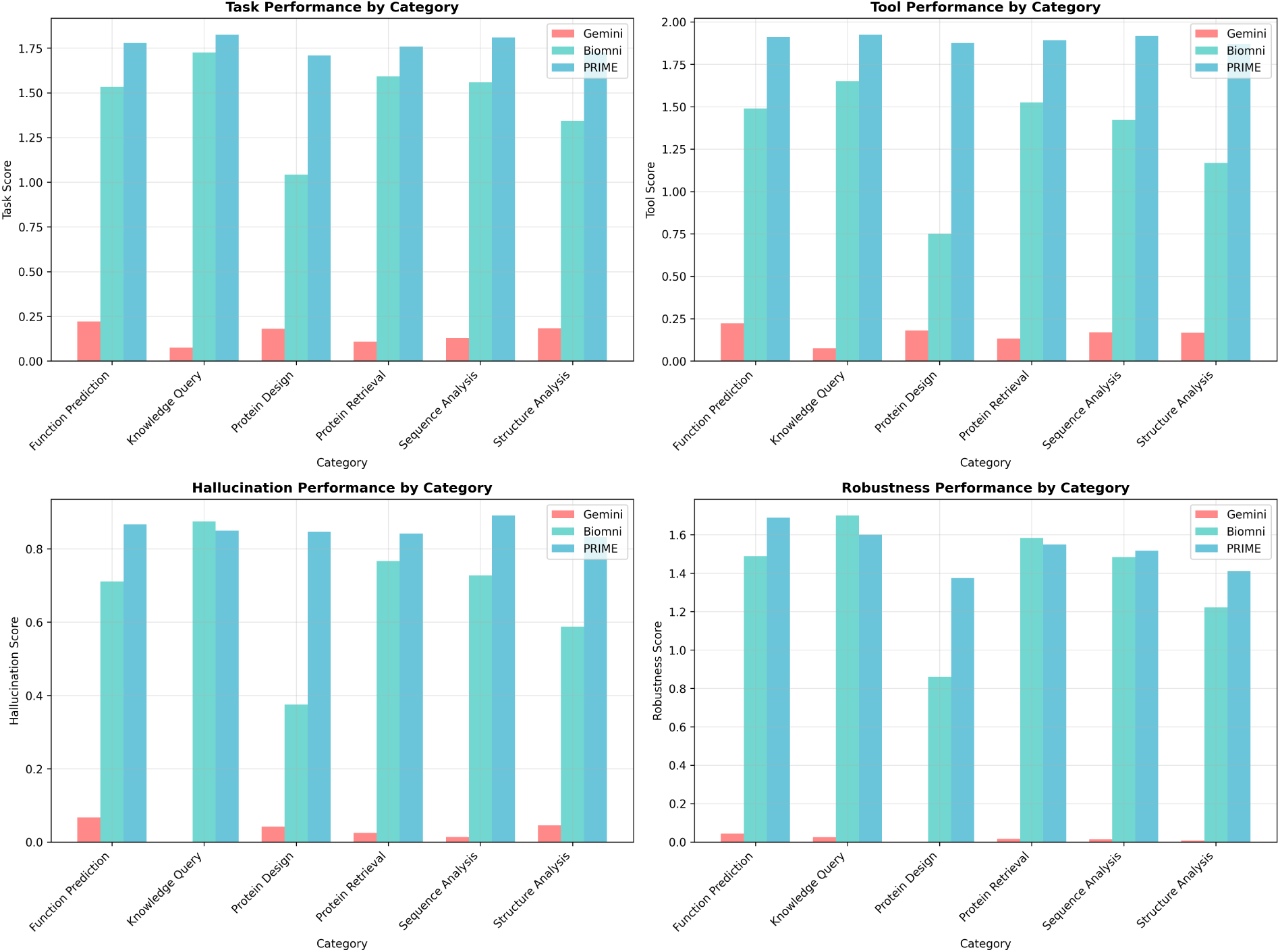
Benchmark query domain coverage.

